# Estimating dispersal using close kin dyads: The kindisperse R package

**DOI:** 10.1101/2021.07.28.454079

**Authors:** Moshe E Jasper, Ary A Hoffmann, Thomas L Schmidt

## Abstract

Investigating dispersal in animal populations can be difficult, particularly for taxa that are hard to directly observe such as those that are small or rare. A promising solution may come from new approaches that use genome-wide sequence data to detect close kin dyads and estimate dispersal parameters from the distribution of these dyads. These methods have so far only been applied to mosquito populations. However, they should have broad applicability to a range of taxa, although no assessment has yet been made on their performance under different dispersal conditions and study designs. Here we develop an R package and Shiny app, kindisperse, that can be used to estimate dispersal parameters from the spatial distribution of close kin. Kindisperse can handle study designs that target different life stages and allows for a range of dispersal kernel shapes and organismal life histories; we provide implementation examples for a vertebrate (*Antechinus*) and an invertebrate (*Aedes*). We use simulations run in kindisperse to compare the performance of two published close kin methodologies, showing that one method produces unbiased estimates whereas the other produces downward-biased estimates. We also use kindisperse simulations to investigate how study design affects dispersal estimates, and we provide guidelines for the size and shape of sample sites as well as the number of close kin needed for accurate estimates. kindisperse is easily adaptable for application to a variety of research contexts ranging from invasive pests to threatened species where non-invasive DNA sampling can be used to detect close kin.

## Introduction

Dispersal is a key ecological and evolutionary process that connects populations in space and time. Assessing dispersal in an invasive population can indicate future invasion risks (García-Berthou, 2007; Renault, Laparie, McCauley, & Bonte, 2018; With, 2004) and inform strategies for population suppression or manipulation (Petit et al., 2013; Stinner, Barfield, Stimac, & Dohse, 1983). For remnant populations requiring conservation, dispersal assessments can indicate isolation between remnants and the potential of populations to become locally extinct and inbred (Di Musciano et al., 2020; Driscoll et al., 2014; Trakhtenbrot, Nathan, Perry, & Richardson, 2005). Depending on the aims and framework of the investigation, dispersal can be treated as a component of an individual’s life history (Howard, 1960) or as an intergenerational process (Wright, 1931). Intergenerational investigations typically consider the parent-offspring dispersal parameter σ, which determines how local genetic variation is spatially structured (Wright, 1946) and can influence rates of coalescence (Wilkins & Wakeley, 2002) and relative rates of gene flow and genetic drift (Barton & Wilson, 1995).

Various methods have been developed for observing and parameterising dispersal, each having advantages when applied to different systems. For large animals, mark-release-recapture (MRR) methods are frequently used for assessing individual movement (Royle & Young, 2008) and ‘recaptures’ can be effectively performed via camera traps for ease of observation (Silver et al., 2004). Assessing dispersal of invertebrates and other small, abundant, or short-lived animals with MRR methods is typically performed using dyes, paint or chemical tags (Hagler & Jackson, 2001; Hagler, Naranjo, Machtley, & Blackmer, 2014), but these methods can suffer from limitations including: (i) they require manipulation of organisms, which may change behaviour; (ii) when conducted across a sufficiently large area to be informative they can be very labour-intensive; and (iii) they typically do not provide estimates of true intergenerational dispersal, which is measured life stage to life stage (e.g. egg to egg). Dispersal distances estimated from tagged individuals are thus not always readily interpretable within established intergenerational analytical frameworks such as Wright’s neighbourhood size (Wright, 1946).

For assessing intergenerational dispersal, methods using molecular markers are commonly applied. These include methods using pairwise genetic similarities among individuals to estimate N_W_, from which σ can be inferred when local density data are available (Broquet et al., 2006).

Genetic relatedness can also be treated as categorical, in which pairwise genetic similarity is used to assign dyads to specific categories of close kin. Parentage studies using these methods have investigated pollen and seed dispersal in plants (Chybicki & Oleksa, 2018), which can work well for assessing dispersal over long distances (Smouse & Sork, 2004). Dispersal inferences from close kin have been successfully extended to animal taxa with appropriate study designs (Burland, Barratt, Nichols, & Racey, 2001; Schmidt et al., 2021).

Recent genetic investigations of dispersal have used genome-wide sequence data to assign dyads to kinship categories across multiple orders of kinship, which can currently be inferred to the 3^rd^ order (e.g. first cousin). These methods build on close kin mark-recapture (CKMR) methods used primarily for demographic analysis of marine populations (Bravington, Skaug, & Anderson, 2016; Waples & Feutry, 2021), in which genetic relationships between close kin are deemed to both ‘mark’ and ‘recapture’. Unlike parentage analysis, CKMR methods do not require any samples from the parental generation, making these methods appealing for assessing dispersal in short-lived taxa or those without overlapping generations (Filipović et al., 2020; Jasper, Schmidt, Ahmad, Sinkins, & Hoffmann, 2019). Genetic approaches like CKMR can take less experimental effort than MRR methods and the genetic data can be used for additional analyses beyond dispersal.

Two CKMR methods have been developed for estimating dispersal parameters. The first of these methods (which we refer to here as the Jasper et al. method) used genome-wide SNPs to detect close kin dyads in a population of *Aedes aegypti* mosquitoes sampled from urban high-rise apartments in Malaysia (Jasper et al., 2019). From 162 *Ae. aegypti* that were sequenced, 98 close kin dyads were detected: 13 full-sibs, 34 half-sibs, and 51 first cousins. Using the spatial distribution of full and half-sibs, the Jasper et al. method decomposes the first cousin dispersal kernel to estimate σ, which was estimated in this population at 46m (95% C.I. 16 – 66m). A second method (which we refer to here as the Filipović et al. method) used close kin dyads to estimate σ as 45.2m (95% C.I. 39.7 – 51.3m) in *Ae. aegypti* sampled from urban high-rise apartments in Singapore (Filipović et al., 2020). The Filipović et al. method first adjusts each kin category by a fixed factor representing inferred acts of movement, then combines them for the final intergenerational estimate. While these two methods report similar estimates of σ in these specific cases, the Jasper et al. and Filipović et al. methods differ in their theoretical approaches and sampling designs. The performance of these methods has yet to be compared with the same data set.

In this paper, we introduce kindisperse, a set of tools for the simulation and parameterisation of intergenerational dispersal from the distribution of close kin dyads. When applied to empirical data, kindisperse can estimate intergenerational dispersal parameters from a set of georeferenced dyads of known kinship category (Table 1), which may be determined genetically or by other means. Simulations run in kindisperse can test the impact of study sampling design on dispersal parameterisation and thus help optimise sampling protocols. kindisperse is implemented as a combined R package and shiny app. Our purpose here is to provide a user-friendly tool implementing these dispersal methods for analysing experimental data and for designing dispersal studies. These tools are designed to allow researchers and practitioners to adopt and effectively implement CKMR dispersal estimation methods.

**Table 1.** Abbreviations for kinship categories used in this paper.

|  | **FS Phase** | **HS Phase** | **PO Phase** |
| --- | --- | --- | --- |
| **Gen 1** | FS (full-sib) | HS (half-sib) | PO (parent-offspring) |
| **Gen 2** | AV (avuncular) | HAV (half avuncular) | GG (grandparent-grandchild) |
|  | 1C (first cousins) | H1C (half first cousins) |  |
| **Gen 3** | GAV (grand avuncular) | <u>HGAV</u> (half grand avuncular) | GGG (great grandparent-great grandchild) |
|  | 1C1 (first cousin once removed) | H1C1 (half first cousin once removed) |  |
|  | 2C (second cousin) | H2C (half second cousin) |  |

As an illustration of kindisperse, we use simulations implemented within kindisperse to compare the performance of the Jasper et al. and Filipović et al. methods (Filipović et al., 2020; Jasper et al., 2019). We find that only the Jasper et al. method (further developed within this package) accurately estimates the underlying dispersal parameters and we show that this method is applicable to a range of dispersal kernel shapes. Finally, we use kindisperse to develop a series of best practice recommendations for designing studies of dispersal built on the Jasper et al. method. The approach should be applicable to any organism where related individuals can be sampled across space.

## Materials and Methods

The following sections describe how dispersal kernels operate and how dispersal can be parameterised from close kin dyads. Many of the examples will make specific reference to the biology of *Aedes aegypti* (yellow fever mosquito), as this mosquito has been the focus of recent kin-based dispersal investigations (Filipović et al., 2020; Jasper et al., 2019). However, these ideas are directly relevant (or can be adapted) to other sexually-reproducing organisms, and the kindisperse package is designed for taxa with a range of dispersal characteristics and life histories (see Supplementary Text 3 section 4.4 for application to *Antechinus*). All close kin abbreviations are listed in Table 1.

### Intergenerational dispersal kernels

#### (i) Dispersal location kernels

A dispersal location kernel, as opposed to a dispersal distance kernel (Nathan, Klein, Robledo-Arnuncio, & Revilla, 2012), is a probability density function describing the distribution of the positions of dispersed individuals relative to an origin (Figure 1a & b). The scale of dispersal is described by the variance component, σ^2^, of whatever probability density function is used to represent the kernel. The most fundamental intergenerational dispersal kernel, the lifespan or parent-offspring (PO) kernel, reflects all dispersal and breeding processes connecting one life stage of a parent to the same life stage of its offspring (e.g. egg_(t-1)_ to egg_(t)_). For other non-PO kinship categories, the positions of dispersed individuals may result from different sets of dispersal events, some of which span less than an entire reproductive cycle and some of which span multiple cycles. When σ appears in the literature, it is typically referring to the PO kernel (e.g. Wright (1946)), but here we use the specific notation σ_PO_ to distinguish it from other dispersal location kernels.

**Figure 1.**
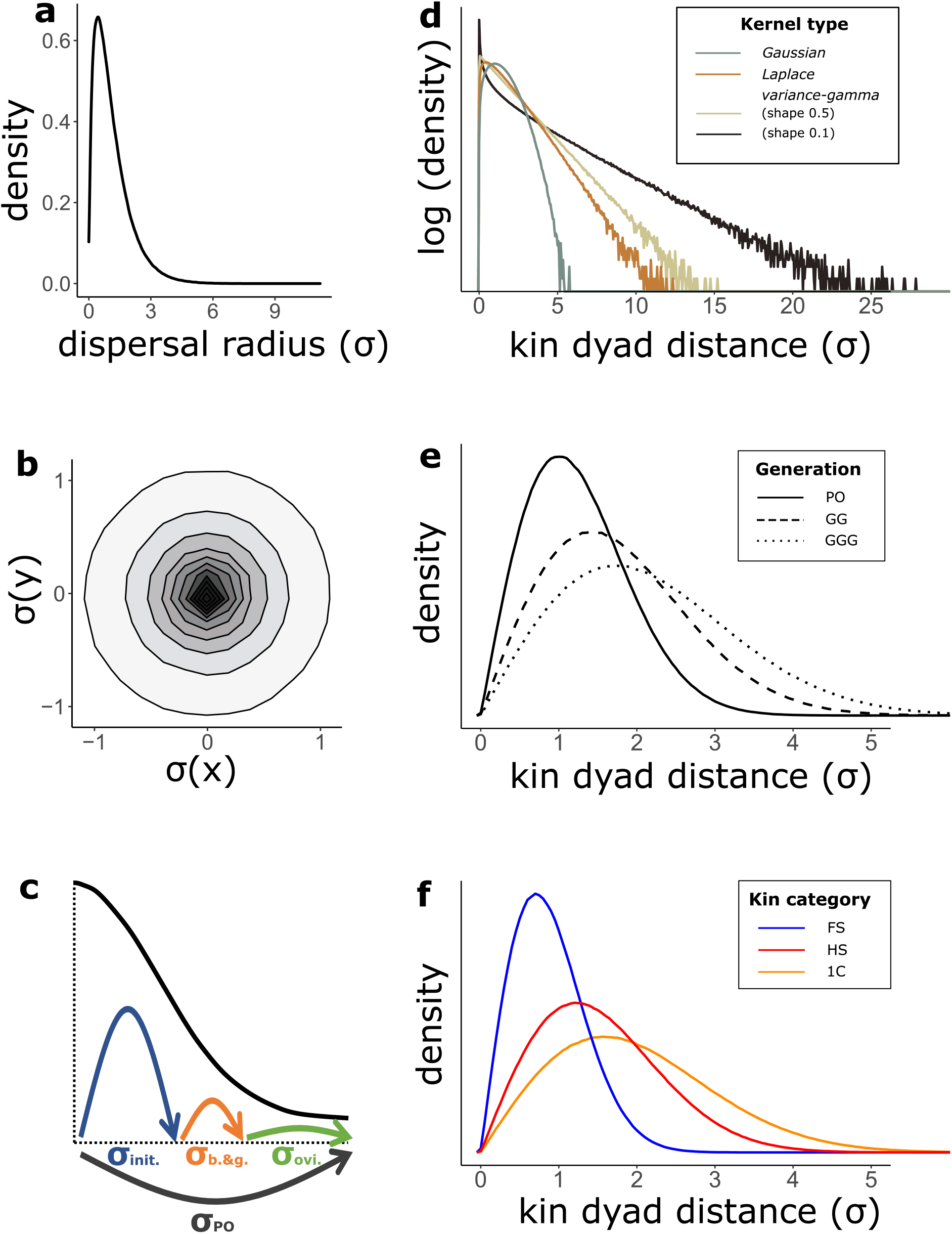
Properties of dispersal kernels used in kindisperse. **a.** Dispersal distance kernel, defined one-dimensionally with respect to the radius from the origin point for dispersal, in contrast to **b.** Dispersal location kernel, defined two-dimensionally with respect to X and Y coordinates around the dispersal origin. Shading represents density. The location kernel preserves information about the relative distances between multiple dispersed individuals, as it captures both distance and angle of dispersal. **c.** Simple vs composite representation of dispersal across a lifespan. Both the entire parent-offspring dispersal event (grey arrow) or the component lifespan dispersal events that make it up (coloured arrows) produce the same dispersal distribution (black line). Composite σ parameters are here taken from the mosquito lifespan. **d-f.** Impact of kernel type, generation, and kin category parameters on outcomes of kinship dispersal simulations (shown as dispersal distance kernels). All simulations set σ_po_ to 1; simulations involving smaller dispersal components divided the po variance equally between each component, i.e. four values of σ = 0.5. **d.** Distribution used to model dispersal kernel (Gaussian, Laplace, and two variance-gamma kernels) – log scale used. **e.**Number of generations modelled (po = 1; gg = 2, ggg = 3). **f.** Modelling of typical kin categories.

#### (ii) Composing and decomposing dispersal kernels

Consider a dyad of immature (egg, larval or pupal) first cousin (1c) mosquitoes, each of which will have one parent that is part of a full-sib (FS) dyad. Each individual’s location results from a separate draw from the underlying PO kernel, in addition to draws from the ovipositional kernel of their shared female grandparent. A simple PO kernel is insufficient to describe these distributions; composite dispersal kernels can instead be constructed using additive properties of variance (Figure 1c). Intergenerational dispersal kernels are constructed by two such extensions: (1) the dispersal of one individual is extended to dispersal across multiple individuals within a pedigree via variance addition; (2) the dispersal of one individual is split into multiple phases across its lifespan by variance subtraction (Figure 1c).

### Modelling intergenerational dispersal with close kin dyads

#### (i) Full-sib, half-sib, and first cousin kernels

To demonstrate how dispersal is inferred from close kin dyads, we consider a population of *Ae. aegypti* that has been sampled within the temporal range of a single gonotrophic cycle (~1-2 weeks) (Jasper et al., 2019; Schmidt, Filipović, Hoffmann, & Rašić, 2018). A sufficiently large sample should contain FS, HS, and 1C, as well as less closely related and completely unrelated dyads.

For a dyad of immature FS, any geographical separation found between them will be due either to their dispersal as immatures (typically negligible) or the mother’s ‘skip oviposition’ behaviour (Colton, Chadee, & Severson, 2003), in which the female will lay a series of eggs across different oviposition sites. The distance between these fs will reflect the movement of their mother during a single gonotrophic cycle of oviposition (σ_ovi_).

In *Ae. aegypti*, fs share both parents, while hs share one parent (the father, given that females normally mate once). Thus, for a hs dyad, the distance will represent the movement of the father between matings (σ_breed_) as well as the movement of the mother searching for bloodmeals (σ_grav_) and during oviposition (σ_ovi_). For our current modelling effort, we combine the hs categories of σ_breed_ and σ_grav_, though these can be considered as separate processes if required (e.g. for sex-biased dispersal).

A 1c dyad will share a set of grandparents. The distance separating a 1C dyad represents an entire lifespan of dispersal (σ^2^_PO_) as well as movement of their mother during oviposition (σ^2^_ovi_), so that the dispersal location kernels of 1C = FS + 2σ^2^_PO_ are relevant. Other close kin relationships can be modelled additively following Table 1.

#### (ii) Composite kernels

Table 1 describes how each category of close kin has one of three ‘phases’: the full-sib (FS), the half-sib (HS), and the parent-offspring (PO) phases. The spatial distribution of dyads from a given close kin category has a variance composited from the phase variance (σ^2^_FS_, σ^2^_HS_, or σ^2^_PO_) plus an additive component representing the number of pedigree generations and the life stage at which the kin were sampled (Table 2). Phase variances are as follows: σ^2^_FS_ = 2σ^2^_ovi_ (i.e. two draws from the oviposition kernel); σ^2^_HS_ = 2(σ^2^_breed & grav_ + σ^2^_ovi_); and σ^2^_PO_ = σ^2^_init_ +σ^2^_breed & grav_ + σ^2^_ovi_, where σ^2^_init_ represents any movement taking place before the initial mating (including dispersal of immatures).

**Table 2.**
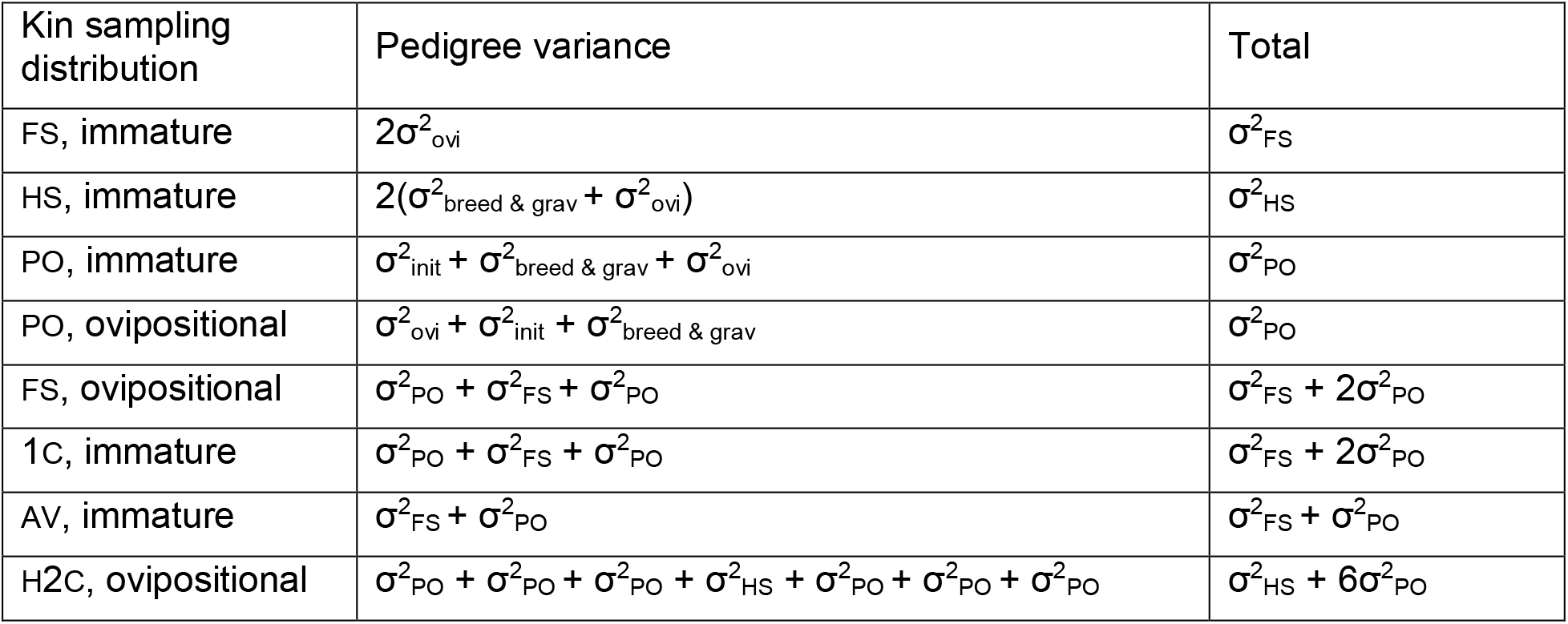
Example of kinship distributions given by additive variances. ‘Immature’ distributions represent the sampling locations of immature (egg, larval, pupal) mosquito life stages, such as (in the case of mosquitoes) those collected from ovitraps. ‘Ovipositional’ distributions represent the sampling locations of adult egg-laying females, such as (in the case of mosquitoes) those collected from gravitraps.

| Kin sampling distribution | Pedigree variance | Total |
| --- | --- | --- |
| FS, immature | $2\sigma_{\text{ovi}}^2$ | $\sigma_{\text{FS}}^2$ |
| HS, immature | $2(\sigma_{\text{breed \& grav}}^2 + \sigma_{\text{ovi}}^2)$ | $\sigma_{\text{HS}}^2$ |
| PO, immature | $\sigma_{\text{init}}^2 + \sigma_{\text{breed \& grav}}^2 + \sigma_{\text{ovi}}^2$ | $\sigma_{\text{PO}}^2$ |
| PO, ovipositional | $\sigma_{\text{ovi}}^2 + \sigma_{\text{init}}^2 + \sigma_{\text{breed \& grav}}^2$ | $\sigma_{\text{PO}}^2$ |
| FS, ovipositional | $\sigma_{\text{PO}}^2 + \sigma_{\text{FS}}^2 + \sigma_{\text{PO}}^2$ | $\sigma_{\text{FS}}^2 + 2\sigma_{\text{PO}}^2$ |
| 1C, immature | $\sigma_{\text{PO}}^2 + \sigma_{\text{FS}}^2 + \sigma_{\text{PO}}^2$ | $\sigma_{\text{FS}}^2 + 2\sigma_{\text{PO}}^2$ |
| AV, immature | $\sigma_{\text{FS}}^2 + \sigma_{\text{PO}}^2$ | $\sigma_{\text{FS}}^2 + \sigma_{\text{PO}}^2$ |
| H2C, ovipositional | $\sigma_{\text{PO}}^2 + \sigma_{\text{PO}}^2 + \sigma_{\text{PO}}^2 + \sigma_{\text{HS}}^2 + \sigma_{\text{PO}}^2 + \sigma_{\text{PO}}^2 + \sigma_{\text{PO}}^2$ | $\sigma_{\text{HS}}^2 + 6\sigma_{\text{PO}}^2$ |

Examples of distributions based on additive variances are given in Table 2. Any intergenerational dispersal kernels that include the FS relationship at a branch point (e.g. full-sib, avuncular, first cousins, etc.) must necessarily include a component of dispersal reflecting this oviposition kernel in addition to any lifespan-related dispersal events. Similarly, two immature HS are separated by their father’s breeding dispersal kernel and the combination of their mothers’ gravid and oviposition kernels; this extends to related categories such as half cousin, half-avuncular, and so forth. Across a two-dimensional landscape, this approach enables intergenerational dispersal to be modelled as the outcome of a series of draws from a bivariate probability distribution function that models each individual dispersal component with symmetric axial sigma σ.

### The kindisperse package: Kinship dispersal simulations

#### (i) Simulation design

The simulation functions contained in the kindisperse package enable the exploration and characterisation of many aspects of intergenerational dispersal kernels. We illustrate this by simulating dispersal in two dimensions using Gaussian, Laplace and variance-gamma (shapes 0.5 & 0.1) probability distributions. These distributions were chosen as they cover a wide range of kernel shapes. Gaussian distributions are common in classical population genetics, such as Wright’s derivation of neighbourhood size (Wright, 1946), while the Laplace distribution is a symmetrical extension of the longer-tailed exponential distribution. A variance-gamma distribution enables exploration of a wide variety of kernel shapes, including very long-tailed kernels, and in its symmetric form contains the Gaussian and Laplace distributions as special cases.

Dispersal simulation functions were built in R v4.0.3 (R Core Team, 2020). Gaussian axial distributions were sampled with the “rnorm” function from the R stats package (R Core Team, 2020). The bivariate Laplace distribution was accessed with the “rmvl” function from the package LaplacesDemon (LLC. Statisticat, 2020). A bivariate variance-gamma distribution was constructed through a Gaussian scale mixture of the univariate gamma distribution with the bivariate normal distribution (McNicholas, McNicholas, & Browne, 2017). This distribution was then rescaled to compensate for the effect of shape on gamma distribution variance. The univariate gamma distribution was sampled with the “rgamma” function from the R stats package, then mixed with a bivariate normal distribution produced with the “rmvn” function from the package LaplacesDemon (LLC. Statisticat, 2020).

All simulations used in this paper were conducted using kindisperse version 0.10.1 (Jasper, 2021) (archived at https://doi.org/10.5281/zenodo.5112802). ‘Simple’ simulations take a single draw from a dispersal distribution (e.g. σ^2^_PO_). ‘Composite’ simulations take multiple draws from across multiple dispersal distributions (e.g. σ^2^_init_ + σ^2^_breed_ + σ^2^_grav_ + σ^2^_ovi_) (see Figure 1c). Composite simulations first randomly seed ancestral individuals for each kin dyad within the simulation area, then calculate the final positions of the descendants using a series of random, independent draws from the underlying statistical kernels describing each intermediate dispersal stage across both sides of the pedigree. X and Y displacements are added to the ancestral coordinate position to produce the descendent coordinate positions. Finally, the Euclidean geographic distance is calculated between the two descendent individuals at their sampling point.

#### (ii) Simulation testing

We tested the performance of kindisperse to model different aspects of dispersal via simulations. We simulated the distributions of distances between kin dyads under different conditions, in each case setting σ_PO_ = 1 and simulating 10,000,000 kin dyads (Figure 1d-f). For each simulated kin dyad, an initial parental location was drawn uniformly from a rectangular sampling area. The final distribution of the kin dyad was determined through summing repeated kernel draws reflecting all kinship phase, pedigree and sampling stage information. Simulations were implemented in the “simulate_kindist_simple” and “simulate_kindist_composite” functions.

We initially explored the effects of the kernel probability distribution on the distribution of simulated PO dyads (Figure 1d). The four probability distributions – Gaussian, Laplace, and variance-gamma (shapes 0.5 & 0.1) – each have varying degrees of kurtosis. As the second moment of a probability distribution function, σ^2^ is independent of the shape of the function, and the same σ_PO_ can in principle be estimated from all three distributions. However, strongly leptokurtic distributions such as the variance-gamma distribution are disproportionately impacted by rare long distance dispersal events – while the most dispersed Gaussian PO dyad in the simulation within this distribution was separated by 5.7σ, the most dispersed variance-gamma (0.1) dyad was separated by 31σ. In real world applications, care should be taken to ensure sampling areas are large enough to adequately capture long distance dispersal events.

We next simulated the dispersion of dyads across three generations of dispersal (Figure 1e). The GG (grandparental) category represents the additive outcome of two PO dispersal processes and can be modelled 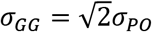. The inverse of the above, 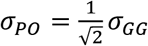, decomposes the GG distribution back to the important PO distribution. This can in principle be extended to arbitrary lifespan steps: so for example the GGG (great-grandparental) distribution could be decomposed to 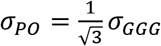, the fourth generation could be decomposed to 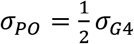, and so on.

Finally, we simulated the distributions of different kin categories (FS, HS, 1C) that are readily identifiable from samples taken in the field (Figure 1f). These were modelled as if collected as immatures. The spatial distribution of a FS dyad results from the ovipositional dispersal of their gravid mother, while that of a HS dyad results from a combination of their father’s breeding dispersal and each gravid mother’s dispersal post-mating. These distributions are out of phase. The fs and 1C distributions, however, are in phase; they are separated by a full lifespan of dispersal while sharing the same component of ovipositional dispersal, and thus can be collectively decomposed with the equation 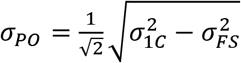.

### Validating close kin mark-recapture (CKMR) dispersal methods

#### (i) Neighbourhood (po) kernel estimation: Jasper et al. (2019)

The dispersal estimation method described in Jasper et al. (2019) decomposes the phase-specific elements associated with the groups FS, HS and 1C to supply an estimate of the axial standard deviation, σ, of the parent-offspring dispersal kernel, corresponding to Wright’s neighbourhood (Wright, 1946). Under an additive variance framework, this is achievable by independently estimating the dispersal kernels of multiple kin categories with the same kinship phase.

Two such phased kin groups are the immature fs and 1c distributions (Table 1). Both share the fs phase and have kernel distributions of *σ*^2^_*FS*_(FS) and *σ*^2^_*FS*_ + *2σ^2^_*PO*_* (1c). These can be combined to estimate the PO kernel: so, 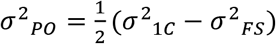, or 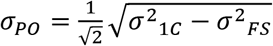. (equation #1 from Jasper et al. (2019)). For any kin dyad, the number of separating parent-offspring dispersal events and the phase of the dyad can be used to extend the Jasper et al. approach to a variety of parallel estimates of the fundamental intergenerational dispersal kernel, such as 1c & 2c, hs immature and ovipositional (Table 2), fs and av, fs & hs with a 1C/h1c mixture, and so on. These extensions of the method are implemented within kindisperse in the functions “axials_standard” or “axpermute_standard” for bootstrap-based confidence intervals.

#### (ii) Comparison of CKMR methods

We used kindisperse to compare two CKMR methods for estimating dispersal: the method of Jasper et al. (2019) described above, and the method described in Filipović et al. (2020). Dispersal estimation functions were implemented from both methods. The Filipović et al. method, initially applied to individuals sampled at the oviposition stage, involves identifying close kin dyads of the first, second and third orders. Its unique features are:

1. Attempting to account for dyads of the same kinship order that have different pedigree relationships (e.g. gg, av and hs which are all second order).
2. Attempting to account for dyads of the same pedigree relationship having different movement patterns, accomplished by enumerating the set of all possible dispersal events as a summation where “dispersed” = 1 and “did not disperse” = 0 across all lifespans within a pedigree.
3. Adjusting distance estimates for each kinship order by dividing raw distance measurements by all possible dispersal event counts as a result of points 1 and 2.

Supplementary Text 1 section 3 provides the code derived from Filipović et al. (2020) and discussion of that paper leading to this derivation.

For both methods, dispersal simulations were run for the fs, hs and 1C kin categories under Gaussian, Laplace, and variance-gamma (shape 0.5) kernel assumptions. In all cases, the axial deviation of the parent-offspring dispersal kernel (σ_PO_) was fixed at one, but the contributions of underlying kernels were allowed to vary. Specifically, the contributions of σ_init_, σ_(breed + grav_) and σ_ovi_ were individually iterated from 0.01σ to 0.99σ, with the remaining two categories equally sharing the remainder of the variance. Accordingly, the three kernels had equal contributions at 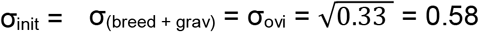. σ_breed_ and σ_grav_ were combined to reflect the hs class. Simulations were run with 10,000 kin dyads, from which 1,000 permutations of each treatment were run with sample sizes of 100 for the estimation of confidence intervals.

### Impact of sampling design on dispersal estimates

Estimating dispersal from the distribution of dispersed individuals can be vulnerable to biases if inadequate attention is given to sampling design. If intergenerational dispersal occurs over hundreds of metres it will not be detected if sampling is restricted to a 50m-by-50m square. We ran a series of simulations to assess the impact of variation in study design in empirical studies of dispersal. For these simulations, the simple PO kinship category was modelled under Gaussian, Laplace, and variance-gamma (shapes 0.5 & 0.1) distributions via the “simulate_kindist_simple” function, in each case setting σ_PO_ = 1 and simulating 10,000,000 kin dyads. For instance, the Laplace kernel was run using the command:

“simulate_kindist_simple(nsims = 10000000, sigma = 1, method = “Laplace”)”. Each dyad in this simulation constitutes a single draw from a bivariate symmetric dispersal kernel defined by axial sigma of 1. These distributions were then sampled with the “sample_kindist” function, in all cases randomly returning 10,000 individuals. For instance: “sample_kindist(kindist = x, n = 10000)”. For the complete code used to generate these sampling results, see Supplementary Text 1 section 4.

To begin with, we investigated the most basic constraints on a sampling site: the maximum upper and minimum lower distances between kin dyads in the dataset. Upper sampling range (a) was investigated across a range of 0 ≤ σ ≤ 5 with the sampling parameter “upper”. Lower sampling range (b) was investigated across a range of 0 ≤ σ ≤ 1 with the sampling parameter “lower”. Upper and lower sampling ranges supply simple estimates of our ability to recapture a dispersal kernel with a particular sampling scheme. However, when designing a study these parameters will not be known *a priori*. We therefore also investigated how sampling site geometry (size and shape) would affect estimates. We investigated overall sampling site size (c) by imposing a square sampling area with side length varying between 0 and 10σ. We investigated sampling site shape (d) by taking the area of the 10σ-sided square (i.e. 100σ^2^) and constructing a sampling rectangle of equivalent area but with an aspect ratio (ratio of lengths of sides) varying between 1:1 (square) and 100:1 (i.e. approaching linearity). Both these simulations were carried out with the sampling parameter “dims” and (for aspect ratio) the helper function “elongate”. Finally, we investigated the impact of the number of sampled kin dyads on confidence intervals (CIs) for each distribution, with the sampling parameter “n” varying between 1 and 250.

## Results

### The kindisperse package

The tools required to replicate all simulations and calculations within this paper are implemented in the kindisperse package. The key functions of the package are further implemented in an embedded shiny app for quick and intuitive application to dispersal estimation and sampling design. Specifically, this interface connects all simulation, sampling and estimation steps, enabling seamless exploration of kernel and site properties for improved study design. It can also be used to import and process field data, and to import and export results to and from the R programming environment or operating system. Detailed information on the use of this package is found in Supplementary Text 3.

### Validating close kin mark-recapture (CKMR) dispersal methods

A comparison of the Jasper et al. and Filipović et al. po kernel estimation methods is shown in Figure 2. These simulations consider kin categories fs, hs and 1c, with σ_PO_ = 1. Simulations model either immatures (Figure 2a,d,g) or gravid ovipositing females (Figure 2b,c,e,f,h,i). Within each simulation, one of the underlying dispersal kernels is allowed to vary (initial, Figure 2a-c; breeding & gravid, Figure 2d-f; oviposition, Figure 2g-i). The 95% confidence range of σ_PO_ is presented for each method using 1,000 permutations of 100 individuals in each.

**Figure 2.**
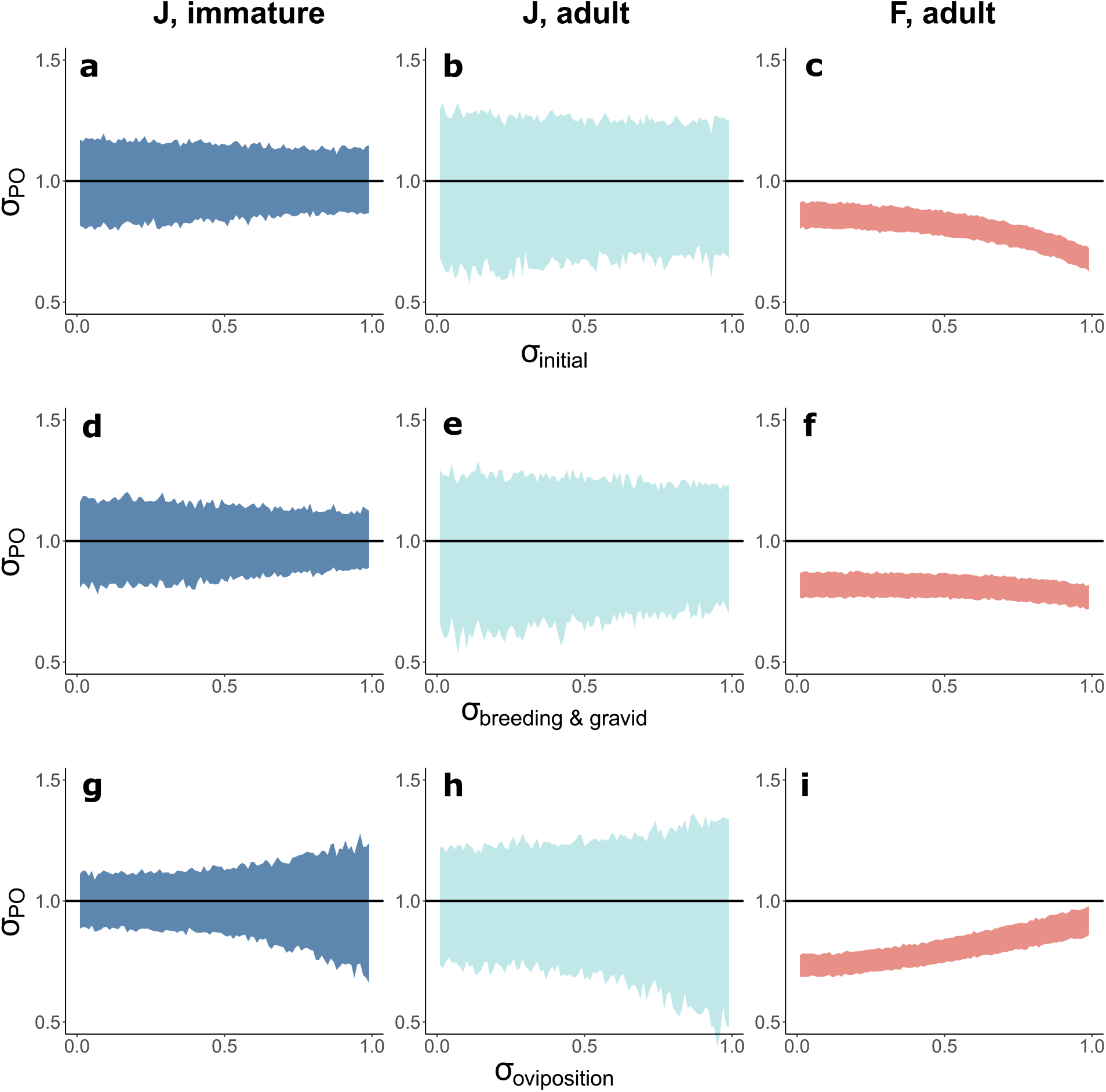
Comparisons of Jasper et al. and Filipović et al. methods for the estimation of σ_PO_. Simulations were conducted with an overall σ_PO_ of 1 and Laplace kernel shapes. For the Jasper et al. method, simulations were performed at the larval and adult (ovipositional) stages (first and second columns), whereas for the Filipović et al. method simulations were restricted to the adult stage (third column), as this method has not currently been adapted to the larval stage. For each simulation, different component stages of dispersal were varied between 0.01σ and 0.99σ, adjusting the other components to preserve the overall σ_PO_ of 1. Ribbon plot shows 95% confidence intervals for each estimate. **a-c.** Initial dispersal. **d-f.** Combined breeding and gravid dispersal. **g-i.** Oviposition dispersal.

For all simulations, the Jasper et al. method (based here on fs and 1C categories) consistently estimated σ_PO_ at close to the correct value of 1(Figure 2a,d,g & b,e,h). This method produced confidence intervals of ~0.6–1.25 for adult (ovipositional) dyads, and ~0.8–1.15 for immature dyads. Confidence intervals for these estimates widened with larger values of σ_ovi_. In contrast, the Filipović et al. method, based on combined _fs, hs_ and 1C distributions, consistently produced estimates of σ_PO_ with confidence intervals which failed to enclose the ‘true’σ_PO_ of 1. These estimates varied substantially in a manner dependent on the underlying dispersal distributions, from σ_PO_ ≈ 0.7 when σ_init_ was weighted higher than other dispersal elements to σ_PO_ ≈ 0.9 for when σ_ovi_ was weighted highest.

### Impact of sampling design on dispersal estimates

Maximum and minimum distances between kin dyads, the size and geometry of sample sites, and the number of close-kin dyads collected in the study all had an impact on dispersal estimates (Figure 3). These factors influenced results for all dispersal kernels, but were more pronounced for kernels that are more leptokurtic (i.e. dominated by long-distance dispersal).

**Figure 3.**
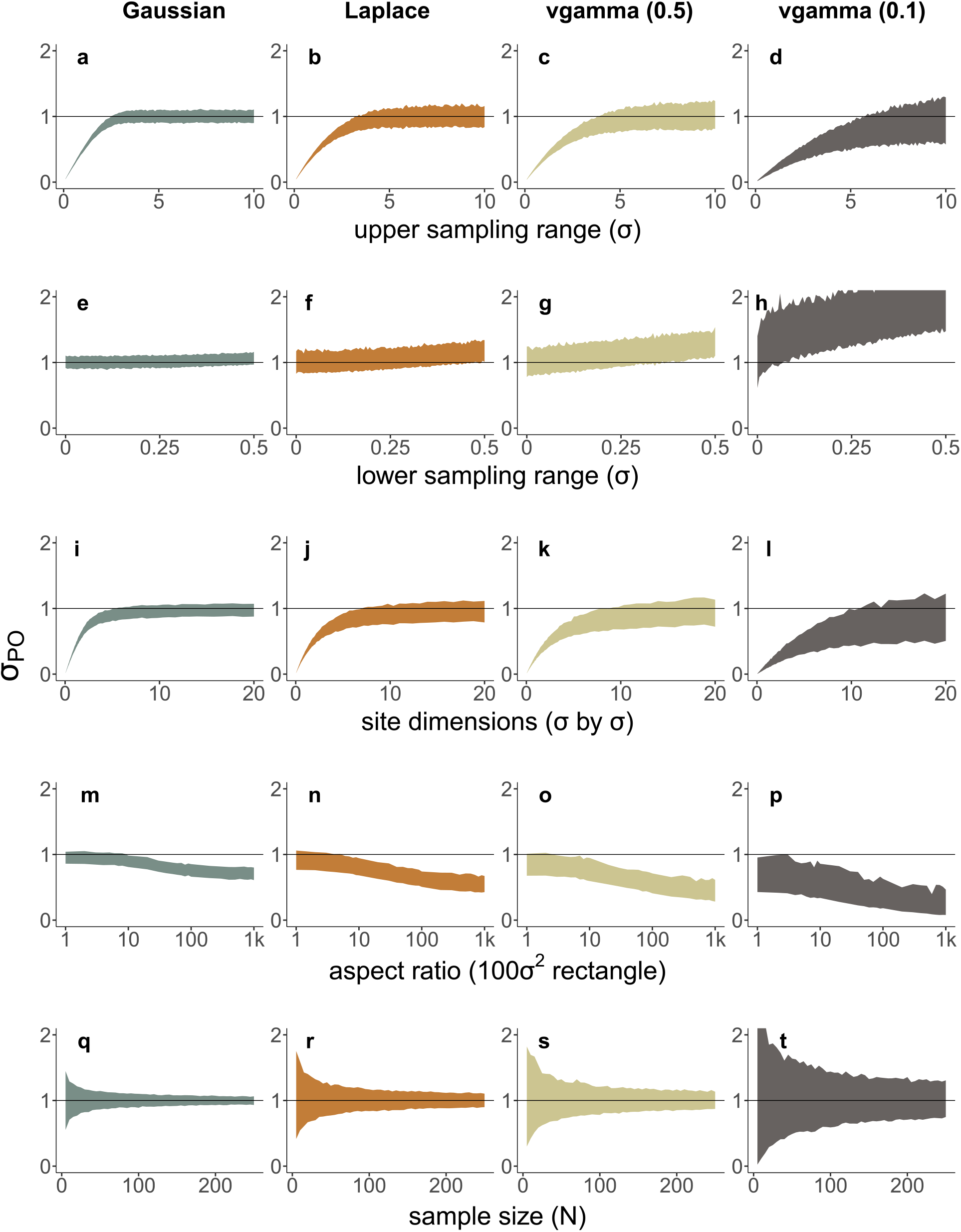
Effects of simulated field sampling conditions on estimates of σ_PO_. Each simulation has been replicated across four kernel shapes: Gaussian (column one), Laplace (column two), and variance-gamma (shape 0.5 column three, shape 0.1 column four). All simulations were conducted with a theoretical σ_PO_ of 1. **a-d.** Upper sampling range (maximum distance between kin dyads). **e-h.** Lower sampling range (minimum distance between kin dyads). **i-l.** Site dimensions (sides of square). **m-p.** Aspect ratio of 100σ^2^ rectangular site. **q-t.** Number of kin dyads sampled overall (testing variation in confidence intervals of estimates).

When the maximum sampling distance between kin dyads is insufficiently large relative to σ, dispersal estimates will be biased downwards (Figure 3a-d). The bias is relatively more severe for leptokurtic distributions, reflecting the higher frequency of long-distance dispersal events in these kernels. In this simulation, a maximum distance between kin dyads of >3σ was sufficient for the Gaussian kernel (Figure 3a); the Laplace kernel requires >5σ (Figure 3b); the variance-gamma kernels required >7σ and >10σ respectively (Figure 3c-d). This bias will exert the strongest influence on the largest dispersal kernel studied; for the fs, hs and 1C kin categories, this will be the 1C kernel.

When the minimum sampling distance between kin dyads is insufficiently small relative to σ, dispersal estimates will be biased upwards (Figure 3e-h). Leptokurtic kernels are again the most sensitive to this, as they have a higher proportion of samples near the origin. For unbiased results, the minimum required sampling distance between kin dyads was for the Gaussian kernel <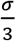 (Figure 3e), Laplace <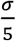 (Figure 3f), and the first variance-gamma <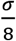 (Figure 3g). Extremely leptokurtic kernels, as in Figure 3h, remained biased across all lower sampling range parameters tested. This bias will exert the strongest influence on the smallest dispersal kernel studied; for the fs, hs and 1C kin categories, this will be the fs kernel.

Larger site dimensions are needed to estimate σ without downward bias (Figure 3i-l). Comparisons with the simple upper sampling limits in Figure 3a-d show that these site dimensions must enclose a much larger area than the maximum distance between kin. The Gaussian kernel required side lengths of ~5σ for accuracy (Figure 3i), while Laplace and the first variance-gamma kernels required ~10σ (Figure 3j-k). The extremely leptokurtic variance-gamma kernel required >15σ (3i). In practice, field sites intending to estimate σ_PO_ should have side lengths of >9σ_PO_ (see Supplementary Text 2 section 1 for further exposition of this).

As sampling within a 10σ-sided square was found to be adequate for estimating σ for less leptokurtic kernels, we next explored the effects of sampling within a 100σ^2^ rectangle of variable aspect ratio (Figure 3m-p). As the study site aspect ratio became increasingly elongated (y >> x), σ estimates increasingly dropped away from the correct value before eventually stabilising at a lower value (in the Gaussian case, 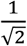 below the true value of σ (Figure 3m)). These stable but downward biased estimates of σ result from the linearization of the study site leading to one axial dimension being ‘stripped out’ of the data (i.e., a shift from a 2D to 1D site). If a study site is one-dimensional, such as sampling along a coastline, a different formulation of the dispersal neighbourhood is needed for accurate estimation (Wilkins & Wakeley, 2002; Wright, 1946). The overall shape of the dispersal kernel impacted the point at which the estimate stabilized (Figure 3n-p).

Finally, confidence interval ranges are impacted by the overall sample size in a way dependent on the underlying kernel (Figure 3q-t). For all sample sizes, more leptokurtic kernels are associated with wider confidence intervals. When > 100 kin dyads per category are included, σ estimates have confidence intervals of ~0.1σ for Gaussian distributions (3q). However, Gaussian estimates of ~0.2σ can still be obtained with < 30 kin dyads per category (3q). For kernels showing a moderate impact of long-distance dispersal, reasonable estimation confidence is preserved at 100 kin dyads (Figure 3r-s). Extremely leptokurtic kernels require many more samples to gain a similarly precise estimate (3t). Note that this simulation was run with PO kin groups – broader confidence intervals are expected for estimates derived from multiple kin categories.

## Discussion

In this paper we have presented kindisperse, an R package for simulating intergenerational dispersal and estimating dispersal parameters from simulated and empirical close kin data. This package has been developed in the context of *Ae. aegypti* dispersal studies. However, it has a much wider applicability in the field of insect dispersal and, provided careful consideration is given to underlying model assumptions, in investigations of other animals and of plants. We have illustrated the application of the package for assisting study design in two contexts: (a) a comparison of two CKMR methods for investigating intergenerational dispersal, and (b) an investigation of sampling requirements needed to ensure an accurate estimation of intergenerational dispersal.

### Comparison of estimation methods

This paper compared the dispersal estimation methods of Jasper et al. (2019) and Filipović et al. (2020), showing that the former provides more accurate estimates of σ_PO_ than the latter (Figure 2). The method of Filipović et al. was also subject to stronger influences from variation in the underlying components of dispersal.

A comparison of the theory underlying each method indicates why this is the case, and is especially important given the claims made in Filipović et al. (2020) about the superiority of their method vis-à-vis the method of Jasper et al. (2019).These claims are: that the Jasper et al. method operates under a restrictive assumption of Gaussian dispersal that is uncommon in biological systems (Claim 1); that the Jasper et al. method has wider confidence intervals and is thus less precise (Claim 2); and that the Jasper et al. method can produce outputs of imaginary numbers, which “raises concerns about the fundamental properties of their method” (Claim 3).

Claim 1 is partly addressed by the simulations in this paper, which show that the Jasper et al. method is not dependent on a Gaussian framework and can successfully handle other distributions, including Laplace and variance-gamma. More fundamentally, dispersal kernels are parametrised by the variance, σ^2^, which is independent of the underlying kernel shape. An estimate of σ_PO_ is therefore readily translatable from one kernel type to another.

The argument around precision in Claim 2 must be put in the context of the failure of the Filipović et al. method to successfully estimate σ_PO_ in any of our simulations. Upon examination, the apparent gains in precision achieved by this method result from an oversimplification of the underlying dispersal processes. In essence, the Filipović et al. method attempts to tease apart all of the component dispersal events that lead to the final distribution of the related individuals within each kinship category. Instead of first building an empirical understanding of these dispersal events, however, this decomposition is achieved by mapping out combinations of possible dispersal events, treating any such event as identical to any other such event, and then rescaling all distance values via simple division. Unfortunately, lacking a proper way to give likelihood-based weightings to different relationship categories within an order of kinship (e.g. gg vs hs), or careful enumeration of all such possible dispersal events, or for that matter a way of separating dispersal events of differing magnitudes, or even the correct method to decompose these distances (dispersal events sum together as variances, not as raw distances), the resulting estimates are biased and will have higher apparent precision only because they do not account for all of the sources of variability in the data. Additionally, the process of rescaling as described in Filipović et al. (Filipović et al., 2020) involves the generation of pseudoreplicates, as a single distance measure is converted into a “set of possible dispersal distances” – varying between two and four elements depending on kinship category – before all being pooled into one final dataset. This pseudoreplication should also inappropriately increase the apparent precision of estimates as it will artificially inflate the sample size without adding additional variability. Note that, in our implementation of this method, we removed this specific form of pseudoreplication by taking the mean of the set possible dispersal distances (Supplementary Text 1 section 3) which does not inflate sample size but still results in tighter confidence intervals than might be expected through a more realistic treatment such as through randomly selecting one distance.

To evaluate Claim 3, that the Jasper et al. method may produce imaginary numbers, it is necessary to understand the variance decomposition equation of the Jasper et al. method, namely 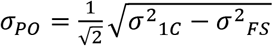. This equation treats fs dispersal as a subset of 1C dispersal, with the difference in variances attributable to parent-offspring dispersal across a lifetime. This equation would produce imaginary numbers in situations where the distribution of 1C overlaps with or is smaller than the distribution of fs. Far from calling into question the method, this feature is essential: if cousins are no further apart than siblings, there is no evidence that substantive intergenerational dispersal is occurring at all! Such a result could reflect a biological reality if the sampled population features absolute panmixia, though it might more likely be obtained when a sampling site is too small and does not capture the full extent of the 1c distribution. This issue underscores the need to design studies that cover an area sufficient to detect long range dispersal. We turn to this question next.

### Sampling design requirements

Our simulations show that sampling design can strongly bias estimates of intergenerational dispersal, and that these biases compound in the case of more leptokurtic kernels. At the one extreme, if sampling is conducted sparsely throughout a region, this low density may lead to minimum distances between kin being too high (>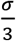 for Gaussian kernels), resulting in upwardly biased estimates. Importantly, this constraint applies to all kernel estimates, not just the anticipated po kernel, and thus trapping density should reflect the smallest anticipated kernel within the dataset. For the Jasper et al. approach applied to immature mosquitoes, this will typically be the fs kernel. As immature full-sibs are closely clustered, investigations of the fs kernel in immatures will require samples to be taken from nearby locations as well as multiple samples from the same location. At the other extreme, if sampling is conducted through too small a region, low maximum sampling distances (< 5σ for Gaussian kernels) will bias a kernel estimate downwards. As mosquito dispersal is frequently leptokurtic (Nathan et al., 2012), this means that study sites must be sufficiently large to capture the tails of the most dispersed kin kernel under investigation. Biases and limitations due to small study sites have been previously described for other dispersal methods (Guerra et al., 2014; Heuertz, Vekemans, Hausman, Palada, & Hardy, 2003) and in theoretical treatments (Rousset, 2001). As seen in Figure 3, it is difficult to accurately estimate σ using kin-based methods when kernels show extreme kurtosis, though close kin dyads can be used to detect recent dispersal over long distances (Schmidt et al., 2021).

All this considered, a typical study site would ideally aim for site dimensions approximately 9 times the length of the expected value of σ_PO_ (see Supplementary text 2 section 1 for a detailed discussion of these issues with respect to kinship category and kernel type). When selecting study sites, the results in Figure 3i-l and 3m-p also suggest that the length of individual site dimensions rather than merely site area is key to ensuring an accurate estimation of σ_po_. In all cases, small numbers of sampled kin increase the risk that an investigation will miss rare but important long-distance dispersal events. This is the reason why estimates involving leptokurtic kernels have larger confidence intervals (Figure 3q-t). These sources of potential bias underscore the need for dispersal studies to carefully consider what is already known about dispersal in a species alongside the statistical properties and limitations of dispersal estimators, such as those considered here. In practice, to avoid the risk of sample bias, a dispersal study based on kin should allow for a wide margin of error within each of the parameters when planning sampling for related individuals. These requirements need to be balanced against other issues; for instance, for mosquitoes, a high density of traps in an area might serve to attract mosquitoes from outside the area, decreasing the likelihood of detecting kin in a sample. Further checks should be performed after a study has been concluded to assess the extent to which the size of the sampling site may have biased estimates of σ_po_ (see Supplementary Text 2 section 2).

### Applications to non-mosquito species

The methods simulated and implemented in kindisperse were developed in the context of mosquito dispersal, a critically important parameter for planning interventions against mosquito borne diseases (Turelli & Barton, 2017). However, they should have wide applicability to the study of dispersal in other organisms, and in Supplementary Text 3 section 4.4 we describe a specific application to vertebrates for the genus *Antechinus*. The simulations and estimation procedures can be generalised to a range of dispersal and breeding situations and sampling life stages.

Several questions should be considered when adapting these methods for other taxa.

i. What are the breeding habits of the study species? *Aedes aegypti* females almost always mate only once, so half-sibs can be assumed to result from different mothers. For taxa where this is not the case, it will be necessary to modify the assumptions about the underlying dispersal biology represented by each kinship category.
ii. Is there overlap between generations in the study species? *Aedes aegypti* have short lifespans, so the simplifying assumptions can be made that immature full-sibs result from the same breeding cycle of the mother, and that second-order kin are most likely half-sibs rather than avuncular. These assumptions are particularly robust when sampling is conducted to select one generation specifically, such as when ovitraps are deployed and removed within the timespan of a single generation (Jasper et al., 2019; Schmidt et al., 2018). In more complex situations, generational overlap will need to be taken into account, particularly when dispersal varies with the age of an organism.
iii. What is the context in which dispersal is being considered? Dispersal estimates based on close kin are ideal for settings dominated by random, short-ranged dispersal and homogenous habitat, but the approach has limited value in understanding long distance migration events. For instance, a species may undergo seasonal migration and return to its natal region to reproduce; the approach might help to indicate whether kin produce nests in the same sample area but does not provide information on the long-distance route. In many contexts, additional factors such as wind-biased dispersal may also substantially undermine an assumption of isotropic dispersal. Animal behaviour is often driven by complex processes that may be shared across multiple individuals, again biasing estimates that assume independence and an additive variance model. All such potential biases and limitations should be carefully considered when using the kindisperse approach to estimate dispersal. Provided that kinship-based dispersal estimation methods are adapted carefully, they should be readily applicable to a wide range of taxa. For threatened species, a particular advantage of the approach is that dispersal can be estimated from DNA directly, which can be collected through non-invasive methods.

## Summary

In this paper, we introduced the R package kindisperse. Kindisperse allows for the simulation and estimation of intergenerational dispersal kernels from the spatial distribution of close kin dyads. We used kindisperse simulations to compare two recently developed methods for estimating dispersal kernels, and found that the Jasper et al. method (Jasper et al., 2019), upon which this package is built, provided unbiased estimates while the other method (Filipović et al., 2020) did not. We also applied the package to investigate the impact of various sampling parameters on kernel estimation and suggest design parameters to safeguard the reliability of future studies on intergenerational dispersal for kernels with an axial deviation defined by σ: minimum sampling distances <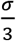, maximum distances in both dimensions > 9σ_po_, and at least 30 dyads of each kin category present in the final dataset.

## Acknowledgements

This research was conducted under Wellcome Trust Translation Award 108508/Z/15/Z and NHMRC Program Grant 1132412. Ary Hoffmann is supported by NHMRC Research Fellowship 1118640. The authors wish to thank Heng Lin Yeap (University of Melbourne) for invaluable advice on several statistical points.

## Data accessibility

A copy of KINDISPERSE v0.10.1 can be found at https://cran.r-project.org/web/packages/kindisperse/, and has also been permanently archived at https://doi.org/10.5281/zenodo.5112802; the most current version is hosted at https://github.com/moshejasper/kindisperse

## Author contributions

Moshe E. Jasper: designed research; performed research; ran simulations; wrote paper; edited paper; wrote R package

Ary A. Hoffmann: designed research; provided theory assistance; edited paper; provided supervision

Thomas L. Schmidt: designed research; provided theory assistance; wrote paper; edited paper; provided supervision

