## Supplementary Text 1 for "Estimating dispersal using close kin dyads: The kindisperse R package"

### Supplementary Text 1. Code used in Paper

Moshe E. Jasper, Ary A. Hoffmann, and Thomas L. Schmidt

23/07/2021

### 1. Setup

This code relies on (a) the tidyverse, (b) the package 'fitdistrplus' and (c) the package 'kindisperse'.

To install kindisperse, make sure you have the package 'devtools', then type

```
devtools::install_github("moshejasper/kindisperse")
```

then, load the needed packages:

```r
library(tidyverse)
library(fitdistrplus)
library(kindisperse)
```

The simulations and graphical outputs in this document take a long time to produce on a typical machine. The next panel contains a series of config values designed to control the scale of simulations run in this document. They are currently scaled to a tenth of the values used in the paper to conserve time. To reproduce the paper figures, alter these values by adding an additional zero.

```r
large_sim_num <- 1000000 # number of iterations for large simulation. paper value 10000000
small_sim_num <- 1000 # number of iterations for small simulations - paper value 10000
permutations <- 100 # number of iterations of permutation test - paper value 1000
```

#### 2. Initial kernel simulations

This code supplies the three simulations reported in Figure 1.

##### Kernel type

Here, we examine Gaussian, Laplace and variance-gamma (shape = 0.5) PO kernels. i.e. Figure 1d

```r
sim_gauss <- simulate_kindist_simple(nsims = large_sim_num, sigma = 1,
                                     method = "Gaussian")
sim_lap <- simulate_kindist_simple(nsims = large_sim_num, sigma = 1,
                                   method = "Laplace")
sim_gam <- simulate_kindist_simple(nsims = large_sim_num, sigma = 1,
                                   method = "vgamma", shape = 0.5)
sim_gamx <- simulate_kindist_simple(nsims = large_sim_num, sigma = 1,
                                    method = "vgamma", shape = 0.1)
# final simulation included for reference but not further implemented here. Substitute
# sim_gamx for sim_gam in subsequent code to explore the output of this extremely
# leptokurtic distribution.

kernelsims <- tibble(gauss = distances(sim_gauss), lap = distances(sim_lap),
                     gam = distances(sim_gam))
```

```r
ggplot(kernelsims) +
  geom_freqpoly(mapping = aes(x = gauss, colour = "Gaussian"),
                binwidth = 0.05, size = 1) +
  geom_freqpoly(mapping = aes(x = lap, colour = "Laplace"),
                binwidth = 0.05, size = 1) +
  geom_freqpoly(mapping = aes(x = gam, colour = "variance-gamma (shape 0.5)"),
                binwidth = 0.05, size = 1) +
  coord_cartesian(xlim = c(0, 5))+
  theme_classic() +
  xlab("kin dyad distance (m)") + ylab("count") +
  scale_colour_manual(breaks = c("Gaussian", "Laplace",
                                 "variance-gamma (shape 0.5)"),
                      values = c("black", "blue", "red"),
                      name = "kernel type")+
  theme(axis.text.y = element_blank(), axis.ticks.y = element_blank())
```

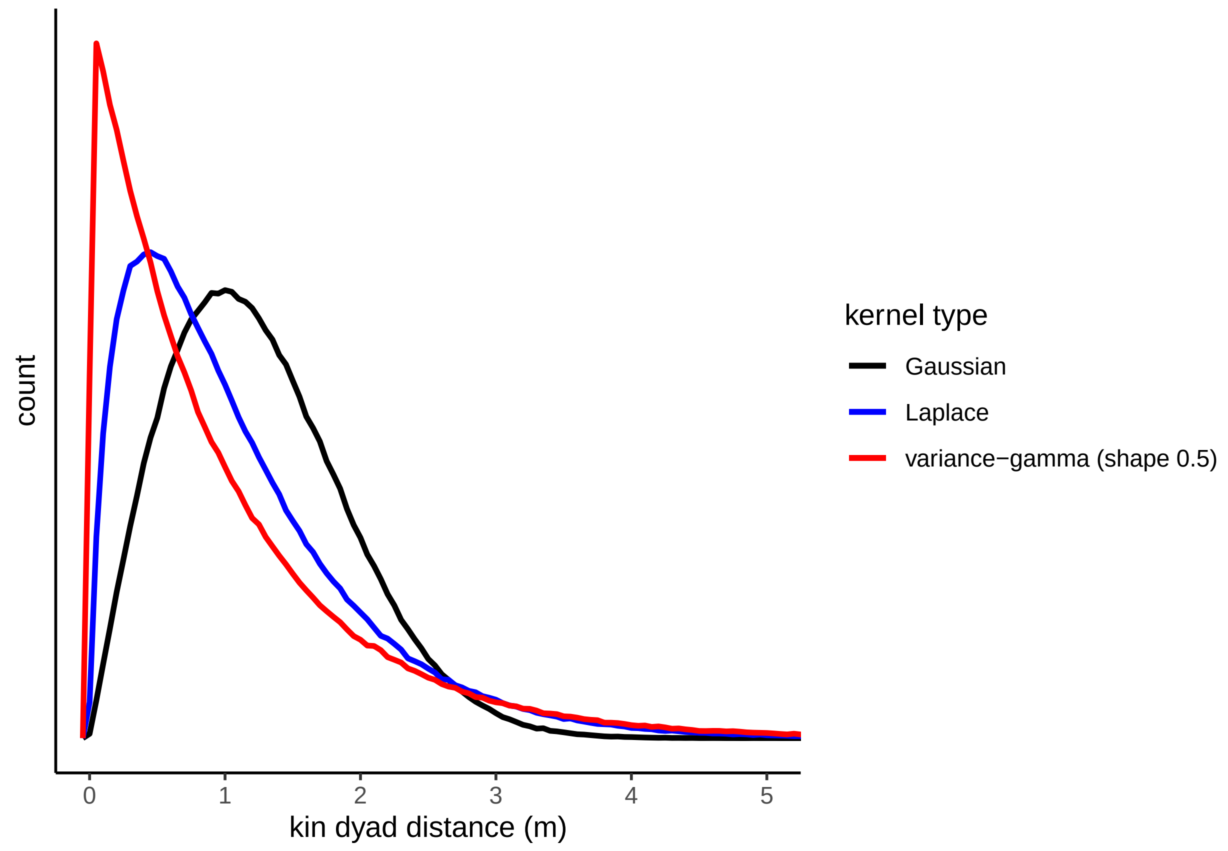

##### Offspring generation

Here, we examine the PO, GG, and GGG distributions in a Gaussian context (Figure 1e)

```
sim_po <- simulate_kindist_simple(nsims = large_sim_num, sigma = 1, kinship = "PO")
sim_gg <- simulate_kindist_simple(nsims = large_sim_num, sigma = 1, kinship = "GG")
sim_ggg <- simulate_kindist_simple(nsims = large_sim_num, sigma = 1, kinship = "GGG")

gensims <- tibble(po = distances(sim_po), gg = distances(sim_gg), ggg = distances(sim_ggg))
```

```
ggplot(gensims) +
  geom_freqpoly(mapping = aes(x = po, colour = "PO"), binwidth = 0.05, size = 1) +
  geom_freqpoly(mapping = aes(x = gg, colour = "GG"), binwidth = 0.05, size = 1) +
  geom_freqpoly(mapping = aes(x = ggg, colour = "GGG"), binwidth = 0.05, size = 1) +
  coord_cartesian(xlim = c(0, 5))+
  theme_classic() +
  xlab("kin dyad distance (m)") + ylab("count") +
  scale_colour_manual(breaks = c("PO", "GG", "GGG"), values = c("black", "blue", "red"),
                      name = "generation")+
  theme(axis.text.y = element_blank(), axis.ticks.y = element_blank())
```

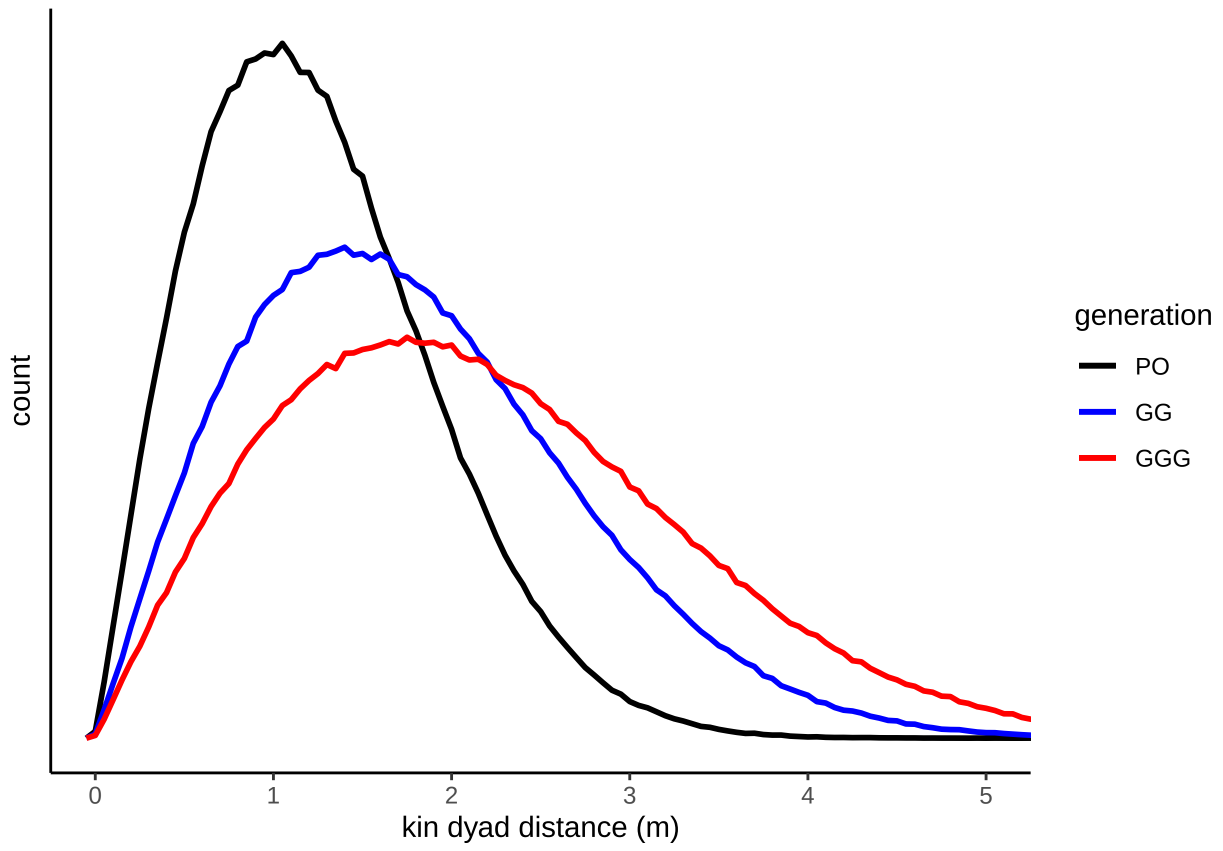

##### Kin category

Finally, we examine three typical kin categories (FS, HS, 1C) all sampled as immatures. A kernel formed by two draws from the PO kernel (i.e. dispersal of two siblings hatched at the identical location) is shown for comparison. (Figure 1f)

```
sim_gg2 <- simulate_kindist_composite(nsims = large_sim_num, initsigma = 0.5,
                                      breedsigma = 0.5,  gravsigma = 0.5,
                                      ovisigma = 0.5, kinship = "GG")
# GG is equivalent to two draws from PO
sim_fs <- simulate_kindist_composite(nsims = large_sim_num, initsigma = 0.5,
                                     breedsigma = 0.5, gravsigma = 0.5,
                                     ovisigma = 0.5, kinship = "FS")
sim_hs <- simulate_kindist_composite(nsims = large_sim_num, initsigma = 0.5,
                                     breedsigma = 0.5, gravsigma = 0.5,
                                     ovisigma = 0.5, kinship = "HS")
sim_1c <- simulate_kindist_composite(nsims = large_sim_num, initsigma = 0.5,
                                     breedsigma = 0.5, gravsigma = 0.5,
                                     ovisigma = 0.5, kinship = "1C")

kinsims <- tibble(po = distances(sim_gg2), fs = distances(sim_fs),
                  hs = distances(sim_hs), c1 = distances(sim_1c))
```

```
ggplot(kinsims) +
  geom_freqpoly(mapping = aes(x = po, colour = "PO (2 draws)"), binwidth = 0.05,
                linetype = 2) +
  geom_freqpoly(mapping = aes(x = fs, colour = "FS"), binwidth = 0.05, size = 1) +
  geom_freqpoly(mapping = aes(x = hs, colour = "HS"), binwidth = 0.05, size = 1) +
  geom_freqpoly(mapping = aes(x = c1, colour = "1C"), binwidth = 0.05, size = 1) +
  coord_cartesian(xlim = c(0, 5))+
  theme_classic() +
  xlab("kin dyad distance (m)") + ylab("count") +
  scale_colour_manual(breaks = c("PO (2 draws)", "FS", "HS", "1C"),
                      values = c("black", "blue", "red", "darkorange"),
                      name = "kin category")+
  theme(axis.text.y = element_blank(), axis.ticks.y = element_blank())
```

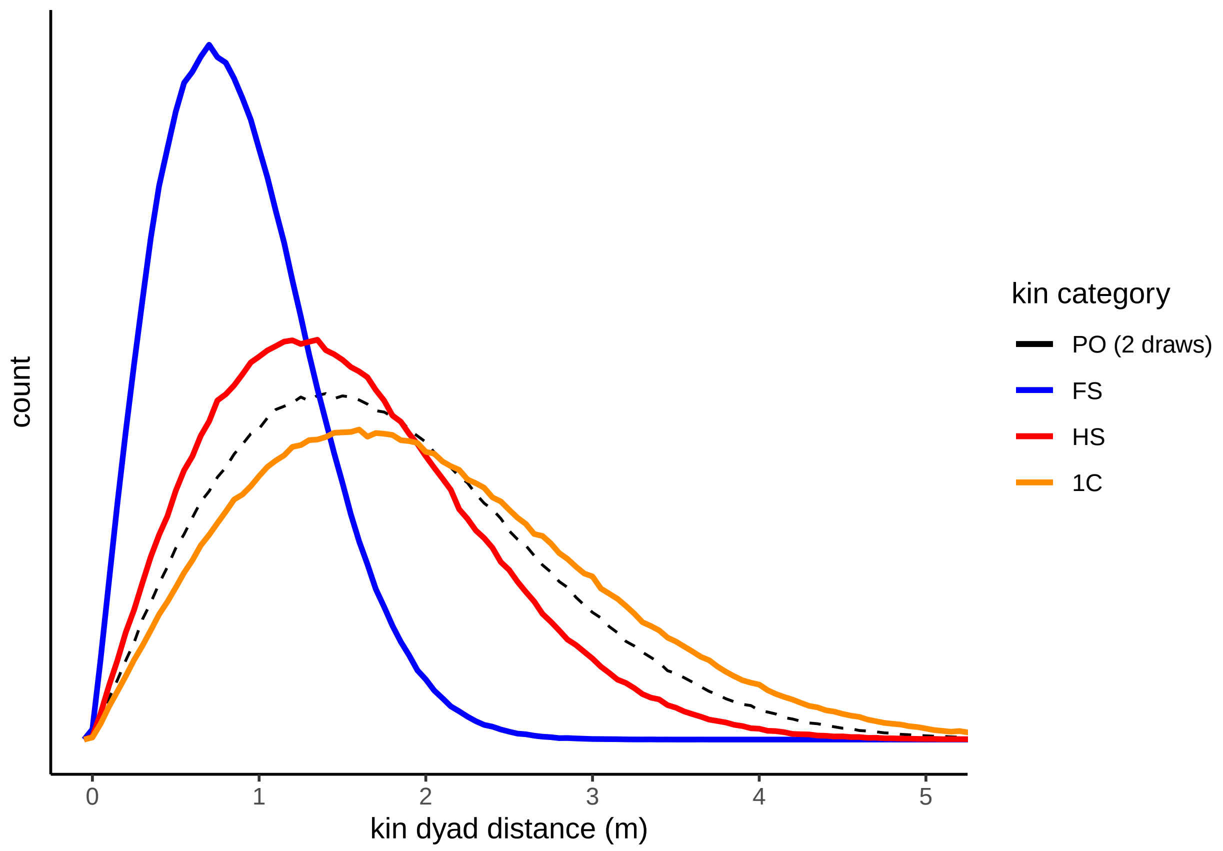

#### 3. Method comparisons

This code supplies the three simulations reported in Figure 2, comparing the Jasper and Filipovic methods for PO kernel estimation

##### Implementing the Filipovic et al. method

The functions implemented in this section are derived from the methodology advocated in Filipovic et al. 2020, "Using spatial genetics to quantify mosquito dispersal for control programs".

As there is no algorithmic implementation of their method, in this section explain how we implemented functions derived from their methodology.

The intention of their method is to 'decompose... the observed separation distances between close kin (sampled as breeding adults) to generate the distribution of **potential effective dispersal distances** and to parametrize the dispesal distance kernel' (emphasis added). They collected adult females using sticky traps (i.e. trapping females during oviposition).

They separated genotyped kin dyads into three categories: full-sibling, 2nd-order relatives, and 3rd-order relatives.

They define the category they are after as 'effective dispersal distance'; that is, 'a distance between the birthplace and the ovipositing place of a female (or a distance between the ovipositing place of a mother and a daughter').

They begin by recognizing that all close-kin categories contain information about the dispersal events that produce them. From here, their reasoning diverges from that in the present paper. Its key features are as follows:

1. (they claim) this information corresponds to the *number*(s) of possible breeding and dispersal events over one generation.
2. For each kin category, there are a set of possible dispersal events that produced the final distribution (so for HS adults, there are either two (the dispersals of each adult) or three (those dispersals plus the skip oviposition of the mother)). Note that this set describes the *number* of dispersal events, not their *quality*, i.e. this scheme treats the oviposition dispersal of the mother as of equal weight with the dipsersal of her daughter across the entire lifespan.
3. These numbers can be represented as sets and combined to form set unities. So, For the second order kinship (which they assume to be ambiguous), the HS = {2, 3, 4, 5}, AV = {3, 4} and GG = {2, 3} are combined together to form a unity set, {2, 3, 4, 5} - which contains all elements of the other sets (and in fact all elements of the original HS set!). Constructed like this, the final union set gives no indication of the *frequencies* of each of the dispersal modes, merely their *possibility*. Various underlying dispersal routes within a category could be described with the same number, and these are not all equally frequent. Furthermore, a consistent usage of this method would in all likelihood allow for several missed numbers. The method was only applied to adult mosquitoes - the difficulties with deploying the assumptions described here rendered it impossible for us to adapt the method to the larval stage for comparison with our own datasets.

Using this process, they first construct defining sets for each of the first, second and third order kin categories (implemented below).

```
FSet <- c(2, 3)
HSet <- c(2, 3, 4, 5)
CSet <- c(2, 3, 4, 5)
```

4. Having identified this set of possible events, they then use them to 'divide the observed spatial distance between each close-kin dyad to create a set of possible effective dispersal distance'. To understand this method, it is important to note that this is a simple division operation: two full-siblings treated as resulting from the same larval site (i.e. FSet 2) that are separated by 100m would be considered to result from two 50m effective dispersal events. In practice, using the FSet values, they would be averaged between 50m and 33m (leading to an estimate of 45m). Rather than treating two-dimensional dispersal events as probabilistically resulting from random dispersal described by a statistical kernel, at this stage the method of Filipovic in fact treats dispersal as happening in discrete events along a line which can be divided evenly between different dispersed movements without redundancy.
5. The division described in step 4 (in the original method) is actually division by a multi-element vector - the set of distances created by division 'contains the same number of elements as the corresponding set

of possible effective dispersal events'. This means the 100m separation described previously is passed down to future analysis steps not as '45m', but in fact as '50m and 33m' (2 datapoints). For The HS (2nd order) and 1C (3rd) categories, with more possible combinations, one datapoint will produce 4 pseudoreplicated datapoints after this step.

6. This pseudoreplication issue is locked in as they combine '... the values from **all pairwise sets** of possible effective dispersal distances into one dataset'.
7. This combined dataset was then fitted with the R package 'fitdistrplus' to a probability distribution (they tried multiple, but found the exponential distribution to fit best).
8. They then performed estimates from (a) individual kinship classes, and (b) combined kinship classes.

We have implemented the above method for the Filipovic paper in six functions (three pairs), shown below.

Note that each pair contains an original version which replicates the Filipovic method as closely as possible, and an adjusted version which avoids the creation of pseudoreplicates. We used the adjusted version in this paper to enable a more meaningful comparison of confidence intervals (which were strongly skewed by the duplication).

The basic function (implemented below) takes a vector of separating distance `valvect`, divides them by the dispersal event set `dcats`, then fits this to an exponential model and returns the derived value of sigma.

Its adjusted version divides each dispersal distance by the mean of the dispersal set rather than the set to avoid pseudoreplication, but is otherwise identical.

```
Filipovic_estimate_basic <- function(valvect, dcats, method = "exp"){

  newvals <- numeric()

  for (val in dcats) {
    newvals <- c(newvals, valvect / val)}
  return(1 / mledist(newvals, method)$estimate)
}

Filipovic_estimate_adjusted <- function(valvect, dcats, method = "exp"){

  facts <- mean(1 / dcats)
  newvals <- valvect * facts
  return(1 / mledist(newvals, method)$estimate)
}
```

The next pair combine information from all three available categories (1st, 2nd, 3rd) to estimate dispersal. They are otherwise similar to the previous pair.

```
Filipovic_combined <- function(v1, v2, v3, s1 = FSet, s2 = HSet, s3 = CSet,
                               method = "exp"){

  newvals <- numeric()

  for (val in s1) {
    newvals <- c(newvals, v1 / val)
  }

  for (val in s2){
    newvals <- c(newvals, v2 / s2)
  }
  for (val in s3){
```

```
    newvals <- c(newvals, v3 / s3)
  }
  return(1 / mledist(newvals, method)$estimate)
}

Filipovic_combined_adjusted <- function(v1, v2, v3, s1 = FSet, s2 = HSet, s3 = CSet,
                                        method = "exp"){

  f1 <- mean(1 / s1)
  f2 <- mean(1 / s2)
  f3 <- mean(1 / s3)

  newvals <- c(v1 * f1, v2 * f2, v3 * f3)

  return(1 / mledist(newvals, method)$estimate)

}
```

The final pair implement permutation bootstrapping to enable the calculation of basic confidence intervals (for comparisons with the Japser et al. method). Medians and 95% C.I.s are returned.

```
Filipovic_estimate_permute <- function(valvect, dcats, method = "exp", nreps = 1000,
                                       num = 50){

  container <- tibble(sig = 0.0, .rows = 0)

  for (val in 1:nreps){
    subvals <- sample(valvect, num, replace = T)
    newsig <- Filipovic_estimate_adjusted(subvals, dcats, method = method)
    container <- container %>% add_row(sig = newsig)
  }
  return(quantile(container$sig, c(0.025, 0.5, 0.975)))
}

Filipovic_combined_permute <- function(v1, v2, v3, s1 = FSet, s2 = HSet, s3 = CSet,
                                       method = "exp", nreps = 1000,
                                       num = 100, output = "vect"){

  container <- tibble(sig = 0.0, .rows = 0)

  for (val in 1:nreps){
    sub1 <- sample(v1, num, replace = T)
    sub2 <- sample(v2, num, replace = T)
    sub3 <- sample(v3, num, replace = T)
    newsig <- Filipovic_combined_adjusted(sub1, sub2, sub3, s1, s2, s3, method = method)
    container <- container %>% add_row(sig = newsig)
  }
  if (output == "confs"){
    return(quantile(container$sig, c(0.025, 0.5, 0.975)))
  }
  else {
    return(sort(container$sig))
  }
```

```
}
```

#### Initial dispersal

Figure 2a-c focuses on varying the initial dispersal kernel as a component of overall PO dispersal.

```
initdata <- tibble(init = 0, fl = 0, fm = 0, fu = 0, lower = 0, mid = 0, upper = 0,
                           limm = 0, mimm = 0, uimm = 0, .rows = 0)

for (init in 1:99/100){

  # set up states...
  bg <-  sqrt((1 - init^2) / 2)
  ovi <- bg
  breed <-  sqrt(bg^2 / 2)
  grav <- breed
  simfs_ovi <- simulate_kindist_composite(nsims = small_sim_num, initsigma = init,
                                          breedsigma = breed, gravsigma = grav,
                                      ovisigma = ovi, method = "Laplace",
                                      kinship = "FS", lifestage = "ovipositional")
  simhs_ovi <- simulate_kindist_composite(nsims = small_sim_num, initsigma = init,
                                          breedsigma = breed, gravsigma = grav,
                                      ovisigma = ovi, method = "Laplace",
                                      kinship = "HS", lifestage = "ovipositional")
  sim1c_ovi <- simulate_kindist_composite(nsims = small_sim_num, initsigma = init,
                                          breedsigma = breed, gravsigma = grav,
                                      ovisigma = ovi, method = "Laplace",
                                      kinship = "1C", lifestage = "ovipositional")
  simfs_imm <- simulate_kindist_composite(nsim = small_sim_num, initsigma = init,
                                          breedsigma = breed, gravsigma = grav,
                                      ovisigma = ovi, method = "Laplace",
                                      kinship = "FS", lifestage = "immature")
  sim1c_imm <- simulate_kindist_composite(nsims = small_sim_num, initsigma = init,
                                          breedsigma = breed, gravsigma = grav,
                                      ovisigma = ovi, method = "Laplace",
                                      kinship = "1C", lifestage = "immature")
  result_ovi <- as.vector(axpermute_standard(sim1c_ovi, simfs_ovi,
                                             nreps = permutations, nsamp = 100))
  result_imm <- as.vector(axpermute_standard(sim1c_imm, simfs_imm,
                                             nreps = permutations, nsamp = 100))
  fi_result_ovi <- as.vector(Filipovic_combined_permute(distances(simfs_ovi),
                                                        distances(simhs_ovi),
                                                        distances(sim1c_ovi),
                                                        nreps = permutations,
                                                        num = 100, output = "confs"))
  initdata <- initdata %>%
    add_row(init = init, lower = result_ovi[1], mid = result_ovi[2],
            upper = result_ovi[3], limm = result_imm[1],
            mimm = result_imm[2], uimm = result_imm[3],
            fl = fi_result_ovi[1], fm = fi_result_ovi[2], fu = fi_result_ovi[3])
}
```

```r
ggplot(initdata) + aes(x = init) +
  geom_ribbon(mapping = aes(ymin = lower, ymax = upper), fill = "grey90")+
  geom_ribbon(mapping = aes(ymin = fl, ymax = fu), fill = "grey75")+
  geom_line(mapping = aes(y = mid), size = 1, colour = "blue") +
  coord_cartesian(ylim = c(0.5, 1.5)) +
  #geom_line(mapping = aes(y = fl), colour = "red") +
  #geom_line(mapping = aes(y = fu), colour = "red") +
  geom_line(mapping = aes(y = limm), linetype = 2, colour = "blue")+
  geom_line(mapping = aes(y = uimm), linetype = 2, colour = "blue")+
  geom_line(mapping = aes(y = lower), linetype = 3, colour = "black") +
  geom_line(mapping = aes(y = upper), linetype = 3, colour = "black") +
  geom_line(mapping = aes(y = fm), colour = "red", size = 1) +
  geom_hline(yintercept = 1, linetype = 2, size = 1, colour = "black")+
  theme_classic() +
  xlab(expression(sigma["initial"])) + ylab(expression(sigma["PO"]))
```

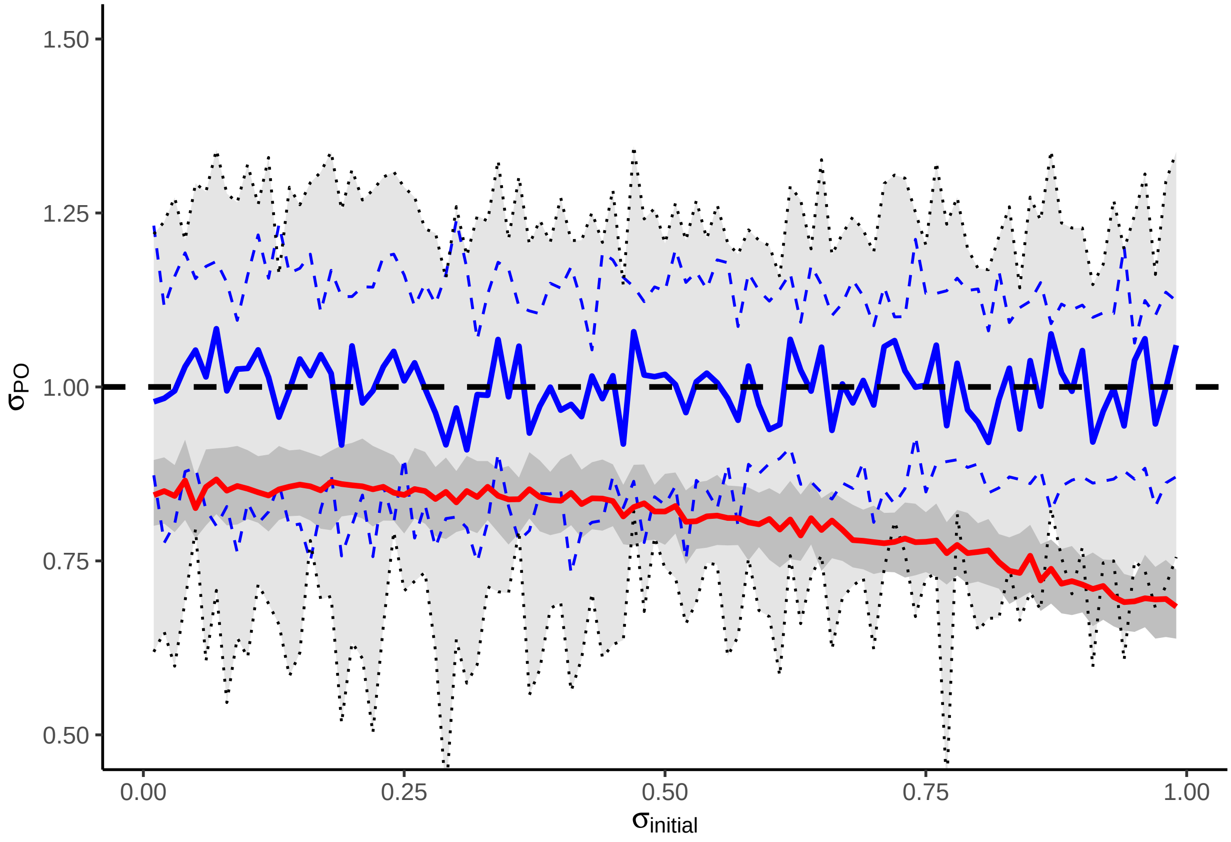

#### Breeding and gravid dispersal

Figure 2d-f focuses on varying the breeding and gravid dispersal kernels as components of overall PO dispersal

```r
bgdata <- tibble(bg = 0, fl = 0, fm = 0, fu = 0, lower = 0, mid = 0, upper = 0,
                 limm = 0, mimm = 0, uimm = 0, .rows = 0)

for (n in 1:99 / 100){
```

```r
  bg <- n

  init <-  sqrt((1 - bg^2) / 2)
  ovi <- init
  breed <-  sqrt(bg^2 / 2)
  grav <- breed
  simfs_ovi <- simulate_kindist_composite(nsim = small_sim_num, initsigma = init,
                                          breedsigma = breed, gravsigma = grav,
                                          ovisigma = ovi, method = "Laplace",
                                          kinship = "FS", lifestage = "ovipositional")
  simhs_ovi <- simulate_kindist_composite(nsims = small_sim_num, initsigma = init,
                                          breedsigma = breed, gravsigma = grav,
                                          ovisigma = ovi, method = "Laplace",
                                          kinship = "HS", lifestage = "ovipositional")
  sim1c_ovi <- simulate_kindist_composite(nsim = small_sim_num, initsigma = init,
                                          breedsigma = breed, gravsigma = grav,
                                          ovisigma = ovi, method = "Laplace",
                                          kinship = "1C", lifestage = "ovipositional")
  simfs_imm <- simulate_kindist_composite(nsim = small_sim_num, initsigma = init,
                                          breedsigma = breed, gravsigma = grav,
                                          ovisigma = ovi, method = "Laplace",
                                          kinship = "FS", lifestage = "immature")
  sim1c_imm <- simulate_kindist_composite(nsim = small_sim_num, initsigma = init,
                                          breedsigma = breed, gravsigma = grav,
                                          ovisigma = ovi, method = "Laplace",
                                          kinship = "1C", lifestage = "immature")
  result_ovi <- as.vector(axpermute_standard(sim1c_ovi, simfs_ovi,
                                             nreps = permutations, nsamp = 100))
  result_imm <- as.vector(axpermute_standard(sim1c_imm, simfs_imm,
                                             nreps = permutations, nsamp = 100))
  fi_result_ovi <- as.vector(Filipovic_combined_permute(distances(simfs_ovi),
                                                        distances(simhs_ovi),
                                                        distances(sim1c_ovi),
                                                        nreps = permutations,
                                                        num = 100, output = "confs"))
  bgdata <- bgdata %>% add_row(bg = bg, lower = result_ovi[1],
                               mid = result_ovi[2], upper = result_ovi[3],
                               limm = result_imm[1], mimm = result_imm[2],
                               uimm = result_imm[3], fl = fi_result_ovi[1],
                               fm = fi_result_ovi[2], fu = fi_result_ovi[3])
}
```

```r
ggplot(bgdata) + aes(x = bg) +
  geom_ribbon(mapping = aes(ymin = lower, ymax = upper), fill = "grey90")+
  geom_ribbon(mapping = aes(ymin = fl, ymax = fu), fill = "grey75")+
  geom_line(mapping = aes(y = mid), size = 1, colour = "blue") +
  coord_cartesian(ylim = c(0.5, 1.5)) +
  #geom_line(mapping = aes(y = fl), colour = "red") +
  #geom_line(mapping = aes(y = fu), colour = "red") +
  geom_line(mapping = aes(y = limm), linetype = 2, colour = "blue")+
  geom_line(mapping = aes(y = uimm), linetype = 2, colour = "blue")+
  geom_line(mapping = aes(y = lower), linetype = 3, colour = "black") +
  geom_line(mapping = aes(y = upper), linetype = 3, colour = "black") +
```

```
  geom_line(mapping = aes(y = fm), colour = "red", size = 1) +
  geom_hline(yintercept = 1, linetype = 2, size = 1, colour = "black")+
  theme_classic() +
  xlab(expression(sigma["breeding and gravid"])) + ylab(expression(sigma["PO"]))
```

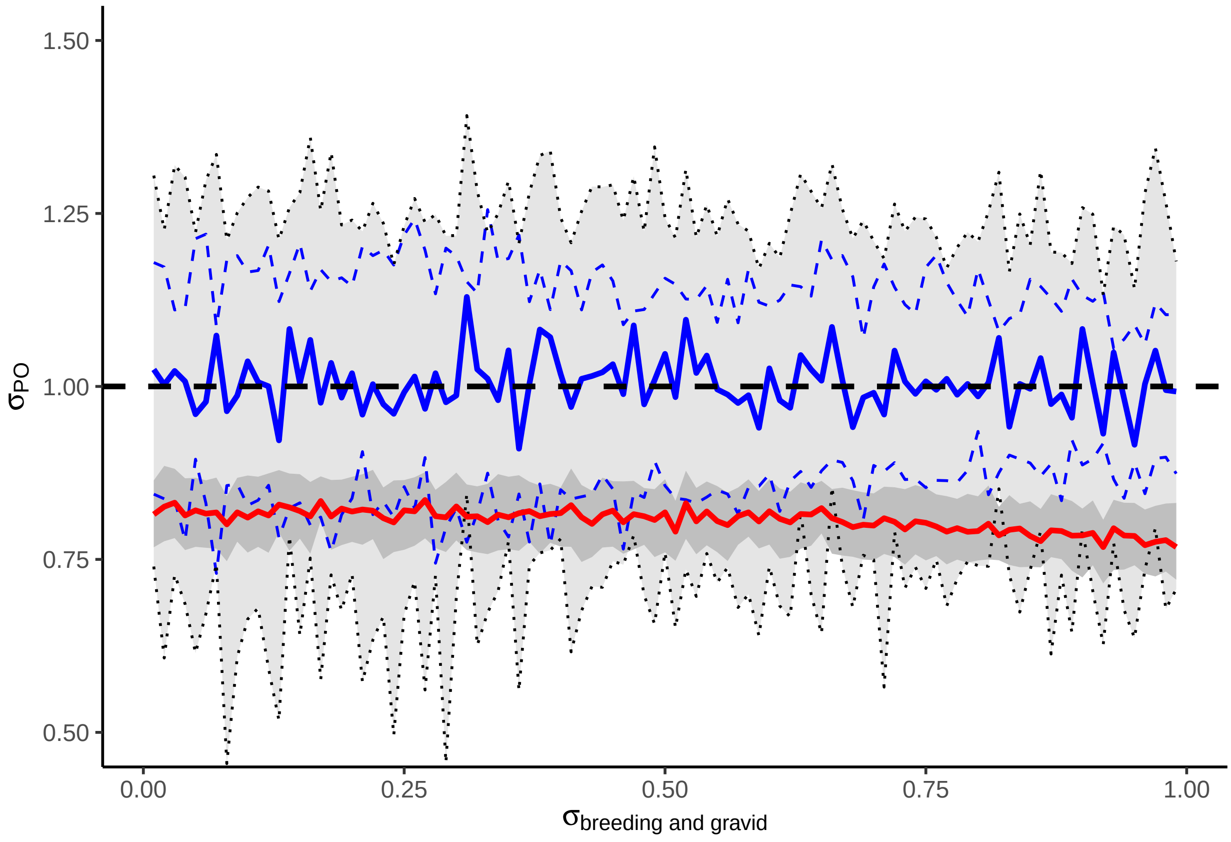

#### Oviposition dispersal

Figure 2g-i focuses on varying the oviposition dispersal kernel as a component of overall PO dispersal

```
ovidata <- tibble(ovi = 0, fl = 0, fm = 0, fu = 0, lower = 0, mid = 0, upper = 0,
                  limm = 0, mimm = 0, uimm = 0, .rows = 0)

for (n in 1:99 / 100){

  ovi <- n

  init <-  sqrt((1 - ovi^2) / 2)
  #ovi <- init
  breed <-  sqrt(init^2 / 2)
  grav <- breed
  simfs_ovi <- simulate_kindist_composite(nsim = small_sim_num, initsigma = init,
                                          breedsigma = breed,
                                          gravsigma = grav, ovisigma = ovi, method = "Laplace",
                                          kinship = "FS", lifestage = "ovipositional")
```

```r
  simhs_ovi <- simulate_kindist_composite(nsims = small_sim_num, initsigma = init,
                                          breedsigma = breed, gravsigma = grav,
                                          ovisigma = ovi, method = "Laplace",
                                          kinship = "HS", lifestage = "ovipositional")
  sim1c_ovi <- simulate_kindist_composite(nsim = small_sim_num, initsigma = init,
                                          breedsigma = breed, gravsigma = grav,
                                          ovisigma = ovi, method = "Laplace",
                                          kinship = "1C", lifestage = "ovipositional")
  simfs_imm <- simulate_kindist_composite(nsim = small_sim_num, initsigma = init,
                                          breedsigma = breed, gravsigma = grav,
                                          ovisigma = ovi, method = "Laplace",
                                          kinship = "FS", lifestage = "immature")
  sim1c_imm <- simulate_kindist_composite(nsim = small_sim_num, initsigma = init,
                                          breedsigma = breed, gravsigma = grav,
                                          ovisigma = ovi, method = "Laplace",
                                          kinship = "1C", lifestage = "immature")
  result_ovi <- as.vector(axpermute_standard(sim1c_ovi, simfs_ovi,
                                             nreps = permutations, nsamp = 100))
  result_imm <- as.vector(axpermute_standard(sim1c_imm, simfs_imm,
                                             nreps = permutations, nsamp = 100))
  fi_result_ovi <- as.vector(Filipovic_combined_permute(distances(simfs_ovi),
                                                         distances(simhs_ovi),
                                                         distances(sim1c_ovi),
                                                         nreps = permutations,
                                                         num = 100, output = "confs"))
  ovidata <- ovidata %>%
    add_row(ovi = ovi, lower = result_ovi[1], mid = result_ovi[2], upper = result_ovi[3],
            limm = result_imm[1], mimm = result_imm[2], uimm = result_imm[3],
            fl = fi_result_ovi[1], fm = fi_result_ovi[2], fu = fi_result_ovi[3])
}
```

```r
ggplot(ovidata) + aes(x = ovi) +
  geom_ribbon(mapping = aes(ymin = lower, ymax = upper), fill = "grey90")+
  geom_ribbon(mapping = aes(ymin = fl, ymax = fu), fill = "grey75") +
  geom_line(mapping = aes(y = mid), size = 1, colour = "blue") +
  coord_cartesian(ylim = c(0.5, 1.5)) +
  #geom_line(mapping = aes(y = fl), colour = "red") +
  #geom_line(mapping = aes(y = fu), colour = "red") +
  geom_line(mapping = aes(y = limm), linetype = 2, colour = "blue")+
  geom_line(mapping = aes(y = uimm), linetype = 2, colour = "blue")+
  geom_line(mapping = aes(y = lower), linetype = 3, colour = "black") +
  geom_line(mapping = aes(y = upper), linetype = 3, colour = "black") +
  geom_line(mapping = aes(y = fm), colour = "red", size = 1) +
  geom_hline(yintercept = 1, linetype = 2, size = 1, colour = "black")+
  theme_classic() +
  xlab(expression(sigma["oviposition"])) + ylab(expression(sigma["PO"]))
```

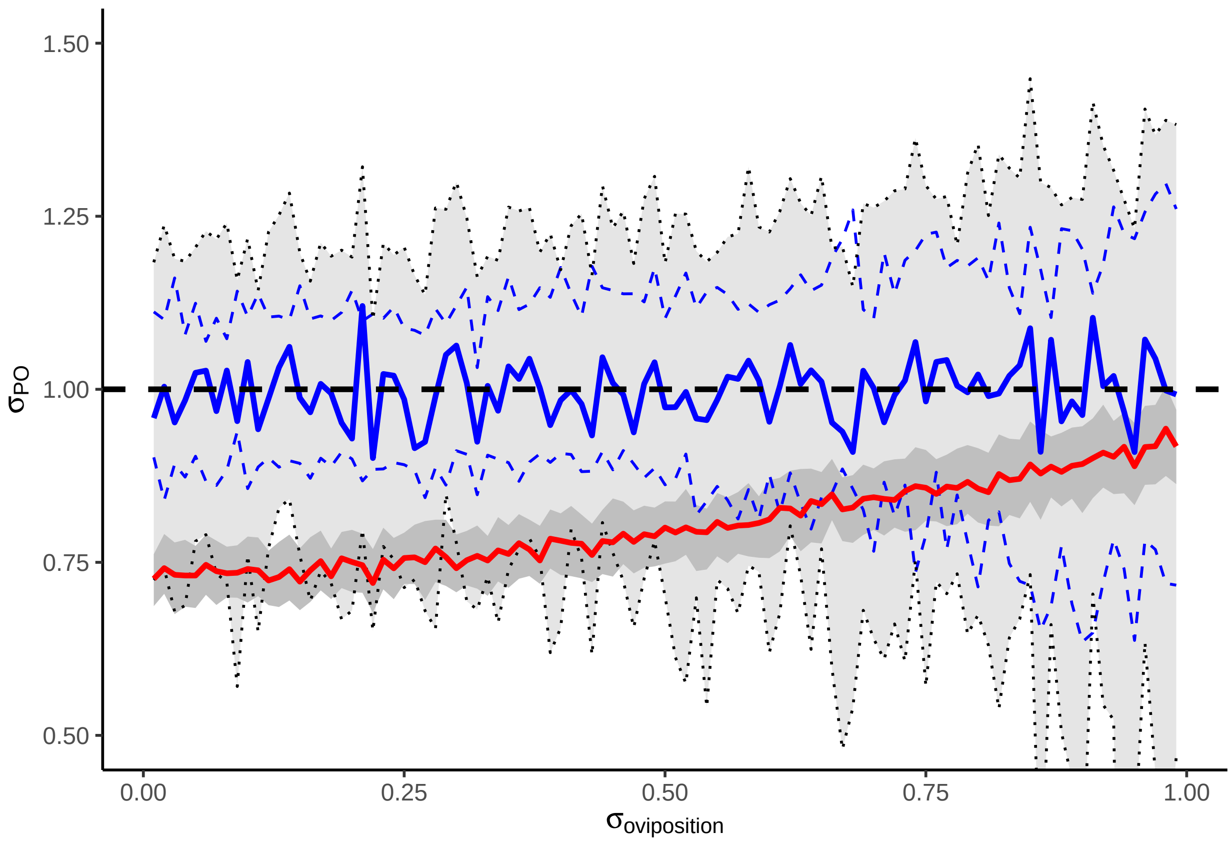

#### 4. Site sampling design

This code supplies the simulations reported in Figure 3, examining the impact of site sampling parameters on estimates of dispersal sigma (only the code for the first three columns is replicated here: rows are combined into composite figures).

##### Maximum sampling distance

Figure 3a-d focuses on maximum sampling distance between kin dyads

```
simbase_gauss <- simulate_kindist_simple(nsims = large_sim_num, sigma = 1, dims = 1000,
                                         method = "Gaussian")
simbase_laplace <- simulate_kindist_simple(nsims = large_sim_num, sigma = 1, dims = 1000,
                                           method = "Laplace")
simbase_vgamma <- simulate_kindist_simple(nsims = large_sim_num, sigma = 1, dims = 1000,
                                          method = "vgamma", shape = 0.5)
```

```
uppercheck <- tibble(n = 0, l = 0, m = 0, u = 0, ll = 0, ml = 0, ul = 0, lg = 0, mg = 0,
                     ug = 0, .rows = 0)

for (n in 1:50 / 10){
  egauss <- simbase_gauss %>% sample_kindist(upper = n, n = small_sim_num) %>%
    axpermute(nreps = permutations, nsamp = 100, composite = 1) %>% as.vector()
```

```
  elap <- simbase_laplace %>% sample_kindist(upper = n, n = small_sim_num) %>%
    axpermute(nreps = permutations, nsamp = 100, composite = 1) %>% as.vector()
  egam <- simbase_vgamma %>% sample_kindist(upper = n, n = small_sim_num) %>%
    axpermute(nreps = permutations, nsamp = 100, composite = 1) %>% as.vector()
  uppercheck <- uppercheck %>% add_row(n = n, l = egauss[1], m = egauss[2],
                                       u = egauss[3], ll = elap[1], ml = elap[2],
                                       ul = elap[3], lg = egam[1], mg = egam[2],
                                       ug = egam[3])
}
```

```
ggplot(uppercheck) + aes(x = n) +
  geom_ribbon(mapping = aes(ymin = lg, ymax = ug), fill = "#ffc5c5")+
  geom_ribbon(mapping = aes(ymin = ll, ymax = ul), fill = "#bbbbff", alpha = 0.7)+
  geom_ribbon(mapping = aes(ymin = l, ymax = u), fill = "#eeeeee", alpha = 1) +
  geom_ribbon(mapping = aes(ymin = lg, ymax = ug), fill = NA, colour = "red", linetype = 3)+
  geom_ribbon(mapping = aes(ymin = ll, ymax = ul), fill = NA, colour = "blue", linetype = 3)+
  geom_ribbon(mapping = aes(ymin = l, ymax = u), fill = NA, colour = "black", linetype = 3) +
  geom_line(mapping = aes(y = m), size = 1)+
  geom_line(mapping = aes(y = ml), colour = "blue", size = 1) +
  geom_line(mapping = aes(y = mg), colour = "red", size = 1) +
  geom_hline(yintercept = 1, linetype = 2, size = 1) +
  theme_bw()+
  coord_cartesian(ylim = c(0, 2), xlim = c(0, 5))+
  xlab("upper sampling range") + ylab(expression(sigma["PO"]))
```

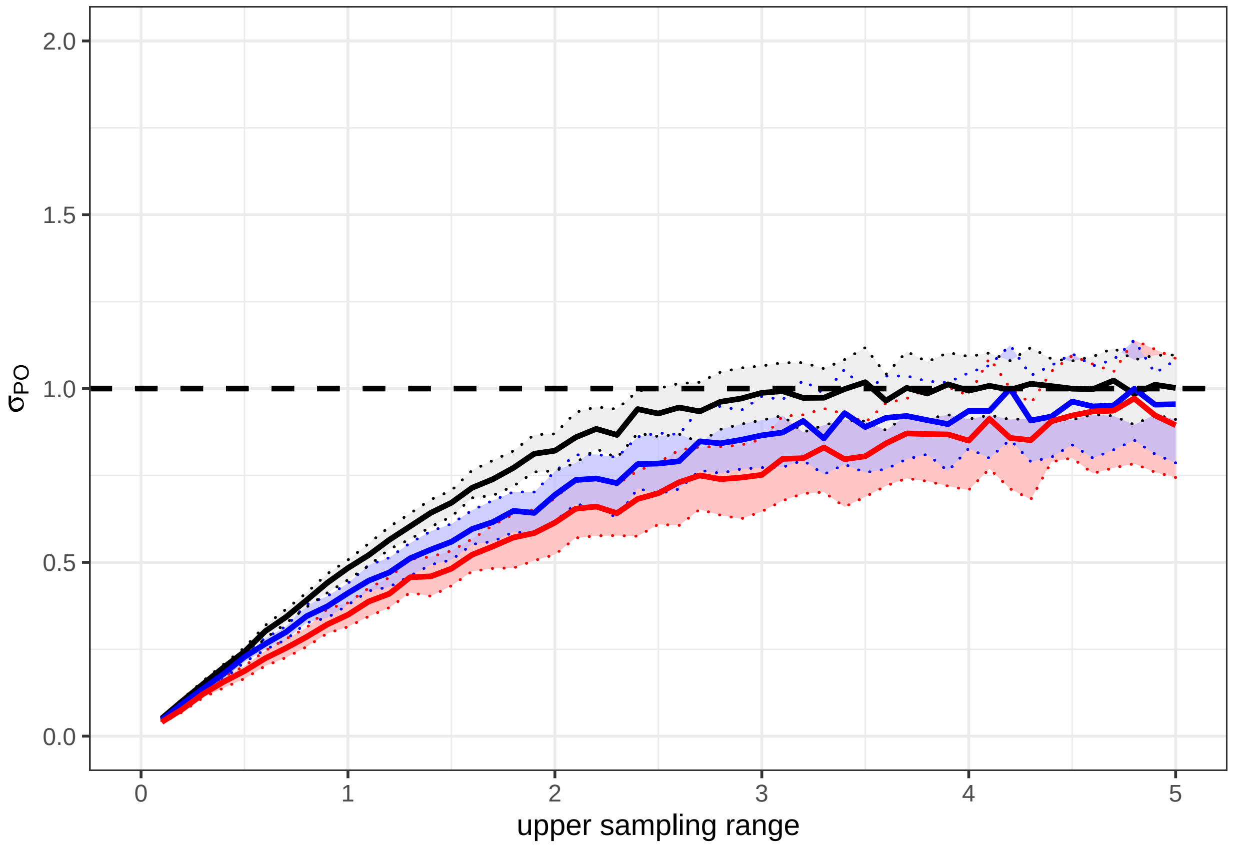

#### Minimum sampling distance

Figure 3e-h focuses on minimum sampling distance between kin dyads

```
lowercheck <- tibble(n = 0, l = 0, m = 0, u = 0, ll = 0, ml = 0, ul = 0, lg = 0, mg = 0,
                     ug = 0, .rows = 0)

for (n in 0:50 / 50){
  egauss <- simbase_gauss %>% sample_kindist(lower = n, n = small_sim_num) %>%
    axpermute(nreps = permutations, nsamp = 100, composite = 1) %>% as.vector()
  elap <- simbase_laplace %>% sample_kindist(lower = n, n = small_sim_num) %>%
    axpermute(nreps = permutations, nsamp = 100, composite = 1) %>% as.vector()
  egam <- simbase_vgamma %>% sample_kindist(lower = n, n = small_sim_num) %>%
    axpermute(nreps = permutations, nsamp = 100, composite = 1) %>% as.vector()
  lowercheck <- lowercheck %>% add_row(n = n, l = egauss[1], m = egauss[2],
                                       u = egauss[3], ll = elap[1], ml = elap[2],
                                       ul = elap[3], lg = egam[1], mg = egam[2],
                                       ug = egam[3])
}
```

```
ggplot(lowercheck) + aes(x = n) +
  geom_ribbon(mapping = aes(ymin = lg, ymax = ug), fill = "#ffc5c5")+
  geom_ribbon(mapping = aes(ymin = ll, ymax = ul), fill = "#bbbbff", alpha = 0.7)+
  geom_ribbon(mapping = aes(ymin = l, ymax = u), fill = "#eeeeee", alpha = 0.7) +
  geom_ribbon(mapping = aes(ymin = lg, ymax = ug), fill = NA, colour = "red",
              linetype = 3, size = 0.75)+
  geom_ribbon(mapping = aes(ymin = ll, ymax = ul), fill = NA, colour = "blue",
              linetype = 3, size = 0.75)+
  geom_ribbon(mapping = aes(ymin = l, ymax = u), fill = NA, colour = "black",
              linetype = 3, size = 0.75) +
  geom_line(mapping = aes(y = m), size = 1)+
  geom_line(mapping = aes(y = ml), colour = "blue", size = 1) +
  geom_line(mapping = aes(y = mg), colour = "red", size = 1) +
  geom_hline(yintercept = 1, linetype = 2, size = 1) +
  coord_cartesian(ylim = c(0, 2), xlim = c(0, 1))+
  theme_bw()+
  xlab("lower sampling range") + ylab(expression(sigma["PO"])) #+
```

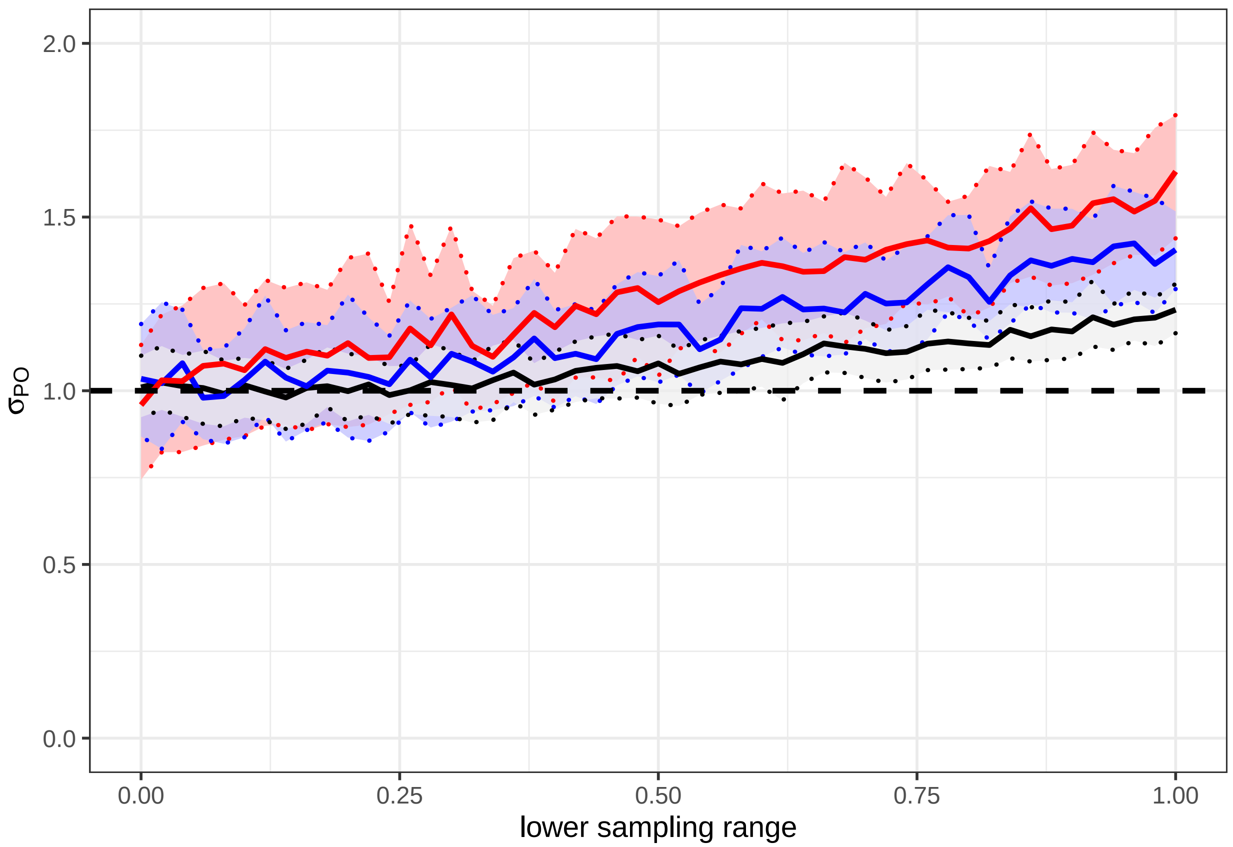

#### Sampling site dimensions

Figure 3i-l focuses on the dimensions of a square sampling site varying between sidelengths of $0 - 10\sigma$

```
eval <- log(10)
nval <- unique((exp((-25:25 / 25) * eval)))
dimensioncheck <- tibble(n = 0, l = 0, m = 0, u = 0, ll = 0, ml = 0, ul = 0, lg = 0,
                         mg = 0, ug = 0, .rows = 0)

for (n in nval){
  egauss <- simbase_gauss %>% sample_kindist(dims = n, n = small_sim_num) %>%
    axpermute(nreps = permutations, nsamp = 100, composite = 1) %>% as.vector()
  elap <- simbase_laplace %>% sample_kindist(dims = n, n = small_sim_num) %>%
    axpermute(nreps = permutations, nsamp = 100, composite = 1) %>% as.vector()
  egam <- simbase_vgamma %>% sample_kindist(dims = n, n = small_sim_num) %>%
    axpermute(nreps = permutations, nsamp = 100, composite = 1) %>% as.vector()
  dimensioncheck <- dimensioncheck %>% add_row(n = n, l = egauss[1], m = egauss[2],
                                               u = egauss[3], ll = elap[1], ml = elap[2],
                                               ul = elap[3], lg = egam[1], mg = egam[2],
                                               ug = egam[3])
}
```

```
ggplot(dimensioncheck) + aes(x = n) +
  geom_ribbon(mapping = aes(ymin = lg, ymax = ug), fill = "#ffcccc", colour = "red",
              linetype = 2)+
```

```
  geom_ribbon(mapping = aes(ymin = ll, ymax = ul), fill = "#ccccff", alpha = 0.7,
              colour = "blue", linetype = 2)+
  geom_ribbon(mapping = aes(ymin = l, ymax = u), fill = "#ffffff", alpha = 0.7,
              colour = "black", linetype = 2) +
  geom_line(mapping = aes(y = m), size = 1)+
  geom_line(mapping = aes(y = ml), colour = "blue", size = 1) +
  geom_line(mapping = aes(y = mg), colour = "red", size = 1) +
  geom_hline(yintercept = 1, linetype = 2, size = 1) +
  scale_x_continuous(breaks = c(0, 2, 4, 6, 8, 10))+
  theme_bw() +
  coord_cartesian(ylim = c(0, 2)) +
  xlab(expression(paste("site dimensions (", sigma, " by ", sigma, ")"))) +
  ylab(expression(sigma["PO"]))# +
```

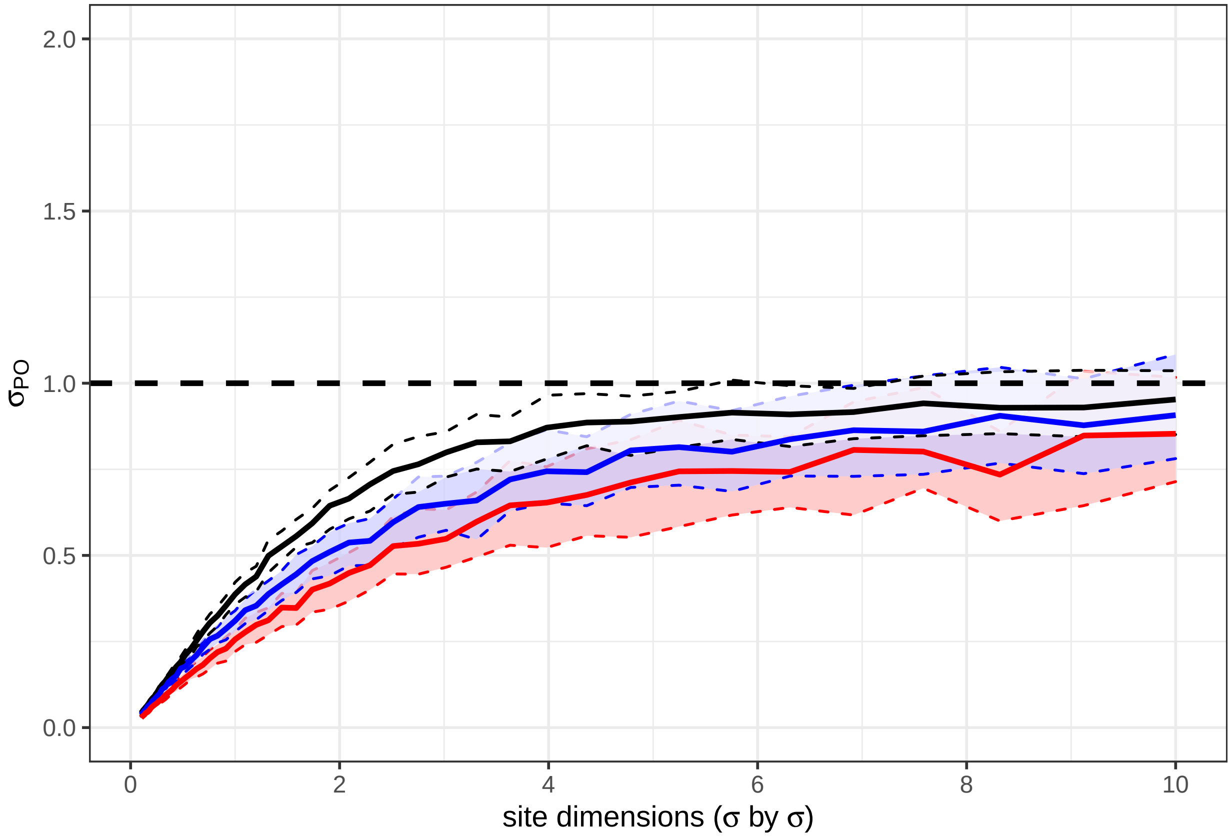

##### Sampling site aspect ratio

Figure 3m-p focuses on the aspect ratio (ratio of sides) of a $100\sigma^2$ rectangle.

```
eval <- log(10000)
nval <- unique(round(exp((1:50 / 50) * eval)))

elongationcheck <- tibble(n = 0, l = 0, m = 0, u = 0, ll = 0, ml = 0, ul = 0, lg = 0, mg = 0,
                          ug = 0, .rows = 0)
```

```r
for (n in c(1:10, 2:10*10)){
  egauss <- simbase_gauss %>% sample_kindist(dims = elongate(10, n), n = small_sim_num) %>%
    axpermute(nreps = permutations, nsamp = 100, composite = 1) %>% as.vector()
  elap <- simbase_laplace %>% sample_kindist(dims = elongate(10, n), n = small_sim_num) %>%
    axpermute(nreps = permutations, nsamp = 100, composite = 1) %>% as.vector()
  egam <- simbase_vgamma %>% sample_kindist(dims = elongate(10, n), n = small_sim_num) %>%
    axpermute(nreps = permutations, nsamp = 100, composite = 1) %>% as.vector()
  elongationcheck <- elongationcheck %>%
    add_row(n = n, l = egauss[1], m = egauss[2], u = egauss[3],
            ll = elap[1], ml = elap[2], ul = elap[3], lg = egam[1], mg = egam[2],
            ug = egam[3])
}
```

```r
ggplot(elongationcheck) + aes(x = n) +
  geom_ribbon(mapping = aes(ymin = lg, ymax = ug), fill = "#ffcccc", colour = "red",
              linetype = 2)+
  geom_ribbon(mapping = aes(ymin = ll, ymax = ul), fill = "#ccccff", alpha = 0.7,
              colour = "blue", linetype = 2)+
  geom_ribbon(mapping = aes(ymin = l, ymax = u), fill = "#ffffff", alpha = 0.7,
              colour = "black", linetype = 2) +
  geom_line(mapping = aes(y = m), size = 1)+
  geom_line(mapping = aes(y = ml), colour = "blue", size = 1) +
  geom_line(mapping = aes(y = mg), colour = "red", size = 1) +
  geom_hline(yintercept = 1, linetype = 2, size = 1) +
  scale_x_log10(breaks = c(1, 3, 10, 30, 100),
                labels = c("1:1", "3:1", "10:1", "30:1", "100:1"))+
  theme_bw() +
  coord_cartesian(ylim = c(0, 2))+
  xlab(expression(paste("aspect ratio (100", sigma^2, " rectangle)")))  +
  ylab(expression(sigma["PO"]))# +
```

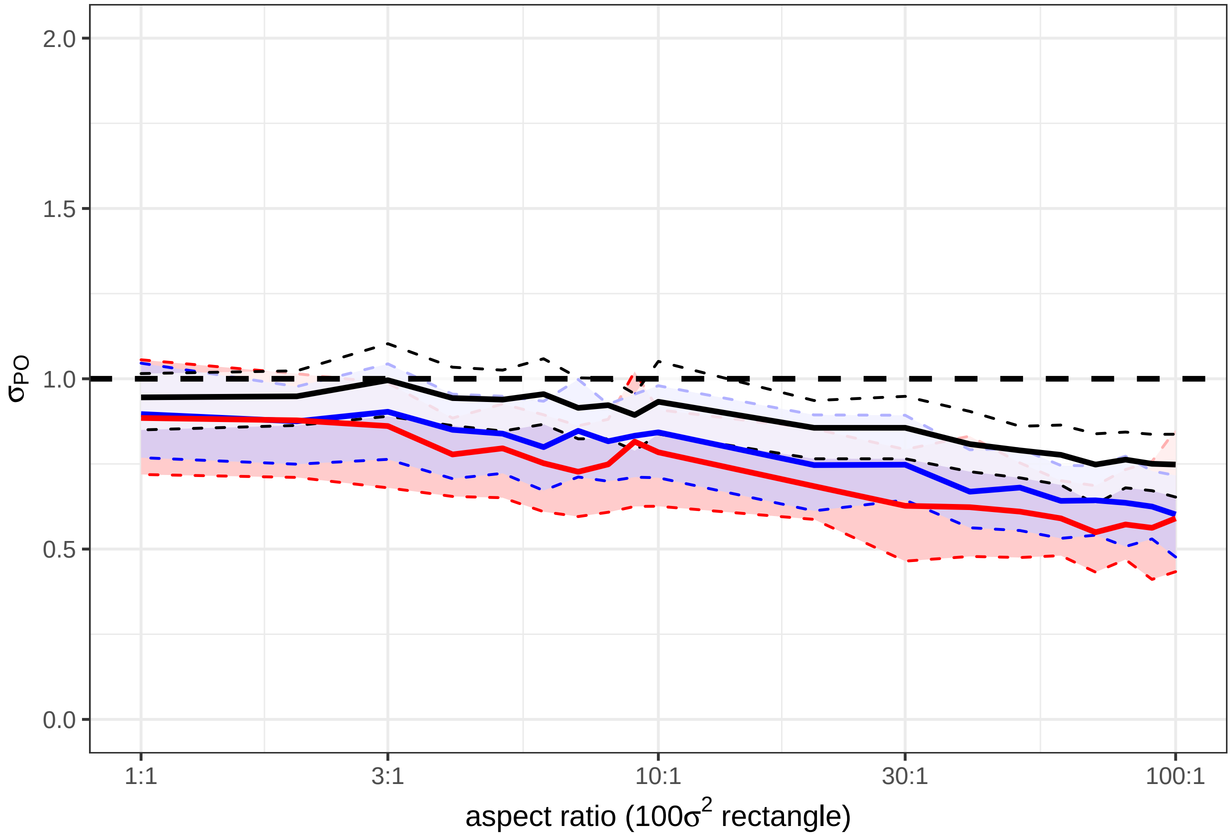

##### Number of sampled kin dyads

Figure 3q-t shows the impact of number of kin dyads sampled on the confidence intervals of an estimate of axial $\sigma$

```
nval <- 1:50 * 5

numbercheck <- tibble(n = 0, l = 0, m = 0, u = 0, ll = 0, ml = 0, ul = 0, lg = 0, mg = 0,
                      ug = 0, .rows = 0)

for (n in nval){
  egauss <- simbase_gauss %>%
    axpermute(nreps = permutations, nsamp = n, composite = 1) %>% as.vector()
  elap <- simbase_laplace %>%
    axpermute(nreps = permutations, nsamp = n, composite = 1) %>% as.vector()
  egam <- simbase_vgamma %>%
    axpermute(nreps = permutations, nsamp = n, composite = 1) %>% as.vector()
  numbercheck <- numbercheck %>%
    add_row(n = n, l = egauss[1], m = egauss[2], u = egauss[3],
            ll = elap[1], ml = elap[2], ul = elap[3], lg = egam[1], mg = egam[2],
            ug = egam[3])
}
```

```
ggplot(numbercheck) + aes(x = n) +
  geom_ribbon(mapping = aes(ymin = lg, ymax = ug), fill = "#ffcccc", colour = "red",
              linetype = 2)+
  geom_ribbon(mapping = aes(ymin = ll, ymax = ul), fill = "#ccccff", alpha = 0.7,
              colour = "blue", linetype = 2)+
  geom_ribbon(mapping = aes(ymin = l, ymax = u), fill = "#ffffff", alpha = 0.7,
              colour = "black", linetype = 2) +
  geom_hline(yintercept = 1, linetype = 2, size = 1) +
  theme_bw() +
  coord_cartesian(ylim = c(0, 2)) +
  xlab("number sampled") + ylab(expression(sigma["PO"])) #+
```

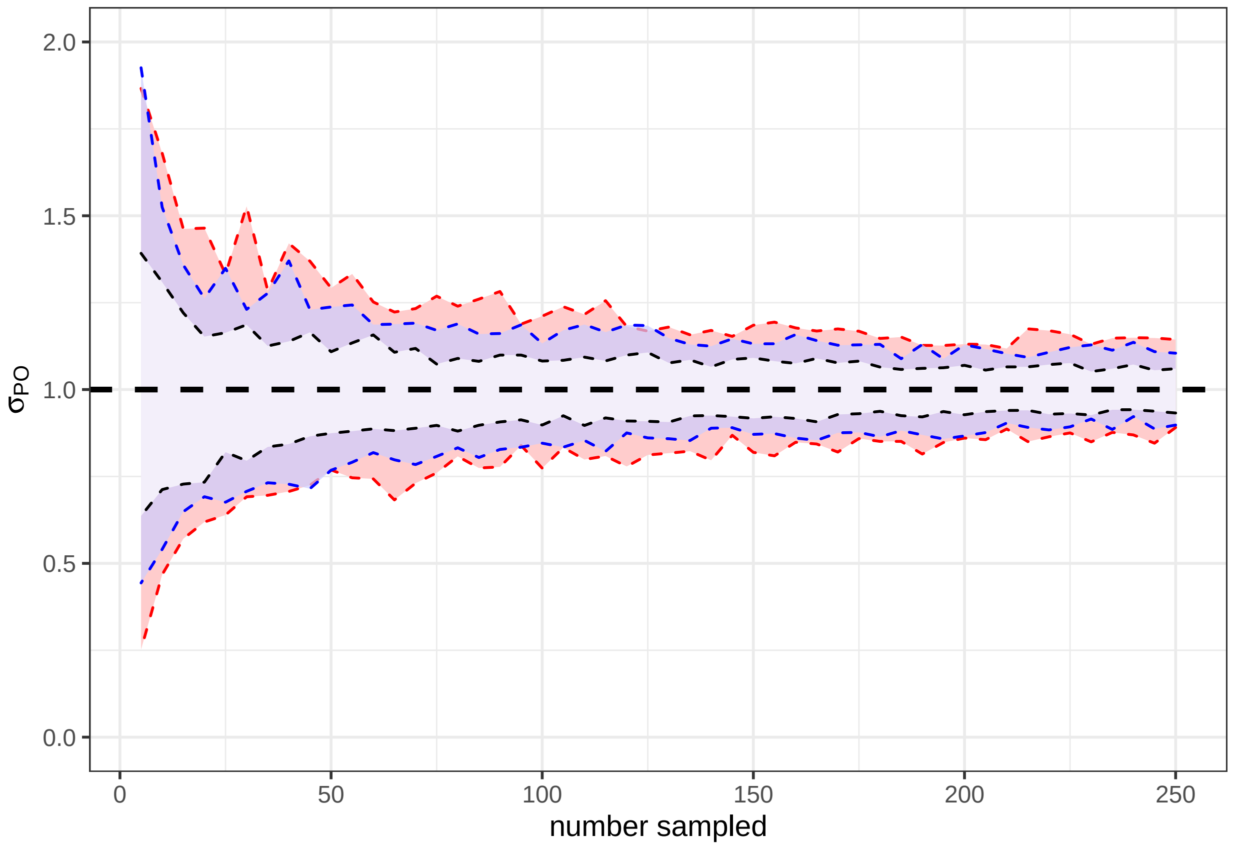
