## Supplementary Text 2 for "Estimating dispersal using close kin dyads: The kindisperse R package"

### Supplementary Text 2. Sampling Considerations

Moshe E. Jasper, Ary A. Hoffmann, and Thomas L. Schmidt

23/07/2021

#### Contents

**1. Planning sampling site area for a close-kin dispersal estimate** **1**

Simulation A. (sampling of immatures). . . . . . . . . . . . . . . . . . . . . . . . . . . . . . . . . . 2

Simulation B. (sampling of ovipositing adults). . . . . . . . . . . . . . . . . . . . . . . . . . . . . . 4

**2. Assessing bias driven by sampling site area** **7**

**3. References** **10**

```
library(tidyverse)
library(kindisperse)
```

### 1. Planning sampling site area for a close-kin dispersal estimate

How large should a sampling area be to enable an accurate estimation of $\sigma_{PO}$? This question is illuminated by the simulations run in this paper, but in practice turns out to be quite complicated, and would vary depending on assumptions about the kernel shape (Gaussian, Laplace, a variance-gamma distribution, etc.), and dependent on other aspects of sampling design.

The first rule is that the more leptokurtic (long-tailed, i.e. dominated by long-distance dispersal events) a dispersal distribution is expected to be, the larger the site dimensions need to be, to capture sufficient of this long-tailed dispersal. In the case of pure PO-sampling (were this possible) along the lines shown in Figure 4c, if we adopt the requirement that a 'good enough' estimate of $\sigma_{PO}$ will be at least 90% of the actual value, we find that while the dimensions of a Gaussian distribution need to be $5\sigma$, those of a Laplace distribution need to be ~$10\sigma$, and a variance-gamma (0.5) distribution $15\sigma$. Careful thought must be put into what is known about long-distance dispersal within a study before these methods are deployed.

However, in practice, the PO kernel cannot be sampled directly; we infer it by sampling from other kernels. A typical study in mosquitoes would involve sampling from the distributions of immature FS kin and immature 1C kin, then using these to extract the PO kernel. But as these distributions are of a different spatial scale to the PO distribution, they will also be impacted differently by spatial characteristics of the sampling area. The smaller FS kernel will be more prone to over-estimation if minimum sampling distances are too large, and the larger 1C kernel will be prone to under-estimation (leading to under-estimation of $\sigma_{PO}$). If these distributions are known, they will assist in determining an appropriate scale at which to sample. In practice, these distributions will not be known in advance, and there will be at best literature estimates of PO dispersal. In a Gaussian context, the 1C distribution is at least $\sqrt{2}$ larger than the PO distribution (as it involves two PO draws + two oviposition draws), Assuming the oviposition dispersal kernel makes up a quarter of the PO kernel (along with initial, breeding and gravid), the 1C distribution is around $1.6\sigma_{PO}$. Applying the insights of Figure 4c, a closer length for site dimensions would be $5 * 1.6\sigma_{PO} = 7.9\sigma_{PO}$; but we still must account for the FS distribution, which, being smaller, is relatively unscathed.

Through simulations (Simulation A., below) we arrive at a final value for the estimation of $\sigma_{PO}$ within a Gaussian framework to be site dimensions of size $9\sigma_{PO}$.This is the estimate given in the main text of the paper. The estimate for Laplace is $9.5\sigma_{PO}$. That for variance-gamma (0.5) is around $10\sigma_{PO}$ (see simulations below).

Note that all of the above assumes samples are collected at the immature lifestage. If samples are collected as ovipositing adults, distributions are expected to be larger, and thus sampling sites will need to be much larger (up to $14/sigma_{PO}$, as suggested by simulation b. and Supplementary Figure 2 below).

#### Simulation A. (sampling of immatures).

```
# Gaussian case
c1 <- simulate_kindist_composite(nsims = 1000000, initsigma = 0.5, breedsigma = 0.5,
                                 gravsigma = 0.5, ovisigma = 0.5, kinship = "1C",
                                 method = "Gaussian")
fs <- simulate_kindist_composite(nsims = 1000000, initsigma = 0.5, breedsigma = 0.5,
                                 gravsigma = 0.5, ovisigma = 0.5, kinship = "FS",
                                 method = "Gaussian")

gauss <- tibble(dims = 0, est = 0, .rows = 0)
for (dimfilter in 20:60/4){


c1d <- c1  %>% sample_kindist(dims = dimfilter)
fsd <- fs  %>% sample_kindist(dims = dimfilter)
est <- axials_standard(c1d, fsd)

gauss <- gauss %>% add_row(dims = dimfilter, est = est)

}

gauss_intercept <- filter(gauss, est > 0.9)[1,]
gauss_intercept$dims
```

```
## [1] 9
```

```
# Laplace case
c1 <- simulate_kindist_composite(nsims = 1000000, initsigma = 0.5, breedsigma = 0.5,
                                 gravsigma = 0.5, ovisigma = 0.5, kinship = "1C",
                                 method = "Laplace")
fs <- simulate_kindist_composite(nsims = 1000000, initsigma = 0.5, breedsigma = 0.5,
                                 gravsigma = 0.5, ovisigma = 0.5, kinship = "FS",
                                 method = "Laplace")

lap <- tibble(dims = 0, est = 0, .rows = 0)
for (dimfilter in 20:60/4){
```

```r
c1d <- c1  %>% sample_kindist(dims = dimfilter)
fsd <- fs  %>% sample_kindist(dims = dimfilter)
est <- axials_standard(c1d, fsd)

lap <- lap %>% add_row(dims = dimfilter, est = est)

}

lap_intercept <- filter(lap, est > 0.9)[1,]
lap_intercept$dims
```

```
## [1] 9.5
```

```r
# vgamma 0.5 case
c1 <- simulate_kindist_composite(nsims = 1000000, initsigma = 0.5, breedsigma = 0.5,
                                 gravsigma = 0.5, ovisigma = 0.5, kinship = "1C",
                                 method = "vgamma", shape = 0.5)
fs <- simulate_kindist_composite(nsims = 1000000, initsigma = 0.5, breedsigma = 0.5,
                                 gravsigma = 0.5, ovisigma = 0.5, kinship = "FS",
                                 method = "vgamma", shape = 0.5)

gam <- tibble(dims = 0, est = 0, .rows = 0)
for (dimfilter in 20:60/4){


c1d <- c1  %>% sample_kindist(dims = dimfilter)
fsd <- fs  %>% sample_kindist(dims = dimfilter)
est <- axials_standard(c1d, fsd)

gam <- gam %>% add_row(dims = dimfilter, est = est)

}

gam_intercept <- filter(gam, est > 0.9)[1,]
gam_intercept$dims
```

```
## [1] 10.25
```

**Supplementary Figure 1. Site dimensions for sigma PO estimation (immature life-stage)**

```r
ggplot(gam) + aes(x = dims, y = est) +
  geom_line(mapping = aes(colour = "variance-gamma 0.5"), size = 1) +
  geom_line(data = lap, mapping = aes(colour = "Laplace"), size = 1)+
  geom_line(data = gauss, mapping = aes(colour = "Gaussian"), size = 1)+
  geom_hline(yintercept = 0.9, linetype = 2) +
  geom_segment(mapping = aes(x = gauss_intercept[[1]], y = gauss_intercept[[2]],
                             xend = gauss_intercept[[1]], yend = 0.85), linetype = 2)+
  geom_segment(mapping = aes(x = lap_intercept[[1]], y = lap_intercept[[2]],
                             xend = lap_intercept[[1]], yend = 0.85), linetype = 2,
               colour = "blue")+
```

```
  geom_segment(mapping = aes(x = gam_intercept[[1]], y = gam_intercept[[2]],
                             xend = gam_intercept[[1]], yend = 0.85), linetype = 2,
             colour = "red")+
  scale_x_continuous(breaks = c(8, 8.5, 9, 9.5, 10, 10.5, 11, 11.5, 12))+
  xlab("Site dimensions") + ylab(expression(paste(sigma["PO"], " estimate")))+
  coord_cartesian(ylim = c(0.85, 0.95), xlim = c(8, 12))+
  scale_colour_manual(breaks = c("Gaussian", "Laplace", "variance-gamma 0.5"),
                      values = c("black", "blue", "red"),
                      name = "Sampling distribution")+
  theme_bw()
```

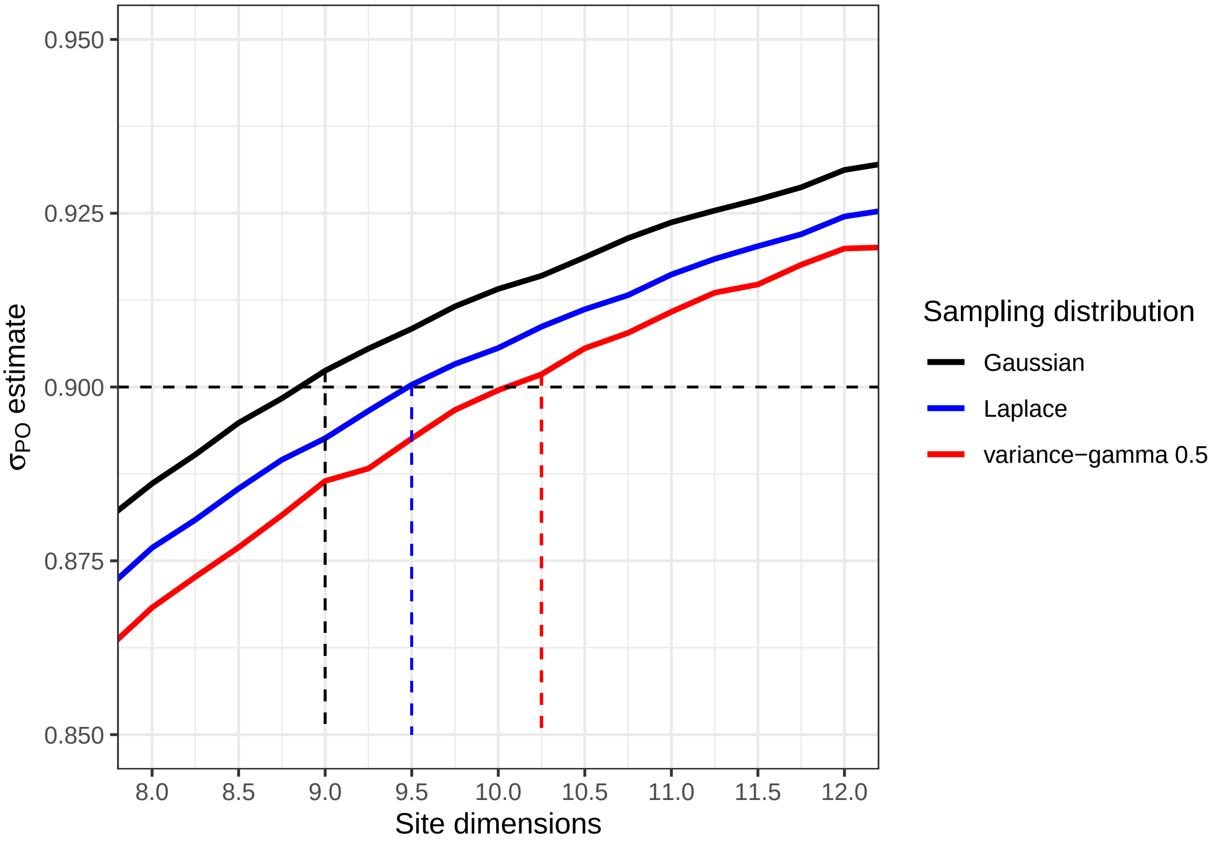


From this we can see that under Gaussian assumptions, site dimensions should be closer to $9\sigma$ (this Figure is used in the paper). The dimensions should be $9.5\sigma$ for Laplace, and $10\sigma$ for variance-gamma 0.5.

##### Simulation B. (sampling of ovipositing adults).

```
# Gaussian case
c1 <- simulate_kindist_composite(nsims = 1000000, initsigma = 0.5, breedsigma = 0.5,
                                 gravsigma = 0.5, ovisigma = 0.5, kinship = "1C",
                                 method = "Gaussian", lifestage = "ovipositional")
fs <- simulate_kindist_composite(nsims = 1000000, initsigma = 0.5, breedsigma = 0.5,
                                 gravsigma = 0.5, ovisigma = 0.5, kinship = "FS",
                                 method = "Gaussian", lifestage = "ovipositional")
```

```
gauss <- tibble(dims = 0, est = 0, .rows = 0)
for (dimfilter in 20:60/4){


c1d <- c1  %>% sample_kindist(dims = dimfilter)
fsd <- fs  %>% sample_kindist(dims = dimfilter)
est <- axials_standard(c1d, fsd)

gauss <- gauss %>% add_row(dims = dimfilter, est = est)

}

gauss_intercept <- filter(gauss, est > 0.9)[1,]
gauss_intercept$dims
```

```
## [1] 14
```

```
# Laplace case
c1 <- simulate_kindist_composite(nsims = 1000000, initsigma = 0.5, breedsigma = 0.5,
                                 gravsigma = 0.5, ovisigma = 0.5, kinship = "1C",
                                 method = "Laplace", lifestage = "ovipositional")
fs <- simulate_kindist_composite(nsims = 1000000, initsigma = 0.5, breedsigma = 0.5,
                                 gravsigma = 0.5, ovisigma = 0.5, kinship = "FS",
                                 method = "Laplace", lifestage = "ovipositional")

lap <- tibble(dims = 0, est = 0, .rows = 0)
for (dimfilter in 20:60/4){


c1d <- c1  %>% sample_kindist(dims = dimfilter)
fsd <- fs  %>% sample_kindist(dims = dimfilter)
est <- axials_standard(c1d, fsd)

lap <- lap %>% add_row(dims = dimfilter, est = est)

}

lap_intercept <- filter(lap, est > 0.9)[1,]
lap_intercept$dims
```

```
## [1] 14.5
```

```
# variance-gamma 0.5 case
c1 <- simulate_kindist_composite(nsims = 1000000, initsigma = 0.5, breedsigma = 0.5,
                                 gravsigma = 0.5, ovisigma = 0.5, kinship = "1C",
                                 method = "vgamma", shape = 0.5,
                                 lifestage = "ovipositional")
fs <- simulate_kindist_composite(nsims = 1000000, initsigma = 0.5, breedsigma = 0.5,
                                 gravsigma = 0.5, ovisigma = 0.5, kinship = "FS",
                                 method = "vgamma", shape = 0.5,
                                 lifestage = "ovipositional")
```

```r
gam <- tibble(dims = 0, est = 0, .rows = 0)
for (dimfilter in 20:60/4){


c1d <- c1  %>% sample_kindist(dims = dimfilter)
fsd <- fs  %>% sample_kindist(dims = dimfilter)
est <- axials_standard(c1d, fsd)

gam <- gam %>% add_row(dims = dimfilter, est = est)

}

gam_intercept <- filter(gam, est > 0.9)[1,]
gam_intercept$dims
```

```
## [1] 14.25
```

**Supplementary Figure 2. Site dimensions for sigma PO estimation (ovipositing adults)**

```r
ggplot(gam) + aes(x = dims, y = est) +
  geom_line(mapping = aes(colour = "variance-gamma 0.5"), size = 1) +
  geom_line(data = lap, mapping = aes(colour = "Laplace"), size = 1)+
  geom_line(data = gauss, mapping = aes(colour = "Gaussian"), size = 1)+
  geom_hline(yintercept = 0.9, linetype = 2) +
  geom_segment(mapping = aes(x = gauss_intercept[[1]], y = gauss_intercept[[2]],
                             xend = gauss_intercept[[1]], yend = 0.85), linetype = 2)+
  geom_segment(mapping = aes(x = lap_intercept[[1]], y = lap_intercept[[2]],
                             xend = lap_intercept[[1]], yend = 0.85), linetype = 2,
               colour = "blue")+
  geom_segment(mapping = aes(x = gam_intercept[[1]], y = gam_intercept[[2]],
                             xend = gam_intercept[[1]], yend = 0.85), linetype = 2,
               colour = "red")+
  scale_x_continuous(breaks = c(12, 12.5, 13, 13.5, 14, 14.5, 15, 15.5, 16))+
  xlab("Site dimensions") + ylab(expression(paste(sigma["PO"], " estimate")))+
  coord_cartesian(ylim = c(0.875, 0.925), xlim = c(12, 15))+
  scale_colour_manual(breaks = c("Gaussian", "Laplace", "variance-gamma 0.5"),
                      values = c("black", "blue", "red"), name = "Sampling distribution")+
  theme_bw()
```

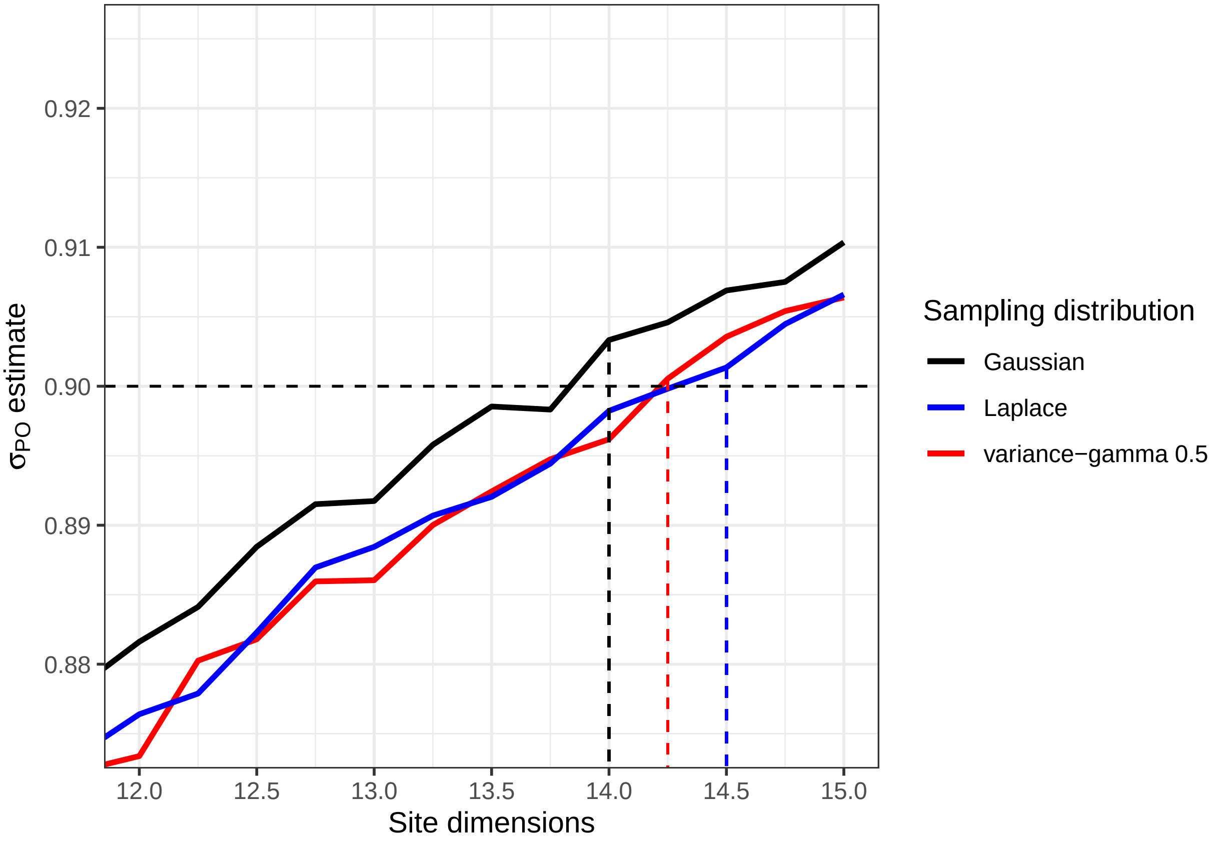


All ovipositional estimate dimension sizes lie around $14\sigma_{PO}$

#### 2. Assessing bias driven by sampling site area

It is important to be able to assess the impact of sample site size on estimates of PO dispersal after a study has been conducted, as well as in the planning phase. To illustrate such a *post-hoc* estimate, we here recalculate then assess the estimates of PO sigma present in our previous paper (Jasper et al. 2019). The close-kin data from this paper is embedded within the kindisperse package as the object `mentari`:

```
mentari
```

```
## # A tibble: 98 x 10
##      id1   id2 kinship distance    x1    y1    x2    y2 lifestage k_loiselle
##    <dbl> <dbl> <chr>      <dbl> <dbl> <dbl> <dbl> <dbl> <chr>          <dbl>
##  1     1    31 1C         45.7  12.1   57.8  11.8  104.  immature      0.0684
##  2     3    17 HS          4.43  7.45  67.3  11.9   67.3 immature      0.0933
##  3     4    29 HS         49.2   7.27  73.2  11.2  122.  immature      0.0807
##  4     7     9 FS          7.18  8.51  48.8  12.4   54.8 immature      0.220
##  5     7    10 FS          9.74  8.51  48.8  12.1   57.8 immature      0.226
##  6     7   142 HS        313.    8.51  48.8 251.   247.  immature      0.0805
##  7     9    10 HS          3.04 12.4   54.8  12.1   57.8 immature      0.155
##  8    10   149 1C        199.   12.1   57.8  91.6  240.  immature      0.0620
##  9    11    21 1C         51.8   7.45  67.3   5.90 119.  immature      0.0664
## 10    13    18 FS         47.4   8.51  48.8  14.8   95.8 immature      0.243
## # ... with 88 more rows
```

This is a dataframe containing location and kin category information for all close-kin dyads identified by sampling within an area of 500m by 250m.

We first load each subcategory into custom objects:

```
m_fs <- df_to_kinpair(mentari, kinship = "FS")
m_hs <- df_to_kinpair(mentari, kinship = "HS")
m_1c <- df_to_kinpair(mentari, kinship = "1C")
```

Note that these are all identified as of the 'immature' lifestage.

Now, we perform the combined estimation of PO sigma used in Jasper et al. (2019).

```
axpermute_standard(avect = m_1c, bvect = m_fs, amix = TRUE, amixcat = "H1C",
                   bcomp = TRUE, bcompvect = m_hs)
```

```
##     2.5%     mean    97.5%
## 22.35334 45.20792 61.80878
```

The mean estimate here is a $\sigma_{PO}$ of 45.2m, half of the estimated neighbourhood radius of 91m. But is the sample site large enough to support a $\sigma_{PO}$ estimate of this size?

As a simple check, we first construct a simulation of this size with the function `simulate_kindist_simple` (we'll leave this on its default Gaussian setting):

```
s1 <- simulate_kindist_simple(nsims = 100000, sigma = 45.2, kinship = "PO")
s1
```

```
## KINDISPERSE SIMULATION of KIN PAIRS
## -----------------------------------
## simtype:      simple
## kerneltype:       Gaussian
## kinship:      PO
## simdims:      100 100
## posigma:      45.2
## lifestage:        immature
##
## tab
## # A tibble: 100,000 x 8
##    id1   id2   kinship distance    x1    y1    x2    y2
##    <chr> <chr> <chr>      <dbl> <dbl> <dbl> <dbl> <dbl>
##  1 1a    1b    PO          81.3 28.4  47.1  -45.6  13.4
##  2 2a    2b    PO         125.  59.2  51.7  -13.7 153.
##  3 3a    3b    PO          68.9 72.0  58.4  124.  104.
##  4 4a    4b    PO          24.4 27.9  95.2   50.2 105.
##  5 5a    5b    PO          56.5 14.0  75.1  -42.1  81.8
##  6 6a    6b    PO          73.7  6.65 47.9   70.1  10.4
##  7 7a    7b    PO          29.3 56.1  89.6   44.8 117.
##  8 8a    8b    PO          65.9 42.9   3.77  57.0  68.1
##  9 9a    9b    PO          46.0 60.5  43.6   33.6  81.0
## 10 10a   10b   PO          48.1 58.4  81.6   10.6  76.2
## # ... with 99,990 more rows
## -----------------------------------
```

Now, we use the function `sample_kindist` to impose our sampling site geometry on the estimate (250m by 500m) then make a basic estimate:

```
s1_samp <- sample_kindist(s1, dims = c(250, 500), n = 10000)
```

```
## Setting central sampling area to 250 by 500

## Down-sampling to 10000 kin pairs

## 10000 kin pairs remaining.
```

```
axials(s1_samp)
```

```
## [1] 42.44765
```

the estimate given is 42, a little less than the initial estimate of 45m. This is fairly close, so not much of an issue. But let's try and find a value of $\sigma_{PO}$ such that our sampling site returns the actual value we found in the study:

```
s2 <- simulate_kindist_simple(nsims = 100000, sigma = 49, kinship = "PO")
s2_samp <- sample_kindist(s2, dims = c(250, 500), n = 10000)
```

```
## Setting central sampling area to 250 by 500

## Down-sampling to 10000 kin pairs

## 10000 kin pairs remaining.
```

```
axials(s2_samp)
```

```
## [1] 45.82194
```

Assuming a Gaussian kernel, the initial sigma may well have been higher than 49m. However, let's now look at a varance-gamma (0.5) kernel:

```
s3 <- simulate_kindist_simple(nsims = 100000, sigma = 49, kinship = "PO",
                              method = "vgamma", shape = 0.5)
s3_samp <- sample_kindist(s3, dims = c(250, 500), n = 10000)
```

```
## Setting central sampling area to 250 by 500

## Down-sampling to 10000 kin pairs

## 10000 kin pairs remaining.
```

```
axials(s3_samp)
```

```
## [1] 38.74242
```

Under a stronger variance-gamma kernel (i.e. assuming a significant amount of long-distance dispersal), a site $\sigma_{PO}$ of 49m completely failed to produce the estimate of sigma we arrived at in this study. Let's try again.

```r
s4 <- simulate_kindist_simple(nsims = 100000, sigma = 60, kinship = "PO",
                              method = "vgamma", shape = 0.5)
s4_samp <- sample_kindist(s4, dims = c(250, 500), n = 10000)
```

```
## Setting central sampling area to 250 by 500
```

```
## Down-sampling to 10000 kin pairs
```

```
## 10000 kin pairs remaining.
```

```r
axials(s4_samp)
```

```
## [1] 44.11594
```

It seems in this more leptokurtic scenario the underlying $\sigma_{PO}$ could have been as high as 60m. Following the recommendations set in the previous section, if substantial long-distance dispersal was expected, the ideal site dimensions for future *Ae. aegypti* dispersal studies in a similar context would need to be $10\sigma$x$10\sigma$ i.e. at least 600m by 600m.
