## Supplementary Text 3 for "Estimating dispersal using close kin dyads: The kindisperse R package"

### Supplementary Text 3. KINDISPERSE 0.10.1 README

Moshe E. Jasper, Ary A. Hoffmann, and Thomas L. Schmidt

23/07/2021

**5. References** **22**

The goal of KINDISPERSE is to simulate and estimate close-kin dispersal kernels.

### 1. Introduction

Dispersal is a key evolutionary process that connects organisms in space and time. Assessing the dispersal of organisms within an area is an important component of estimating risks from invasive species, planning pest management operations, and evaluating conservation strategies for threatened species.

Leveraging decreases in sequencing costs, out new method instead estimates dispersal from the spatial distribution of close-kin. This method requires that close kin dyads be identified and scored for two variables: (i) the geographical distance between the two individuals in the dyad, and (ii) their estimated order of kinship (1st order e.g. full-sib; 2nd order e.g. half-sib; 3rd order e.g. first cousin).

Close-kin-based dispersal can provide an estimate of the intergenerational (or parent-offspring) dispersal kernel - a key factor that connects biological events across the lifespan of an organism with broader demographic and population-genetic processes such as isolation by distance. A dispersal kernel is the probability density function describing the distributions of the locations of dispersed individuals relative to the source point. Intergenerational dispersal kernels themselves can be framed in terms of any number of breeding and dispersal processes, defined by both reference life-stage and number of generations, and leave their mark in the spatial distribution of various categories of close kin, which can be treated as samplings from a set of underlying kernels. Actual kernels vary, but are typically described in terms of sigma, the second moment of the kernel, also known as its scale parameter. More complex kernels can also incorporate a parameter for shape or kurtosis (kappa), representing the fourth moment of the kernel.

In the case of an insect like the mosquito, the most basic intergenerational kernel, the lifespan or parent-offspring kernel, reflects all dispersal and breeding processes connecting e.g. the immature (egg, larval, pupal) location of a parent to the immature location of its offspring. However, this kernel can be combined with additional breeding, dispersal and sampling events to produce other, composite dispersal or distribution kernels that contain information about intergenerational dispersal. For example, the distribution of two immature full-sibling mosquitoes reflects not a full lifespan of dispersal, but two 'draws' from the component kernel associated with the mother's ovipositing behaviour. Were we to sample the same full-sibling females as ovipositing adults, this would instead represent two draws from a composite 'lifespan and additional oviposition' kernel. Avuncular larvae, should they exist, would represent draws from related but distinct intergenerational dispersal kernels - an oviposition kernel, and a composite 'oviposition and lifespan' kernel. The avuncular distribution kernel would thus reflect a further compositing of these dispersal events.

There is a rich literature examining the kernels of basic dispersal events, and analysing them in terms of various kernel functions, whether Gaussian, exponential, or others with differing properties and shapes, often reflecting the tendency of dispersal events to be disproportionately clustered around the source and/or be dispersed at great distances from the source (i.e. for the kernel to be fat-tailed). Most of this literature explores dispersal in terms of probability of a dispersed sample being at a certain radius from the dispersed source. In the case of close-kin recaptures of e.g. first cousins, we are instead presented with dispersal events that must be approached in two dimensions with respect to both radius of dispersal and additionally angle of dispersal. A successful estimator of intergenerational dispersal using close-kin recaptures must find strategies to decompose the extraneous spatial and breeding components affecting the kernels, and ultimately re-express dispersal in terms of an axial sigma - that aspect of dispersal which operates within one dimension across a two-dimensional space. This is the sigma component relied upon by Wright (1946) for isolation by distance , and which is reflected in estimations of neighbourhood area.

The method we have developed relies upon the fact that different kinship categories reflect different but related underlying intergenerational dispersal composites, and uses the relationships between these kinship distribution kernels to extract information about the core parent-offspring dispersal kernel that produced the derivative kernels. For example, the immature distribution kernel of full siblings differs from the immature distribution of first cousins by a single lifespan: using an additive variance framework, the first cousin variance that is not accounted for by subtracting the full sibling variance constitutes an estimate of the parent-offspring distribution, from which an intergenerational kernel estimate can be derived. This is because both immature full-sibling and immature first cousins are 'phased' with respect to the organisms' life cycle - that is, they are separated by an integer multiple of parent-offspring dispersal events. It is this phasing that enables the extraction of a 'pure' effective dispersal estimate, via the additive property of variance. Other examples of phased relationships include half sibling immatures to half cousin immatures (one cycle), full sibling immatures to second cousin immatures (two cycles), or even (for mosquitoes) full sibling immatures to second cousin ovipositing adults (three cycles).

Further details can be found in Jasper et al. (2019), "A genomic approach to inferring kinship reveals limited intergenerational dispersal in the yellow fever mosquito."

This package supplements these papers by supplying methods for (a) importing and exporting information about distances and kinship relationships for dyads of individuals, (b) estimating the axial distribution (axial sigma for dispersal or position distributions) from empirical distributions of kin-dyads, and (c) estimating the intergenerational (parent-offspring) dispersal distribution (axial sigma) that underlies the distributions of multiple phased kin categories. This package also implements several simulation tools for further exploring and testing the properties of intergenerational dispersal kernels, as well as to assist in designing experiment layouts and sampling schemes. Finally, for ease of use, the package supplies an integrated shiny app which also implements the vast majority of package functionality in a user-friendly interface.

#### 2. Installation

You can install the released version of kindisperse from CRAN with:

```
install.packages("kindisperse")
```

And the development version from GitHub with:

```
# install.packages("devtools")
devtools::install_github("moshejasper/kindisperse")
```

Once installed, load the package as follows:

```
library(kindisperse)
#> kindisperse 0.10.1
```

#### 3. The shiny app

The kindisperse app bundles most of the tools supplied in this package for ease of use.

##### 3.1 Running the app

To run the app, enter the function `run_kindisperse()` and in a moment the app will appear in a separate window. To close, exit this window, or alternatively hit the 'stop' button or equivalent in RStudio.

##### 3.2 External interface

To calculate axial values, etc. of objects within the app, they first must be passed to the app from the computer or the R package environment. One option is to save objects to the computer via either .csv or .kindata formats, then load them using the in-app interface.

Alternatively, objects you have loaded or created in the R package environment can be passed to the app by first mounting them to the special `appdata` environment which can be accessed from within the app via the `Load` tab. Mounted objects must be of class `KinPairData` or `KinPairSimulation`. To mount an object, use the `mount_appdata(obj, "nm")` function (unmount with `unmount_appdata("nm")`). The `appdata` environment can be viewed with `display_appdata()` and cleared with `reset_appdata()`. Objects mounted to appdata from within the app can also be retrieved with `retrieve_appdata()` or `retrieveall_appdata()`.

```
fullsibs <- simulate_kindist_composite(nsims = 100, ovisigma = 25, kinship = "FS")
reset_appdata()
mount_appdata(fullsibs, "fullsibs")
display_appdata()
#> <environment: kindisperse_appdata>
#> parent: <environment: namespace:kindisperse>
#> bindings:
#>  * fullsibs: <KnPrSmlt>
fullsibs2 <- retrieve_appdata("fullsibs")
reset_appdata()
```

The app also uses a temporary environment for in-app data handling and storage. Following a session, objects stored in this space can be bulk-accessed via the function `retrieve_tempdata()`, and reset via the function `reset_tempdata()`.

### 4. The R package

Package functions and typical usage are introduced below

#### 4.1 Simulations and Sampling

KINDISPERSE is is built to run three types of simulation. The first is a graphical simulation showing the dispersal of close kin over several generations. The second and third are both simulations of close kin dyads, one using a simple PO kernel, the other a composite one. The package also includes one subsampling function to assist in using these simulations for field study design.

##### 4.1.1 Graphical simulations

This is designed primarily for introducing, exploring, and easily visualising dispersal concepts. It is packaged in two parallel functions: the simulation function (`simgraph_data()`) and the visualisation function(`simgraph_graph()`). A standard example of their use is shown below, in this case, modeling families including first cousins with a kernel sigma of 25m, and site dimensions of 250x250m. The first graph shows the dispersal events leading to first cousins within five of these families.

```
## run graphical simulation
graphdata <- simgraph_data(nsims = 1000, posigma = 25, dims = 250)
simgraph_graph(graphdata, nsim = 5, kinship = "1C")
```

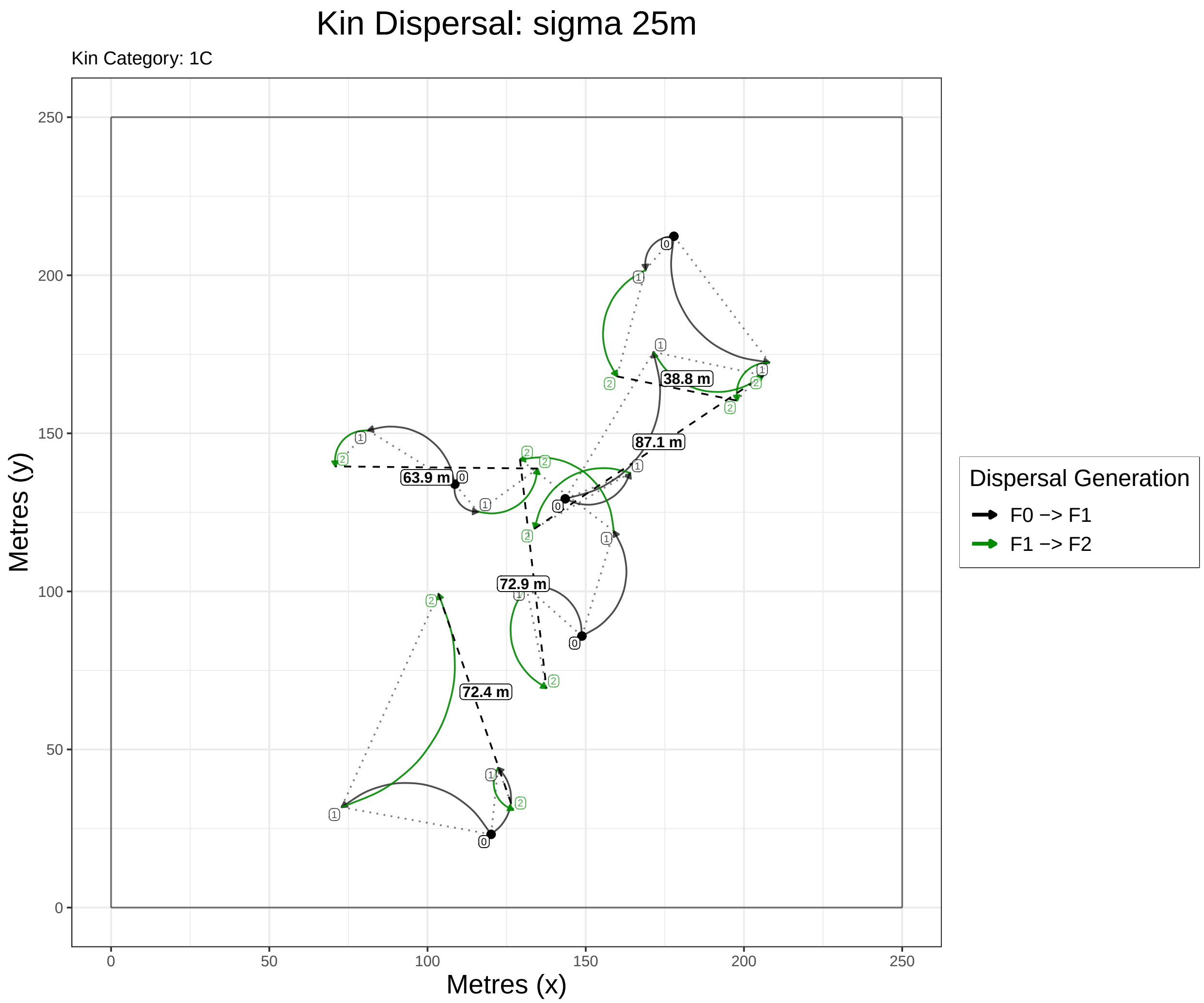

However, the options of both can be tweaked to show other data types, e.g. a pinwheel graph focused on 1,000 first cousin dyads.

```
graphdata <- simgraph_data(nsims = 1000, posigma = 25, dims = 250)
simgraph_graph(graphdata, nsims = 1000, pinwheel = T, kinship = "1C")
```

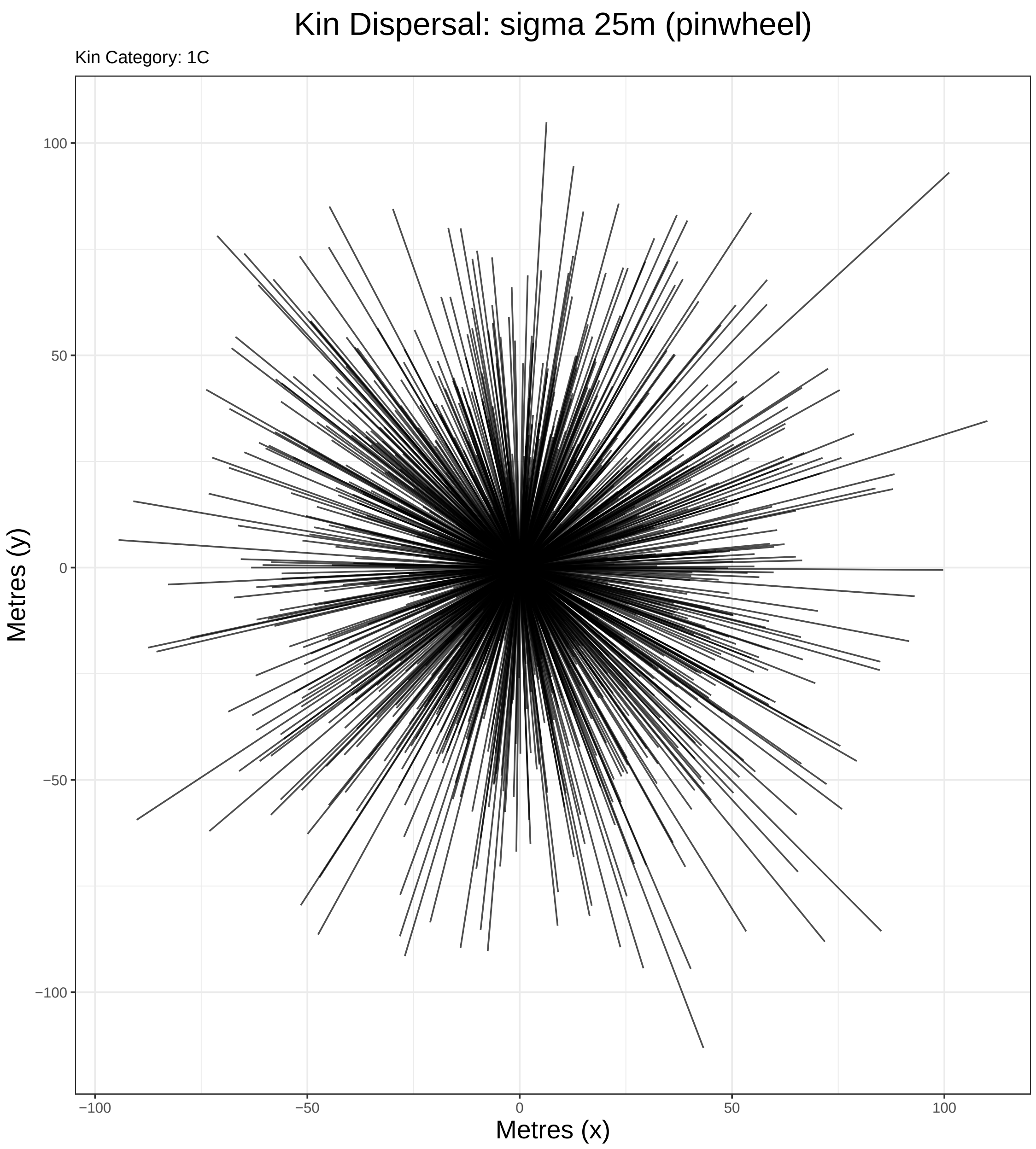

or a histogram of first cousin dyads:

```
graphdata <- simgraph_data(nsims = 1000, posigma = 25, dims = 250)
simgraph_graph(graphdata, nsims = 1000, histogram = T, kinship = "1C")
```

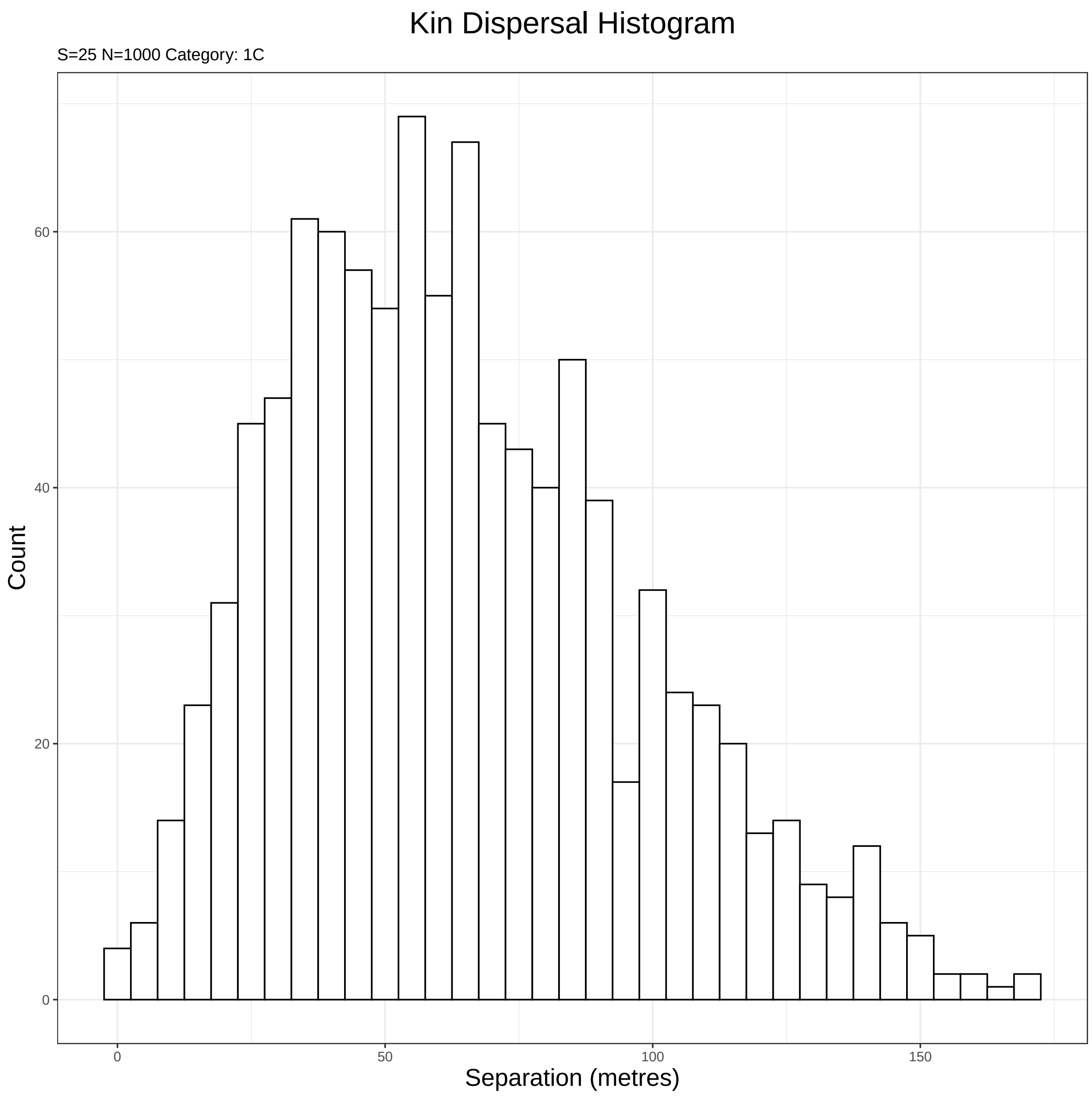

The simgraph functions are also implemented in the 'Tutorial' tab of the kindisperse app.

###### 4.1.2 Kinpair Simulations

These are designed for simulating and testing the impacts of various dispersal and sampling parameters on a dataset, and for testing and validating the estimation functions. They return an object of class `KinPairSimulation`, which supplies a tibble (dataframe) of simulation results, as well as metadata capturing the simulation parameters.

Three kernel types are supported for the next two simulations at present: `Gaussian`, `Laplace`, and `vgamma` (variance-gamma). These are passed to the functions with the `method` parameter. If using vgamma, also supply its shape parameter with the argument `shape`. Small values of `shape` correspond to an increasingly leptokurtic kernel - i.e. a strong central clustering with an increased number of very widely spaced individuals (long tails).

The simple simulation, `simulate_kindist_simple()`, simulates intergenerational dispersal for each kin category based on a simple parent-offspring dispersal sigma, with no attempt to distinguish between the various breeding and dispersal events across a lifespan. For this reason, it cannot distinguish between full and half siblings (for example), as immature full and half siblings have been separated by less than a lifespan's worth of dispersal; it would render them as at distance 0 from their parent (if they were in the adult oviposition stage, however, they would be rendered as at one lifespans' dispersal from parents).

Example usage is shown below:

```
simulate_kindist_simple(nsims = 5, sigma = 100, method = "Gaussian", kinship = "PO", lifestage = "immatu
#> KINDISPERSE SIMULATION of KIN PAIRS
#> -----------------------------------
#> simtype:      simple
#> kerneltype:       Gaussian
#> kinship:      PO
#> simdims:      100 100
#> posigma:      100
#> lifestage:        immature
#>
#> tab
#> # A tibble: 5 x 8
#>   id1   id2   kinship distance    x1    y1    x2    y2
#>   <chr> <chr> <chr>      <dbl> <dbl> <dbl> <dbl> <dbl>
#> 1 1a    1b    PO         248.   85.3 64.5   91.2 312.
#> 2 2a    2b    PO         131.   87.5 96.3  194.  173.
#> 3 3a    3b    PO         139.   33.1 72.7  -74.6 -15.1
#> 4 4a    4b    PO          78.7  87.1  4.27 164.  -11.5
#> 5 5a    5b    PO         169.   43.4 97.2   23.8 265.
#> -----------------------------------
```

The composite simulation, `simulate_kindist_composite()`, defines four smaller dispersal movements which make up the lifestage dispersal kernel. It distinguishes between full and half siblings, cousins, etc. and handles immature kin dyads that are separated by less than a lifespan of dispersal (e.g. immature FS). The four phases are 'initial' (handling any dispersal between hatching and breeding), 'breeding' (movement of the male across the breeding aspect of the cycle), 'gravid' (movement of the female after breeding but before deposition of young), and 'oviposition' (movement made while ovipositing/ bearing young). The addition of the variances of these four kernels together consitutes the lifespan dispersal kernel; the relationships between different categories inform the phase. For example, full-siblings, whether sampled at oviposition or immature states, differ in hatch position based on the ovipositing movements of the mother (including e.g. skip oviposition in the case of some mosquitoes). These categories (and any others containing a full-sib relationship buried in the pedigree) are thus of the 'full-sibling' or 'FS' phase. Half siblings, in mosquitoes (which this package is modelled on) are expected to be due to having the same father and separate mothers: the last contribution of the father's dispersal is at the breeding stage, so the 'HS' phase are differentiated by the breeding, gravid, and oviposition phases, but share in common the initial phase. The parent-offspring 'PO' phase, on the other hand, share all (or none) of the component dispersal distributions.

An example composite simulation is demostrated below:

```
simulate_kindist_composite(nsims = 5, initsigma = 50, breedsigma = 30, gravsigma = 50, ovisigma = 10, me
#> KINDISPERSE SIMULATION of KIN PAIRS
#> -----------------------------------
#> simtype:      composite
#> kerneltype:       Laplace
#> kinship:      H1C
#> simdims:      100 100
```

```
#> initsigma         50
#> breedsigma        30
#> gravsigma         50
#> ovisigma      10
#> lifestage:        ovipositional
#>
#> tab
#> # A tibble: 5 x 8
#>   id1   id2   kinship distance      x1      y1     x2     y2
#>   <chr> <chr> <chr>      <dbl>   <dbl>   <dbl>  <dbl>  <dbl>
#> 1 1a    1b    H1C        206.    -0.240   -8.96   9.46 197.
#> 2 2a    2b    H1C         70.8   12.8     85.3  -45.6   45.1
#> 3 3a    3b    H1C        333.    19.6   -160.    53.1  171.
#> 4 4a    4b    H1C        127.    82.1    121.    40.1    1.72
#> 5 5a    5b    H1C        203.   151.     105.    14.0  -45.0
#> -----------------------------------
```

Finally, a custom simulation function, `simulate_kindist_custom()` enables the simulation of dispersal in organisms with breeding cycles different to the original mosquito species this package was modeled for. These simulations take a model object (of class `DispersalModel`, generated with the function `dispersal_model()`) which contains detailed information about breeding phases, the full sibling (FS) and half sibling (HS) branch point, the sampling point, and further optional parameters defining the accessible breeding cycle more carefully. We illustrate this functions's use in the context of implementing kindisperse in a new species subsequently, but an initial example is given here. First, we generate a custom dispersal model:

```
dmodel <- dispersal_model(juvenile = 50, breeding = 40, gestation = 30, .FS = "juvenile", .HS = "breedi
dmodel
#> KINDISPERSE INTERGENERATIONAL DISPERSAL MODEL
#> ---------------------------------------------
#> stage:        juvenile   breeding    gestation
#> dispersal:    50 40  30
#>
#> FS branch:    juvenile
#> HS branch:    breeding
#> sampling stage:   gestation
#> cycle:        0 0
#> ---------------------------------------------
```

Next, we use this model to simulate dispersal in our organism:

```
simulate_kindist_custom(nsims = 5, model = dmodel, kinship = "PO")
#> KINDISPERSE SIMULATION of KIN PAIRS
#> -----------------------------------
#> simtype:      custom
#> kerneltype:       Gaussian
#> kinship:      PO
#> simdims:      100 100
#> juvenile              50
#> breeding              40
#> gestation             30
#> cycle:            0  0
#> lifestage:        gestation
#>
```

```
#> tab
#> # A tibble: 5 x 8
#>   id1   id2   kinship distance    x1    y1    x2    y2
#>   <chr> <chr> <chr>      <dbl> <dbl> <dbl> <dbl> <dbl>
#> 1 1a    1b    PO         137.   97.5 80.1  223.  135. 
#> 2 2a    2b    PO          94.8  21.6 44.9   81.1 -28.8
#> 3 3a    3b    PO         232.   65.9  6.35  71.9 239. 
#> 4 4a    4b    PO         133.   55.9 43.0   41.0 175. 
#> 5 5a    5b    PO          61.3  35.3 68.1   13.2 125. 
#> -----------------------------------
```

##### 4.1.3 Simulating Field Sampling of Kinship Distributions

This is done via another function, `sample_kindist()`, and enables the examination of how field sampling conditions could bias the estimation of axial sigma. It works with the `KinPairSimulation` or `KinPairData` classes and filters based on the study area size, number of kin expected to be found, and trap spacing. It is demonstrated below.

```
compsim <- simulate_kindist_composite(nsims = 100000, kinship = "H2C")

sample_kindist(compsim, upper = 1000, lower = 200, spacing = 50, n = 25)
#> Removing distances farther than 1000
#> Removing distances closer than 200
#> Setting trap spacing to 50
#> Down-sampling to 25 kin pairs
#> 25 kin pairs remaining.
#> KINDISPERSE SIMULATION of KIN PAIRS
#> -----------------------------------
#> simtype:      composite
#> kerneltype:       Gaussian
#> kinship:      H2C
#> simdims:      100 100
#> initsigma         100
#> breedsigma        50
#> gravsigma         50
#> ovisigma      25
#> lifestage:        immature
#>
#> FILTERED
#> --------
#> upper:            1000
#> lower:            200
#> spacing:      50
#> samplenum:        25
#>
#> tab
#> # A tibble: 25 x 8
#>    id1    id2    kinship distance    x1    y1     x2     y2
#>    <chr>  <chr>  <chr>      <dbl>  <dbl> <dbl>  <dbl>  <dbl>
#>  1 12193a 12193b H2C          375  188.  256.   -99.5   2.42
#>  2 54008a 54008b H2C          475 -141.   80.4  332.   17.4 
#>  3 49279a 49279b H2C          225 -110.   34.5   91.9  148.
```

```
#>  4 61235a 61235b H2C          725   55.5 -12.0  124.   709.
#>  5 89078a 89078b H2C          325 -101.   197.  -262.   -84.3
#>  6 18208a 18208b H2C          275  148.  499.   152.   231.
#>  7 99070a 99070b H2C          275  -59.2 -36.3 -342.    43.2
#>  8 57154a 57154b H2C          325 -269.   -49.6   43.1 -180.
#>  9 82782a 82782b H2C          425   75.3 138.   103.  -293.
#> 10 48382a 48382b H2C          375  -11.6 246.    84.3 -119.
#> # ... with 15 more rows
#> -----------------------------------
```

#### 4.2 Data Management

##### 4.2.1 Reading and writing files

Files can be loaded and saved to and from three separate formats: .csv and .tsv (via functions `csv_to_kinpair()`, `tsv_to_kinpair()`, or to save, `kinpair_to_csv()` and `kinpair_to_tsv()`, as well as the package-specific .kindata format which wraps an rds file storing package objects (via functions `read_kindata()` and `write_kindata()`). These files read to or save from an object of class `KinPairData` (including simulation objects of class `KinPairSimulation`).

.csv or equivalent files used should have a single column with the header 'distance' that contains the geographical distances between kin dyads, and preferably another column labelled 'kinship' which carries the kinship category in a form recognized by this package (see documentation for further details). Example below:

```
kinobject <- simulate_kindist_simple(nsims = 25, kinship = "FS", lifestage = "immature")
#kinpair_to_csv(kinobject, "FS_kin.csv") # saves file
#csv_to_kinpair("FS_kin.csv") # reloads it
```

##### 4.2.2 Converting objects to KinPairData format

Within the package, there are several ways to convert measures of kin dispersal distances into the `KinPairData` format required for calculations of axial distance: `vector_to_kinpair()` which takes a vector of kinpair distances, and `df_to_kinpair()` which takes a `data.frame` or `tibble` with a similar layout to the `.csv` files mentioned earlier (column of geographical distances labelled 'distance' and optional columns of kin categories ('kinship') and lifestages ('lifestage')). Inverse function is `kinpair_to_tibble()`. See relevant documentation. Example below:

```
kinvect <- c(25, 23, 43, 26, 14, 38)

vector_to_kinpair(kinvect, kinship = "H1C", lifestage = "immature")
#> KINDISPERSE RECORD OF KIN PAIRS
#> -------------------------------
#> kinship:      H1C
#> lifestage:        immature
#> cycle:            0
#>
#> tab
#> # A tibble: 6 x 4
#>   id1   id2   kinship distance
#>   <chr> <chr> <chr>      <dbl>
#> 1 1a    1b    H1C           25
```

```
#> 2 2a     2b    H1C           23
#> 3 3a     3b    H1C           43
#> 4 4a     4b    H1C           26
#> 5 5a     5b    H1C           14
#> 6 6a     6b    H1C           38
#> --------------------------------
```

Once converted, in most cases these `KinPairData` objects can be sampled in the same way as the simulations above with the function `sample_kindata()`.

#### 4.3 Estimating dispersal

The package contains a series of functions to estimate and manipulate axial sigma values (axial distributions) of simulated and empirical close-kin distributions, as well as to leverage several such distributions of related categories to supply a bootstrapped estimate of the intergenerational dispersal kernel axial sigma.

##### 4.3.1 Basic estimation of axial sigma

Axial sigma is most simply estimated with the function `axials(x, composite = 1)`. This function estimates the axial value of a simple kernel assuming that all distances measured represent one dispersal event governed by the kernel (e.g. the distance between a parent and their offspring at the equivalent lifestage, such as both as eggs). For slightly more complex situations, such as full siblings, where the distances between them result from two or more draws from the same underlying distribution (ovipositing parent to offspring #1, ovipositing parent to offspring #2), the value of `composite` can be adjusted to reflect the number of such symmetrical component events (for this specific case, you can also use `axials_norm()`). (e.g. the great-grandparent to great-grandchild category, 'GGG' is a combination of three draws from the PO distribution, and thus would take `composite = 3`):

```
paroff <- simulate_kindist_simple(nsims = 1000, sigma = 75, kinship = "PO")
axials(paroff)
#> [1] 74.85109
```

```
fullsibs <- simulate_kindist_composite(nsims = 10000, ovisigma = 25, kinship = "FS")
axials(fullsibs, composite = 2)
#> [1] 24.94615
```

Various auxillary functions exist to further manipulate axial distances within an additive variance framework, enabling the stepwise combination or averaging or decomposition of axial sigma values representing different distributions. These include `axials_decompose()` (divides into component parts as in the composite option above), `axials_add()` (adds two distributions together, e.g. FS + FS + PO + PO = 1C), `axials_combine()` (mixes two distributions together equally, e.g. 1C and H1C becomes the distribution of an even mix of both), and `axials_subtract()` subtracts a smaller distribution from a greater distribution to find the residual distribution (e.g. GG - PO = PO; FS(immature) - FS(ovipositional) = PO). For confidence intervals, there are also the permuting functions `axpermute()` and `axpermute_subtract()`.

```
axials_subtract(24, 19)
#> [1] 14.66288
```

##### 4.3.2 Estimation of axial sigma of intergenerational (PO) dispersal

Building on the above functions, the final estimation function `axials_standard()` and its permuted implementation `axpermute_standard()` take information about dispersal information across several phased categories and use it to make an estimate of the core, parent-offspring dispersal kernel (defined by axial sigma). Using this equation requires knowing representative spatial distributions of at least two **phased** kinship classes that are separated by at least one complete lifespan. In some cases, this phased requirement can be met by compositing two known distributions to approximate the distribution of a mixed category (e.g. mixing FS and HS categories to create a composited category that can be compared to an undistinguishe mixture of 1C and H1C individuals).

The function works by subtracting out the phased component of the distributions (e.g. the additional oviposition present in FS and 1C) leaving the residual lifespan components, then decomposing these down to a single span. When bootstrapped as in the `axpermute_standard()` function, these equations output the 95% confidence intervals of the resulting PO sigma estimate, as well as the estimate of median sigma. This estimate is the same sigma that interacts with Wright's neighbourhood size (the radius of NS is equal to 2x the axial sigma estimate).

Let's try out some simulated values see the function in action. First, we'll set up our individual axial sigmas for the component distributions.

```
# set up initial sigma values

init = 50
brd = 25
grv = 75
ovs = 10

# calculate theoretical PO value
po_sigma <- sqrt(init^2 + brd^2 + grv^2 + ovs^2)
po_sigma
#> [1] 94.07444
```

Here we have set up a baseline of the theoretical value of the intergenerational kernel (axial) sigma for comparison below.

First, a simple example (full sibs and first cousins) - note that the larger value must be inputted first, i.e. as `avect` in the equation. Because they are simulated objects, categories don't need to be supplied.

```
# set up sims

fullsibs <- simulate_kindist_composite(nsims = 75, initsigma = init, breedsigma = brd, gravsigma = grv,

fullcous <- simulate_kindist_composite(nsims = 75, initsigma = init, breedsigma = brd, gravsigma = grv,

# calculate PO axial sigma C.I.

axpermute_standard(fullcous, fullsibs)
#>      2.5%      mean     97.5%
#>  82.13327  92.62105 103.33133
```

As we can see, the C.I. neatly brackets the actual axial value, though with fairly large wings due to the small sample size. Now we set up a more complex case, involving a mixture of full and half cousins and a compensating compositing of full and half siblings (this will involve some data-wrangling):

```
# Set up new distributions
halfsibs <- simulate_kindist_composite(nsims = 75, initsigma = init, breedsigma = brd, gravsigma = grv,

halfcous <- simulate_kindist_composite(nsims = 75, initsigma = init, breedsigma = brd, gravsigma = grv,

# combine cousin distributions and recompose as object. Chaning kinship
# to standard value for unknown as I will be combining the distributions.
fc <- dplyr::mutate(kinpair_to_tibble(fullcous), kinship = "UN")
hc <- dplyr::mutate(kinpair_to_tibble(halfcous), kinship = "UN")
cc <- tibble::add_row(fc, hc)
cousins <- df_to_kinpair(cc)
cousins
#> KINDISPERSE RECORD OF KIN PAIRS
#> -------------------------------
#> kinship:      UN
#> lifestage:        immature
#> cycle:            0
#>
#> tab
#> # A tibble: 150 x 9
#>    id1   id2   kinship distance      x1     y1     x2     y2 lifestage
#>    <chr> <chr> <chr>      <dbl>   <dbl>  <dbl>  <dbl>  <dbl> <chr>
#>  1 1a    1b    UN          67.3    66.2  -17.3   14.0   -59.7 immature
#>  2 2a    2b    UN         102.    -68.5   74.8   6.90  144.  immature
#>  3 3a    3b    UN         141.   -111.   152.    22.1   107.  immature
#>  4 4a    4b    UN         195.    131.    54.3 -60.5    17.2 immature
#>  5 5a    5b    UN         166.    284.  -145.  169.    -25.0 immature
#>  6 6a    6b    UN         333.    160.    87.0 415.   -127.  immature
#>  7 7a    7b    UN         264.    -25.9  137.  212.     24.3 immature
#>  8 8a    8b    UN         223.    246.    31.5  23.6    24.5 immature
#>  9 9a    9b    UN         250.     -1.70  26.1 186.   -139.  immature
#> 10 10a   10b   UN          78.7   100.    35.6  27.3    65.1 immature
#> # ... with 140 more rows
#> -------------------------------
```

Note this is now a `KinPairData` object rather than a `KinPairSimulation`. The conversion to tibble and back has stripped the simulation class data. Now to run the estimation function, supplying missing category data:

```
# amix allows supply of additional (mixed) kin category H1C to acat 1C;
# bcomp allows supply of distribution to composite with bvect (this is done to match
# the cousin mixture in phase)
axpermute_standard(avect = cousins, acat = "1C", amix = TRUE, amixcat = "H1C", bvect = fullsibs, bcomp =
#>     2.5%     mean    97.5%
#> 71.77284 83.43440 95.30415
```

This estimate is a lot more convoluted, and not as 'spot on'- but the theoretical value of 94 is well within the confidence intervals.

#### 4.4 Adapting to a new species: *Antechinus*

Using custom dispersal simulations and parameters, we are well placed to explore what is typically involved in adapting this method and package to a species with a different life history and breeding structure to that of *Ae. aegypti* and other related species. The example chosen here is a species of *Antechinus* - a small marsupial native to Australia. Note that this example is for illustrative purposes only.

##### 4.4.1 Assemble known background information

Breeding and dispersal can be highly diverse processes between organisms - simply copying and pasting a method from one species to another without careful consideration of their differences and unique contexts is unwise. What relevant information can we find about species of *Antechinus*?

For *Antechinus*, mating takes place across a single week each year, and is promiscuous. Males only mate once, and die shortly after mating. Females live up to two years, producing two litters in that time. Each litter will likely contain offspring from multiple males (Cockburn, Scott, and Scotts 1985). In the same paper, Cockburn et al. recognize seven life history stages:

| No. | Stage | Duration |
|---|---|---|
| 1 | Pouch young | 5-7 weeks |
| 2 | Nest young | 8-10 weeks |
| 3 | Juveniles | rest of year 1 |
| 4 | Reproductives | 2 weeks |
| 5 | Mothers | 4-5 weeks gestation, then ongoing |
| 6 | 2nd year reproductives | 2 weeks |
| 7 | 2nd year mothers | 4-5 weeks gestation, then ongoing |

Pouch young exhibit obligatory attachment to the mother's teat. Nest young still feed on the teat, but are left in the nest (typically a hole in a tree) when the mother forages for food, until weaning. Juveniles describe the post-weaning, physiologically independent animals before synchronised reproduction occurs. This interval covers most of the first year. Reproductives (male and female) describe the animals within the very short mating window each year (males mate with multiple females, and vice versa). Mothers covers pregnancy, lactation (overlapping with nest and pouch young) and post-lactation (overlapping with the juvenile phase).

Natal dispersal (occuring after weaning) is strongly male-biased (Cockburn, Scott, and Scotts 1985), with males dispersing from the nest and often from the maternal home range, while females dispersal is more localised. In that time, males can disperse over hundreds of metres - in some species (e.g. *Antechnius stuartii*), more than a kilometre (Banks and Lindenmayer 2014). Female dispersal is not frequently beyond 50 metres (Fisher 2005).

##### 4.4.2 Identify useful life stages and kinship categories

Our key research questions will drive which aspects of the above life history we want to focus on further. For this exercise, we want to be able to estimate parent-offspring dispersal so that we can gain an estimate of the neighbourhood area. Importantly, we need this estimate to get around the sex-biased disersal in this species.

Let's define a life cycle. Pouch and nest young are still completely dependent on the mother, so will show no independent dispersal. We start our description of a single intergenerational breeding cycle with the juvenile stage, followed by breeding. We will break down the 'mother' lifestage into 'gestation' and 'pouch.'

So, our basic breeding cycle will be something like: juvenile –> breeding –> gestation –> pouch.

What kinship categories do we expect to see in *Antechinus* populations? Let's break this down by order:

**First order kinship** Within this category we have the `PO` and `FS` classes. Full sibs share the same mother, and the fathers only mate during one breeding cycle, so we can expect all full sibs to be part of the same cohort, and FS phased disperal to begin in the juvenile phase, as offspring leave the nest.

**Second order kinship** The HS kinship class can be generated by a male mating with multiple females, or a female mating with multiple males. Both of these have different dispersal modes (the former is shaped by breeding dispersal, the latter in a similar manner to the FS class). For our initial simulation, we will only treat the former kind of HS dispersal. Note that as females bear young over two generations, a third class of HS is possible, between an adult female mother and the pouch young of her (now 2nd yr) mother - cases like this are readily distinguishable by other life history traits, e.g. age.

The GG kinship class (between 2nd yr mother and her grandchildren)

The AV kinship class (between an adult female and the offspring of her full sibling - a partially sex-biased category)

**Third order kinship** Important categories here include 1C and HAV. (GAV is also possible, but clearly distinguishable by life history).

1C - first cousins. Part of same generational cohort and the FS dispersal phase.

HAV - half avuncular. Most will be intergenerational (females of previous generation to males and females of present generation). However, because females breed across two cycles, this category can occur within the same generational cohort (see below).

1C individuals will be part of the same generational cohort and result from parent-offspring dispersal events, making them a prime target (along with the FS category) for developing an intergenerational dispersal estimate.

##### 4.4.3 Build a dispersal model

Armed with the above categories, we are well placed to put together a rudimentary model of *Antechinus*. We will assign dispersal parameters to approximately reflect what we consider important in the above. As we are focusing on intergenerational dispersal in general, for now, we will ignore sex-biased aspects of dispersal (though we will take them into account when planning sampling).

```
antechinus_model <- dispersal_model(juvenile = 100, breeding = 50, gestation = 25, pouch = 25, .FS = "ju
antechinus_model
#> KINDISPERSE INTERGENERATIONAL DISPERSAL MODEL
#> ---------------------------------------------
#> stage:         breeding   gestation   pouch   juvenile
#> dispersal:     50 25   25   100
#>
#> FS branch:     juvenile
#> HS branch:     breeding
#> sampling stage:   juvenile
#> cycle:         0 0
#> ---------------------------------------------
```

Note that at this stage we have initially set sampling stage to juvenile i.e. sampling free-living individuals before their first breeding season. We will return to this later.

##### 4.4.4 Build a custom dispersal simulation

We begin with a simple PO simulation.

```
library(magrittr)
ant_po <- simulate_kindist_custom(nsims = 10000, model = antechinus_model, kinship = "PO")
ant_po
#> KINDISPERSE SIMULATION of KIN PAIRS
#> -----------------------------------
#> simtype:      custom
#> kerneltype:       Gaussian
#> kinship:      PO
#> simdims:      100 100
#> breeding              50
#> gestation             25
#> pouch             25
#> juvenile              100
#> cycle:            0 0
#> lifestage:        juvenile
#>
#> tab
#> # A tibble: 10,000 x 8
#>    id1   id2   kinship distance    x1    y1     x2    y2
#>    <chr> <chr> <chr>      <dbl> <dbl> <dbl>  <dbl> <dbl>
#>  1 1a    1b    PO          95.0 22.4  54.8    1.40 -37.8
#>  2 2a    2b    PO         118.  98.1  76.7   18.6  163.
#>  3 3a    3b    PO         111.  16.5  52.1  -38.7  149.
#>  4 4a    4b    PO         124.  98.2   9.02  72.6  130.
#>  5 5a    5b    PO         220.  73.9   3.45 287.    54.7
#>  6 6a    6b    PO          85.0 59.3  67.8   93.2  146.
#>  7 7a    7b    PO          89.0  1.45 95.0   28.6   10.2
#>  8 8a    8b    PO         237.  58.7  56.7  262.   179.
#>  9 9a    9b    PO          60.6 86.1  96.8   27.6  113.
#> 10 10a   10b   PO          89.9 77.6  53.0   65.3  142.
#> # ... with 9,990 more rows
#> -----------------------------------
```

Now we'll use the `axials()` function to characterise our 'default' dispersal for the model:

`axials(ant_po)`: 117.6050103

The value is around 117 the 'expected' value of PO we should get back from more complex estimation processes.

###### 4.4.5 Validate axial dispersal estimates and refine model

For a basic PO estimation, we are going to combine the FS and 1C categories:

```
ant_fs <- simulate_kindist_custom(nsims = 10000, model = antechinus_model, kinship = "FS")
ant_1c <- simulate_kindist_custom(nsims = 10000, model = antechinus_model, kinship = "1C")

axials_standard(ant_1c, ant_fs) # larger dispersal category goes first.
#> [1] 116.5609
```

The FS/1C strategy has been validated theoretically - but an important issue remains: the `HAV` category. While all males only breed during one breeding season, in some *Antechinus* species, many females breed for a second season. This means that the situation will arise where a mother bears offspring in one breeding season, and both the mother and her offspring bear young in the subsequent breeding season. As the second batch of young she bears are related to her previous litter as half-siblings, they are related to the that litter's offspring under the half-avuncular `HAV` kinship category. As `HAV` is of the same order of kinship (3rd) as `1C`, and (via this pathway) will be of the same lifestage, yet both pass through differing dispersal routes, without further information it would be impossible to use this class. Similar issues would hold for the `H1C` (half-cousin) and `1C1` (first cousin once removed) categories at the fourth order of kinship.

If we were simply sampling juvenile *Antechinus* as in our initial setup, there would be no way to correct for this ambiguity in the data. We need to include richer pedigree information to distinguish between the various classes. Instead of sampling at the juvenile stage, let's switch the focus to females with pouch young, and instead of genotyping one individual, plan to genotype all pouch young of a female:

```
antechinus_model <- dispersal_model(juvenile = 100, breeding = 50, gestation = 25, pouch = 25, .FS = "ju
                                     .sampling_stage = "pouch", .breeding_stage = "breeding", .visible_s
antechinus_model
#> KINDISPERSE INTERGENERATIONAL DISPERSAL MODEL
#> ---------------------------------------------
#> stage:        juvenile   breeding    gestation   pouch
#> dispersal:    100    50  25  25
#>
#> FS branch:    juvenile
#> HS branch:    breeding
#> sampling stage:   pouch
#> cycle:        0 0
#> ---------------------------------------------
```

Note that we have made several other previously implied model parameters explicit this time also (the default values are preserved here, but the concepts are important). Firstly, the `breeding_stage` parameter defines which stage in the breeding cycle breeding actually occurs at. This is by default anchored to the `HS` branch, but in situations where we might with to model `HS` dispersal that begins later (e.g. where offspring have multiple fathers but the same mother) - we might shift the `HS` branch to the juvenile stage, but preserve our information on breeding with this parameter. Second, the `visible_stage` parameter defines at what point in the life cycle an individual is considered 'available by default for sampling' in preference to its parent. This is by default anchored to the `FS` branch, and approximates birth, hatching, etc. in many species - but in many species (e.g. marsupials) an animal will be born, but still attached to the parent from the perspective of dispersal. In such situations, the `visible_stage` parameter describes which individual will be sampled by default at an overlapping lifestage. As in our model, `visible_stage` is anchored to the FS-branch juvenile stage, the default sampled pouch individuals will be the mothers rather than the offspring. In a simulation, the pouch offspring can be accessed by setting the breeding cycle parameter to `-1`.

##### 4.4.6 Finalize target kinship categories

By sampling at this stage, we unlock four different kinds of generational comparisons, all synced to the same life point: (1) intra-pouch relationships (i.e. between different pouch young carried by the same mother), (2) inter-pouch relationships (kinships between young carried by different females), (3) kinship between adult females, and (4) kinships between pouch young and adult females other than their mother.

All of these categories can be combined with the genotypic data to resolve pedigree information and enable more thorough calculations of breeding dispersal, via the following resolution:

(a) FS between pouch young: this is now a trivial category, as these will be measured before any substantial dispersal has occurred within this category. This 'zeroing' of the FS phase will simplify additional relationships.

(b) HS between pouch young (same pouch) – these correspond to the portion of HS dispersal that results from multiple males mating with the same female. Also a trivial category.
(c) HS between pouch young (different pouches) as these have different maternal ancestors, they will share the same paternal ancestor. This category thus supplies a HS estimate of the combined breeding, gestational and pouch phases.
(d) FS between (female) parents. As the FS phase is zeroed this category constitutes an estimate of lifespan dispersal for *Antechinus* females. However, as dispersal within this species is sex-biased, this does doesn't constitute the true intergenerational dispersal distance (for IBD, gene flow, etc.). Note that in this context all compared offspring are expected to fall into the 1C category.
(e) HS between (female) parents. These will result from a mixture of a shared male or female parent (i.e. the dispersal modes found in (b) and (c) above). This category thus contains the true HS phase in addition to a female-dispersed lifespan.
(f) 3rd order (female) parents. Depending on the species, it may be impossible to distinguish between the 1C and HAV kinships for individuals of this category.
(g) 3rd order between pouch young (different pouches). By themselves, this category will be indeterminate between the 1C and HAV categories. One approach would be to combine this category with category (f) to cancel out the composite phase and leave an estimate of PO dispersal – but as the more dispersed category are female adults, this would once again only produce an estimate of female dispersal across the breeding cycle. Is there another way?

Yes! We can check other pedigree relationships to distinguish between the 1C and HAV categories. Firstly, we compare the parents. If they are FS, their offspring are 1C and in isolation constitute an estimate of female intergenerational dispersal as in (f) above. But an even more useful test is to reciprocally cross-check the kinship between pouch young and the mother of their putative cousins. If the two mothers were not full siblings, we expect this pairing to produce an unrelated kin category in the case of 1C offspring. However, in the HAV case, one of the mothers must also be the grandmother of the other pouch young! This would produce the 2nd order (GG) relationship between the pouch young and their grandmother. Thus, pedigree information helps us to distinguish between HAV and 1C pouch young. Once 1C pouch young have been identified (via all approaches) they will constitute an estimate of the elusive intergenerational category PO (sex-independent). Similarly, the HAV offspring where the GG individual is not the parent of the other mother can be used to derive an estimate of male intergenerational dispersal!

For this reason, our key kinship category targets are:

1. first cousins (pouch young)
2. full siblings (pouch young) - a trivial but important category

Pedigree relationships we also need to test include:

1. kinship between parents (to identify FS parents (1C offspring) as well as PO parents (HAV offspring))
2. kinship between pouch young and non-parent females where a 1C/HAV relationship exists between offspring (if 2nd order where parents are not FS, interpret as GG category, meaning offspring are HAV. Otherwise, interpret as 1C)

We run the initial simulations here: (setting the dispersal model to vgamma to allow longer-tailed dispersal)

```
ant_1c_juv <- simulate_kindist_custom(nsims = 100000, model = antechinus_model, kinship = "1C", cycle =
ant_1c_juv
#> KINDISPERSE SIMULATION of KIN PAIRS
#> -----------------------------------
#> simtype:      custom
#> kerneltype:       vgamma
#> kernelshape:      0.5
```

```
#> kinship:      1C
#> simdims:      100 100
#> juvenile              100
#> breeding              50
#> gestation             25
#> pouch             25
#> cycle:            -1 -1
#> lifestage:        pouch
#>
#> tab
#> # A tibble: 100,000 x 8
#>    id1   id2   kinship distance     x1       y1    x2     y2
#>    <chr> <chr> <chr>      <dbl>  <dbl>    <dbl> <dbl>  <dbl>
#>  1 1a    1b    1C          54.2 -31.0      7.97  -26.1   62.0
#>  2 2a    2b    1C         256.  301.      82.3    54.4   16.7
#>  3 3a    3b    1C          34.4  -7.24  113.     -26.8  141.
#>  4 4a    4b    1C         181.   35.2   151.     -35.3  -15.9
#>  5 5a    5b    1C         206.   54.1   -77.9    43.5 -284.
#>  6 6a    6b    1C          67.9 128.     -0.480 193.    20.9
#>  7 7a    7b    1C         177.  159.    159.      41.2   26.5
#>  8 8a    8b    1C         141.   45.9   248.     -69.8  167.
#>  9 9a    9b    1C          98.1 366.    -24.7   296.   -93.6
#> 10 10a   10b   1C         143.  214.   -242.    356.  -224.
#> # ... with 99,990 more rows
#> -----------------------------------
```

```
ant_fs_juv <- simulate_kindist_custom(nsims = 100000, model = antechinus_model, kinship = "FS", cycle =
ant_fs_juv
#> KINDISPERSE SIMULATION of KIN PAIRS
#> -----------------------------------
#> simtype:      custom
#> kerneltype:       vgamma
#> kernelshape:      0.5
#> kinship:      FS
#> simdims:      100 100
#> juvenile              100
#> breeding              50
#> gestation             25
#> pouch             25
#> cycle:            -1 -1
#> lifestage:        pouch
#>
#> tab
#> # A tibble: 100,000 x 8
#>    id1   id2   kinship distance    x1    y1    x2    y2
#>    <chr> <chr> <chr>      <dbl> <dbl> <dbl> <dbl> <dbl>
#>  1 1a    1b    FS             0 72.1   81.3 72.1   81.3
#>  2 2a    2b    FS             0 52.6   58.8 52.6   58.8
#>  3 3a    3b    FS             0 52.2   56.9 52.2   56.9
#>  4 4a    4b    FS             0  9.86  28.0  9.86  28.0
#>  5 5a    5b    FS             0  2.64  55.1  2.64  55.1
#>  6 6a    6b    FS             0 80.1   86.3 80.1   86.3
#>  7 7a    7b    FS             0 81.5   32.4 81.5   32.4
```

```
#>  8 8a    8b    FS             0 30.5   71.1 30.5   71.1
#>  9 9a    9b    FS             0 48.9   41.6 48.9   41.6
#> 10 10a   10b   FS             0 91.2   82.3 91.2   82.3
#> # ... with 99,990 more rows
#> -----------------------------------
```

Inspecting the results, we see that the 1C category is well-dispersed, while the FS category is entirely zero (FS offspring in the pouch are not dispersed at all).

Finally, we run a simple PO estimation with these simulations (we override a key check on the breeding cycle that would otherwise be triggered by FS being used before their breeding cycle start point, as they have not dispersed at all, so will not confound the estimate)

```
axpermute_standard(ant_1c_juv, ant_fs_juv, nsamp = 100, override = TRUE)
#>     2.5%     mean    97.5%
#> 102.3822 117.5524 135.1330
```

This is excellent so far. Mean dispersal is still around 117, so assuming sampling is adequate, this approach will lead us to intergenerational dispersal.

##### 4.4.7 Simulate sampling site and finalise study design

Now, before we go any further, we need to estimate how large a study site we will need to gain an adequate understanding of dispersal, and avoid missing rarer long-tailed dispersal events. We know that our FS pouch young haven't dispersed, so we won't need to worry about them. But what about the 1C category? At this point in an actual study, the existing model should be refined as much as possible to provide 'realistic' estimates of dispersal at each stage (erring on the side of larger estimates if uncertain).

Let's check an initial sampling site of 100m by 100m:

```
ant_1c_juv %>% sample_kindist(dims = 100, n = 1000) %>% axpermute_standard(ant_fs_juv, nsamp = 100, ove
#> Setting central sampling area to 100 by 100
#> Down-sampling to 1000 kin pairs
#> 1000 kin pairs remaining.
#>     2.5%     mean    97.5%
#> 25.47659 27.63455 29.96706
```

A 100x100 metre sampling area is woefully inadequate (estimating the kernel to only ~ 27m, well short of the 117 we need)! We try again, this time in a 1km x 1km site:

```
ant_1c_juv %>% sample_kindist(dims = 1000, n = 1000) %>% axpermute_standard(ant_fs_juv, nsamp = 100, ov
#> Setting central sampling area to 1000 by 1000
#> Down-sampling to 1000 kin pairs
#> 1000 kin pairs remaining.
#>      2.5%      mean     97.5%
#>  85.25080  98.05554 111.11095
```

We are doing better here: with an average of ~100m. But we're still 15% short, and barely including the correct value in our C.I.s What about 2km x 2km?

```
ant_1c_juv %>% sample_kindist(dims = 2000, n = 1000) %>% axpermute_standard(ant_fs_juv, nsamp = 100, ove
#> Setting central sampling area to 2000 by 2000
#> Down-sampling to 1000 kin pairs
#> 1000 kin pairs remaining.
#>      2.5%      mean     97.5%
#>  99.22454 112.75552 127.63553
```

This estimate is acceptable, with a mean only a few metres short, and the true value well within C.I.s. Accordingly, we make the decision to sample within a grid of at least 2km by 2 km.

###### 4.4.8 Run the study!

Now we are as prepared as possible to perform sampling, genotype individuals, etc.

###### 4.4.9 Load data and generate in-field dispersal estimates

Follow the instructions given in 4.2 and 4.3 to load sample data into the program and supply estimates. The `axials_standard` and `axials_permute` functions contain the parameters `acycle` and `bcycle`, which enable the calibration of the estimation process to pouch young (remember to use the `override` parameter in this context). Or you could simply avoid phase information, set the FS category to zero (as they are non-dispersed), and perform a 1C-FS subtraction as is (which will effectively just decompose the 1C into two PO increments - works in this case as at the pouch phase we have synced FS dispersal to PO dispersal (as all FS offspring coincide with maternal parent)).

###### 4.4.10 Cross-check field estimates for bias and calibrate

Once you have generated in-field estimates of dispersal, it is always good practice to substitute these estimate back into the original simulation and rerun the sampling analysis in 4.4.7 again. If the new estimate is significantly underestimated by the simulation after subsampling to the dimensions of your study site, it is likely that the study site is too small, and is biasing estimates of dispersal - one approach from here would be to progressively increase the dispersal distance until the new subsampled estimate matches the one generated by the study (this will be a more likely figure for dispersal in the species, and should inform future studies).
